# Shared subcortical arousal systems across sensory modalities during transient modulation of attention

**DOI:** 10.1101/2024.09.16.613316

**Authors:** Aya Khalaf, Erick Lopez, Jian Li, Andreas Horn, Brian L. Edlow, Hal Blumenfeld

**Author notes:** Correspondence to: Hal Blumenfeld, MD, PhD, Yale Depts. Neurology, Neuroscience, Neurosurgery, 333 Cedar Street, New Haven, CT 06520-8018.

## Abstract

Subcortical arousal systems are known to play a key role in controlling sustained changes in attention and conscious awareness. Recent studies indicate that these systems have a major influence on short-term dynamic modulation of visual attention, but their role across sensory modalities is not fully understood. In this study, we investigated shared subcortical arousal systems across sensory modalities during transient changes in attention using block and event-related fMRI paradigms. We analyzed massive publicly available fMRI datasets collected while 1,561 participants performed visual, auditory, tactile, and taste perception tasks. Our analyses revealed a shared circuit of subcortical arousal systems exhibiting early transient increases in activity in midbrain reticular formation and central thalamus across perceptual modalities, as well as less consistent increases in pons, hypothalamus, basal forebrain, and basal ganglia. Identifying these networks is critical for understanding mechanisms of normal attention and consciousness and may help facilitate subcortical targeting for therapeutic neuromodulation.

## Introduction

Different sensory modalities elicit distinct neural signatures in the brain. However, it can be proposed that there is a fundamental subset of circuits shared across modalities, supporting core functions such as conscious perception and attention control. Subcortical arousal systems are known to play a key role in controlling sustained changes of attention and long-lasting states such as sleep/wake and levels of vigilance^1^. Previous studies on patients with disorders of consciousness confirmed the critical influence of subcortical arousal systems in maintaining states of consciousness^2–4^. However, the role of these subcortical systems in dynamic modulation of attention has been less studied and when examined, the focus has often been on a single sensory modality without considering the shared networks dynamically modulating attention across perceptual modalities^5–7^. Moreover, most research investigating dynamic changes in attention has focused on cortical large-scale networks involved in top-down attentional salience and bottom-up attention control with little emphasis on subcortical systems^8–11^.

Subcortical systems have been increasingly recognized as playing an important role in cognition^12^. Studies on healthy participants and patients with impaired consciousness have demonstrated that the midbrain reticular formation and central thalamus are key subcortical structures that modulate attention^2–4, 13–17^. Additionally, deep brain stimulation studies in humans and animal models demonstrated that stimulation of the central thalamus significantly improves arousal and restores consciousness^18–22^. Previous research suggests that arousal systems in the thalamus, upper brainstem and basal forebrain may contribute to dynamic modulation of attention and conscious perception^5, 6, 23–25^. The dynamic modulation of attention by the subcortical arousal systems is a key mechanism that facilitates conscious perception. Previously, we introduced a data-driven model that describes the sequence of neural mechanisms required to produce conscious awareness of sensory events^26^. The model hypothesizes that one of the mechanisms critical for conscious perception is an attention mechanism that operates through cortical and subcortical arousal systems, mediating stimulus detection, dynamic modulation of arousal, bottom-up attentional salience, and top-down attentional control. In this framework, the subcortical arousal networks provide an early dynamic transient pulse that facilitates subsequent widespread signals necessary for conscious perception^26^. While multiple cortical systems have been implicated in sensory detection, attention and conscious perception^9, 27–32^, the potential key role of subcortical arousal networks in modulating attention and perception across sensory modalities requires further investigation.

Functional magnetic resonance imaging (fMRI) experiments typically use block designs, event-related designs, or a combination of both to identify and characterize both sustained and transient blood-oxygenation level-dependent (BOLD) responses^33–37^. Previous studies have noted cortical BOLD fMRI signal increases at the onset and offset of blocks and events^38–40^, with a few studies investigating BOLD fMRI signal increases in subcortical networks at the onset of blocks and events^6, 7, 15^. Further research is needed to explore the shared subcortical networks facilitating these sustained and transient attention modulations at block onsets and in response to individual event stimuli, respectively.

In this study, we investigate shared subcortical systems during dynamic modulation of attention across sensory modalities with large sample sizes using both block and event-related fMRI designs. Previous studies have highlighted the early transient signals in subcortical arousal systems, suggesting that a model-free fMRI analysis may be more effective at detecting these signals compared to traditional general linear models, which may not adequately capture such early responses^6, 7, 41, 42^. Therefore, we conducted a model-free fMRI analysis for each task included in the study by calculating percentage change in BOLD fMRI signals to identify the subcortical regions activated at task block onsets or in response to individual events, depending on the task design. A conjunction analysis was performed to identify the subcortical regions sharing common early activity across different tasks and sensory modalities. Similarly, we performed a conjunction analysis to identify the shared cortical networks across sensory modalities. Our findings revealed a shared early transient surge in fMRI activity within subcortical arousal systems. Furthermore, we observed similar patterns in the cortical salience and attention networks. These findings provide new insights into brain mechanisms of arousal and attention and may help identify potential therapeutic targets for restoring arousal and consciousness in patients with neurological disorders.

## Methods

### Participants and behavioral tasks

We analyzed 3 Tesla (3T) task fMRI data collected from healthy adults while performing 11 different tasks spanning four sensory modalities: vision, audition, taste, and touch. The data were obtained from six publicly available datasets, providing a large overall sample size (Table 1). The datasets included were the Washington University-University of Minnesota (WU-Minn) Human Connectome Project (HCP) Young Adult ^43, 44^, University of California Los Angeles (UCLA) Consortium for Neuropsychiatric Phenomics^45^, Glasgow University^46^, Jagiellonian University^47^, and two datasets from Yale University^48, 49^. The HCP dataset provided a significant portion of our data; specifically, task-fMRI data from the HCP 1200 Subjects Data Release were used in this study (N= 1113; mean age = 28.8; age range: 22–37 years; females = 606)^50^. In the visual domain, we examined six tasks, including, the gambling^51^, relational processing^52^, working memory^53, 54^, social cognition^55^, and motor^56^ tasks from the HCP dataset as well as the spatial capacity task (SCAP)^45^ from the UCLA Consortium (N=130; mean age ± SD = 31.26 ± 8.74 years; age range = 21-50 years; females = 62). For auditory tasks, we analyzed the language task^57^ from the HCP dataset and the passive listening task ^46^ from the Glasgow University dataset (N = 218; mean age ± SD = 24.1 ± 7.0 years; age range = NA; females =101). As for the taste modality, fMRI data from two tasks^48, 49^ collected at Yale University were incorporated in the analysis (N = 28; mean age ± SD = 27.14 ± 4.75 years; age range = 18–37 years; females = 20, and N=48; mean age ± SD = 27.71 ± 3.94 years; age range = 23-39 years; females = 29). Lastly, in the tactile modality, we analyzed fMRI data of a tactile task^47^ collected at the Jagiellonian University (N=25; mean age ± SD= 25.68 ± 3.3 years; age range = 22-32 years; females = 25). Additional details about the behavioral tasks, fMRI acquisition parameters, and data used in the analysis can be found in the Supplementary Information. The purpose of including different tasks from multiple sensory modalities with large sample sizes was to provide a robust basis for our analyses, allowing identification of shared subcortical systems irrespective of sensory modality, presented stimuli, or task demands.

**Table 1:** Overview of the tasks employed in this study, including key design characteristics and analysis-relevant details.

| Dataset | Task | Stimulus<br>modality | Analysis<br>type | Duration of task,<br>rest blocks (s) | Blocks of<br>task, rest per<br>run | Number of runs,<br>number of blocks or<br>events analyzed per run | Participants |
| --- | --- | --- | --- | --- | --- | --- | --- |
| HCP | Gambling | Visual | Block | 28, 15 | 4, 4 | 2, 3 | 1088 |
| HCP | Relational Processing | Visual | Block | 16, 16 | 6, 3 | 2, 2 | 1045 |
| HCP | Working Memory | Visual | Block | 25, 15 | 8, 4 | 2, 3 | 1091 |
| HCP | Social Cognition | Visual | Block | 23, 15 | 5, 5 | 2, 4 | 1054 |
| HCP | Motor | Visual | Block | 12, 15 | 10, 3 | 2, 2 | 1086 |
| HCP | Language | Auditory | Event | 28, NA | 8, NA | 2, 12-15<sup>a</sup> | 1054 |
| UCLA | Spatial Capacity | Visual | Event | NA, NA | NA, NA | 2, 48 | 130 |
| Glasgow | Passive Listening | Auditory | Block | 8, 12 | 40, 20 | 1, 19 | 217 |
| Yale | Taste Perception I | Taste | Block | 36-72, 15 | 8, 8 | 4, 7 | 28 |
| Yale | Taste Perception II | Taste | Block | 36-72, 10 | 12, 12 | 2, 11 | 48 |
| Jag. Univ. | Reading Braille | Tactile | Block | 4-6, 4-8 | 72, 7 | 8, 0-2<sup>b</sup> | 25 |

<sup>a</sup> Each run in the language task contains 4 story blocks and 4 math blocks. Each story block contained one story and each math block contained between 2-3 math problems yielding 12-15 events per run.
<sup>b</sup> Of the 8 runs in the tactile Braille reading task, rest-to-tactile task transitions occurred in 2 blocks for 5 runs, in 1 block for two runs, and in 0 blocks for 1 run.
\*NA: Not Applicable
Data are from the Human Connectome Project (HCP)<sup>43, 44</sup>, University of California Los Angeles (UCLA) Consortium for Neuropsychiatric Phenomics<sup>45</sup>, Glasgow University<sup>46</sup>, Yale University<sup>48, 49</sup>, and Jagiellonian University (Jag. Univ.)<sup>47</sup>.

Depending on the task-design, we analyzed fMRI data either at task block onset to investigate subcortical and cortical networks modulating transitions from baseline blocks to task blocks (block transitions), or at event onset to investigate networks modulating transitions from baseline to events (event transitions). Event transitions were specifically considered for analysis when the time between consecutive events was jittered, while block transitions were examined when a task block was preceded by a baseline block. These criteria led to the analysis of block transitions in nine tasks and to analysis of event transitions in two tasks (Table 1). Information on the number of baseline blocks, task blocks, events, and the number of blocks/events used in the analysis is reported in Table 1. Note that in some cases the number of task blocks analyzed per run was fewer than the number of task blocks per run. This was because we only analyzed task blocks that were preceded by a baseline block (baseline to task transitions), and for some run designs the first task block had no preceding baseline block, while in others, several task blocks were presented sequentially without baseline blocks separating them. More detailed descriptions of the tasks and fMRI data acquisition parameters are reported in the Supplementary Information.

### Preprocessing and artifact rejection

A standard fMRI data preprocessing pipeline was implemented using the Statistical Parametric Mapping (SPM12) toolbox (http://www.fil.ion.ucl.ac.uk/SPM) in MATLAB (Mathworks, Inc.). This pipeline was applied to all tasks except those from the HCP dataset. The pipeline comprises three main steps, including motion correction, nonlinear spatial normalization to the standard Montreal Neurological Institute (MNI) space, and spatial smoothing with a Gaussian kernel. For motion correction, functional images acquired in each run were spatially realigned to the first image in that run using 3D rigid-body transformation with three translation and three rotation parameters in x, y, and z directions. To transform the motion-corrected functional images to the MNI space, the structural scan for each participant was coregistered to the mean functional image of the motion-corrected functional images within each run. Next, that structural scan was transformed by non-linear warping to MNI space and the corresponding transformation matrix was applied to the motion-corrected functional images. Finally, the normalized functional images were spatially smoothed with an isotropic Gaussian kernel (FWHM = 6mm). Because the tactile task had a long repetition time (TR) of 3 seconds, we applied slice-timing correction in SPM12 before the standard preprocessing steps.

To ensure computational feasibility, we utilized the preprocessed version of the HCP data that had undergone minimal processing using the HCP pipelines^58^. The primary preprocessing steps for the HCP data aligned with the standard pipeline we implemented, with one main difference: the HCP pipeline included additional corrections for susceptibility distortions. These corrections required acquisition of field mapping scans, which were not available for the non-HCP datasets, rendering this step infeasible to replicate. Because the preprocessed HCP data were unsmoothed, we applied the Gaussian smoothing step (FWHM = 6mm) from our standard pipeline to maintain consistency with the non-HCP datasets.

For both HCP and non-HCP tasks, runs were excluded from the analysis if transient head movement exceeded 2 mm of translation and 1° of rotation in any of the three directions. These criteria resulted in excluding 2506 runs out of 13643 runs (18.37%) from the analysis. Information on excluded runs per task are reported in the Supplementary Information (Suppl Table S1). To further remove spatial and temporal noise from the data, the smoothed BOLD functional images were next passed through a five step denoising procedure as in previous work from our group^6, 7, 42^. Data were (1) grey matter masked to exclude non-grey matter voxels using a standard gray matter mask from MarsBaR (http://marsbar.sourceforge.net/) modified to include the midbrain and pons (The mask volume in MNI space will be publicly available upon publication here: https://github.com/BlumenfeldLab/Khalaf-et-al_2024), (2) filtered using a 1/128 Hz high-pass filter, (3) corrected for motion artifacts by utilizing a general linear model with the six rigid-body motion parameters estimated during functional image realignment to regress out motion, (4) subjected to rejection of individual volumes if the volume-to-volume root mean squared difference in BOLD signal (DVARS) at certain time point exceeded a threshold of 5^59, 60^, and (5) subjected to rejection of individual volumes if instantaneous changes in head position, known as framewise displacement (FD) exceeded a threshold of 0.3 at a certain time point^59, 60^. FD is calculated as the sum of the absolute values of change in head movement among the six rigid-body motion parameters.

### Percent Change Analysis

To identify the subcortical and cortical networks showing transient BOLD changes at block and event onset, we performed a model-free fMRI analysis by calculating the percent change in BOLD signal across time for the whole brain as in previous work^6, 7, 61^. Through the percent change analysis, we obtained percent change brain maps and percent change time courses showing the transient BOLD changes associated with block and event transitions. The tasks included in the analysis utilized four different TR values as follows: 0.72 seconds (6 HCP tasks), 1 second (2 Yale tasks), 2 seconds (2 Glasgow and UCLA tasks), and 3 seconds for the tactile task. Percentage change analyses were conducted using the original TRs at which the tasks were acquired, and timing was later adjusted as described below. All analyses were completed in MATLAB using custom functions as well as functions from SPM12.

#### Percent Change Brain Maps

The BOLD percent change was calculated for the time course of each voxel relative to the mean BOLD signal of that voxel across the entire run. The BOLD volume corresponding to the onset of a specific block/event was defined as the volume that immediately preceded the block/event onset. A block/event epoch included all volumes corresponding to the 15 seconds before the block/event onset to the 15 seconds after. Epochs were averaged across blocks/events within the same run then across runs, resulting in a single 30-second-long average percent change epoch for each subject. Temporal resolution of the percent change brain map calculations was retained at the original acquisition TR with the exception of tactile task which was upsampled from TR=3 seconds to TR=2 seconds. Specifically, the upsampling was performed on the subject-level percentage change epochs using linear interpolation. Spatiotemporal cluster-based permutation testing was applied to the averaged epochs across subjects for each task to identify the statistically significant voxels and time points compared to the baseline before the block/event onset (see Statistical Analysis section below).

#### Anatomical Localization and Percent Change Time Courses

To precisely localize the observed subcortical activity on the structural MRI template, we used regions of interest (ROIs) from several published *a priori* anatomical atlases. We used the Harvard ascending arousal network (AAN) atlas^62^ for brainstem nuclei, the Morel atlas^63^ for thalamic nuclei, the basal ganglia human area template (BGHAT) atlas^64^ for basal ganglia, as well as an atlas of basal forebrain and hypothalamus nuclei^65^. For the amygdala, we used the amygdala ROI available through the MNI PD25 atlas^66^. These atlases were used for anatomical localization in the figures, and in Table 2.

**Table 2:**
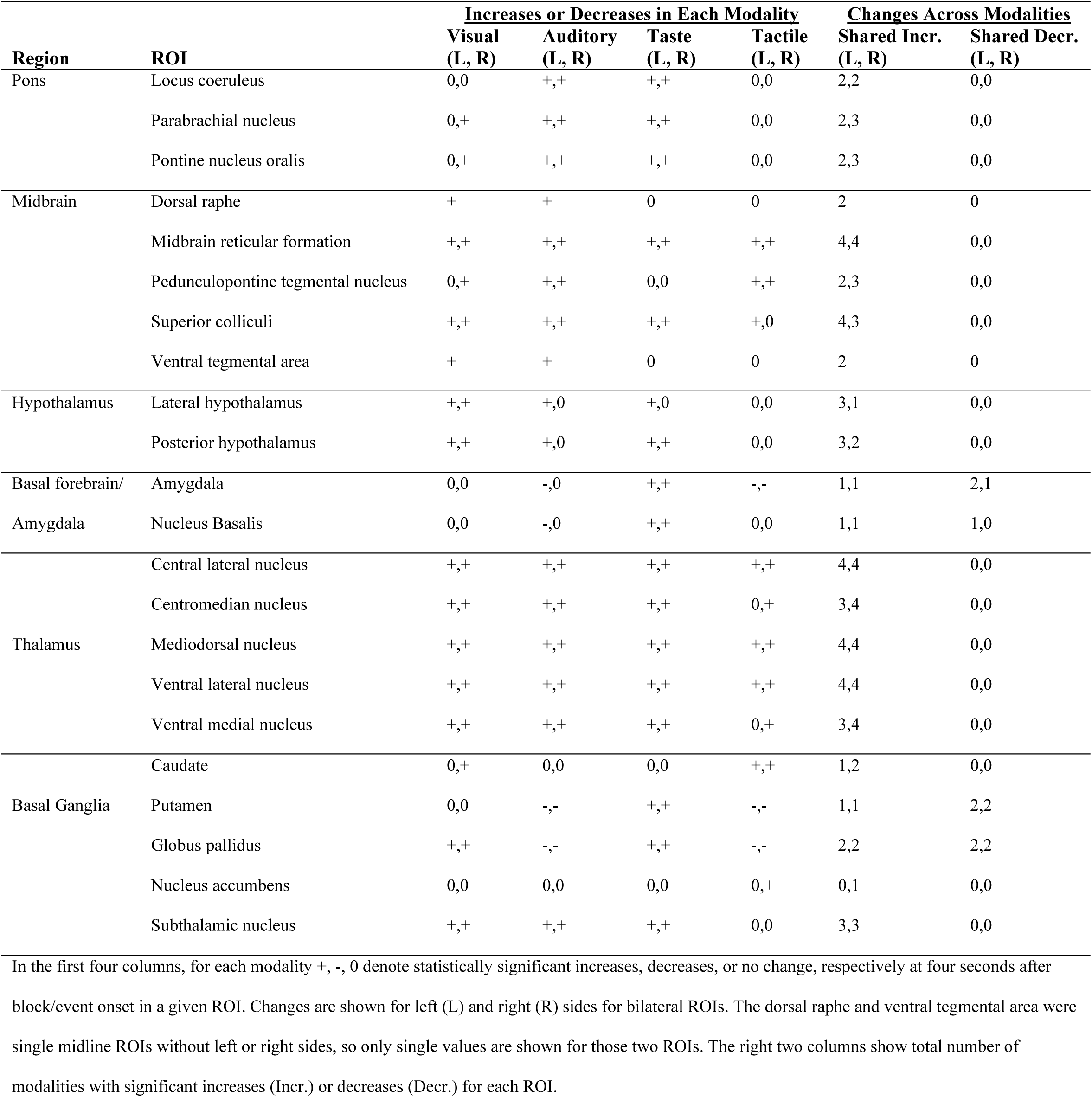
Early transient BOLD fMRI changes in different subcortical ROIs across sensory modalities within four seconds from block/event onset.

In addition to anatomical localization, we used the anatomical atlases to define ROIs for time course analysis in two regions showing shared changes across all modalities, represented by the midbrain reticular formation (AAN atlas) and the thalamic intralaminar central lateral nucleus (Morel atlas). To obtain the time-course of each ROI per subject, we averaged the percentage change time courses across voxels within that ROI using the data from the subject-level percent change maps (see *Percent Change Brain Maps* section). For percent change time course calculations for these two ROIs, subject-level percentage change epochs for all tasks were upsampled through applying linear interpolation to a common TR of 0.72 seconds (HCP sampling rate). To identify the statistically significant time points compared to the baseline before the block/event onset, temporal cluster-based permutation testing was applied to the ROI time courses across subjects as described in the temporal analysis portion of the next section.

### Statistical Analysis

Our overall approach was to first perform separate statistical analyses for each of the 11 tasks in each modality, and to then perform conjunction and disjunction analyses across sensory modalities. Spatiotemporal cluster-based permutation testing was employed to identify voxels and time points showing statistically significant changes in post block/event percent change signals compared to the baseline prior to block/event onset^7^. This approach overcomes the multiple comparisons problem through calculating a single test statistic for the entire spatiotemporal percent change data grid instead of evaluating the statistical significance at each voxel-time point pair^67^. No assumptions are made about the hemodynamic response time course, thus avoiding problems where time course models may not fit the data in some brain regions^41, 42,68^. Additionally, this nonparametric approach does not have assumptions about the distribution of the data which limits false positive rates, especially in high-dimensional data such as fMRI, unlike parametric methods that may incorrectly model functional MRI data, leading to higher false positive rates than their nominal rates^69^. The cluster-based permutation statistical approach implemented in this study was adapted from the Mass Univariate ERP Toolbox ^70^ in MATLAB.

#### Spatiotemporal Analyses

Spatiotemporal statistical analysis was conducted using the original TRs at which the tasks were acquired, except for the tactile task, for which the percentage change data were upsampled to a TR of 2 seconds prior to statistical analysis, as already described. Given the high dimensionality of fMRI data, to improve computational efficiency we implemented two versions of our statistical analysis^7^; a high-resolution version to identify statistically significant changes in subcortical areas, as well as a lower-resolution version to identify statistically significant changes in the whole brain. In the high-resolution subcortical statistical analysis, the spatial resolution of the data was preserved at 2 mm isotropic, but to speed processing the voxels included in the analysis were restricted to the subcortical grey matter voxels in the brainstem, thalamus, basal ganglia, basal forebrain and hypothalamus. In the lower-resolution whole-brain analysis, all the voxels in the grey matter were included, adding the cerebral cortex and cerebellum, but reducing the spatial resolution of the data from 2×2×2 to 6×6×6 mm^3^ to improve computational efficiency. Both the high-resolution subcortical and low-resolution whole-brain statistical analyses were applied to the percent change epochs across subjects in a given task to identify the statistically significant voxels and time points post block/event onset compared to the baseline before the block/event onset. The baseline was defined as the 6 seconds prior to block/event onset.

For the whole brain analysis, spatial resolution was reduced by combining spatially adjacent 2×2×2 mm^3^ grey matter voxels to form larger 6×6×6 mm^3^ voxels. Specifically, the central voxels for each of the 6×6×6 mm^3^ lower-resolution voxels were defined as the original 2×2×2 mm^3^ voxels positioned with exactly 2 intervening voxels until the next central voxel in the x, y, and z directions. Next, all adjacent voxels sharing a face, edge, or vertex with a central voxel were found. These adjacent voxels combined with the central voxel formed the 6x6x6 mm^3^ voxel. Finally, the BOLD percent change signal value within each of the lower spatial resolution 6x6x6 mm^3^ voxels was determined by computing the mean BOLD signal across all 2×2×2 mm^3^ voxels within each of the lower resolution 6×6×6 mm^3^ voxels. If all the adjacent voxels for a certain central voxel were located in the grey matter, the 6×6×6 mm^3^ voxel would include 27 (33) of the 2×2×2 mm^3^ voxels. Otherwise, the larger voxel would combine all available adjacent voxels resulting in a non-cuboidal shaped voxel.

Cluster-based spatiotemporal permutation analysis was performed as in prior work^7^ by generating the spatiotemporal cluster null distribution through 5000 permutation iterations. For each permutation, the mean of the 6-second percent change baseline at a specific voxel and the percent change value of that voxel at the tested time point were randomly shuffled based on the direction of subtraction (time point minus baseline or baseline minus time point) for each participant. Next, a paired, two-tailed t-test compared the permuted values across participants to identify the statistically significant voxels at each tested time point (p < 0.05) from −15 seconds before block/event onset up to 15 seconds afterwards.) Statistically significant spatiotemporal clusters were formed by considering spatial and temporal adjacencies. Negative and positive clusters were created independently. Spatially adjacent voxels were defined as statistically significant voxels (in the same direction) sharing a face, edge, or vertex. Temporal adjacency was found if a voxel was statistically significant (in the same direction) at two or more sequential time points. For each spatiotemporal cluster, the summed absolute value of t-values was computed across all voxels and time points belonging to that cluster. The largest negative and positive cluster determined separately by summed absolute value of t-values was selected from each permutation. Because the positive and negative values were randomly shuffled, we assumed symmetry in the permutation distribution, so we only retained negative clusters and created a one-sided distribution to reduce computations. Therefore, the p-value threshold was set at 0.025 (equivalent to 0.05 in a two-sided distribution). For each permutation, we retained only the negative cluster with the largest absolute t-value and collected these values across 5000 permutations to create a permutation distribution. After generating the spatiotemporal cluster null distribution, the spatiotemporal cluster forming analysis described above was applied to the unpermuted data. Positive and negative clusters were identified separately, and summed t-values with absolute value above the top 2.5% of the permutation distribution were considered significant.

The cluster-based spatiotemporal analysis was performed separately on the whole brain at 6×6×6 mm^3^ resolution, and on subcortical regions at 2×2×2 mm^3^ resolution. Importantly, the high resolution 2×2×2 mm^3^ analysis improved the spatial identification of small subcortical regions, but did not add any new regions to the final conjunction analysis results that were not seen in the whole brain lower resolution analysis. Therefore, for display purposes when showing results of whole brain 6×6×6 mm^3^ resolution analysis on cortical brain slices, we superimposed the 2×2×2 mm^3^ resolution results for subcortical structures on the same slices/surfaces (e.g. Figures 3, 5 and Supplementary Presentations S1 – S3).

#### Temporal Analyses

We implemented a temporal cluster-based permutation test, which is an adapted version of the spatiotemporal cluster-based permutation test described above to identify the statistically significant changes in the ROI percent change time courses^7^. In particular, the cluster-forming approach in the temporal analysis considered only temporal adjacency unlike the spatiotemporal version, which considers both spatial and temporal adjacencies to form spatiotemporal clusters. For each ROI, the temporal cluster-based permutation test was applied to the percent change time courses across subjects for a given task to identify the statistically significant time points post block/event onset compared to the baseline before the block/event. The baseline was defined as the 6 seconds prior to block/event onset. As was already mentioned, before applying temporal statistical analysis, ROI percentage change data for all tasks were resampled to a common TR of 0.72 seconds.

### Subcortical and Whole-brain Conjunction and Disjunction Analyses

#### Binary Conjunction Analysis

To identify the shared subcortical and cortical networks across tasks and sensory modalities, we performed a binary conjunction analysis at each of the time points within an epoch (15 seconds pre and post block/event onset) across tasks. We refer to this as binary conjunction because a voxel was either included or not in the results based on all-or-none statistical criteria. For a voxel to be included in this conjunction, it had to show statistically significant changes (based on permutation testing) in the same direction (i.e., positive or negative) across all modalities and across all 11 tasks at the same time point. Voxels with both positive and negative changes at a given time point were not included in the binary conjunction brain maps. The binary conjunction analysis was performed separately for the high-resolution subcortical and lower-resolution whole-brain statistical results from the permutation testing (Figure 1A, B; Figure 3A; Supplementary Presentation S1).

**Figure 1.**
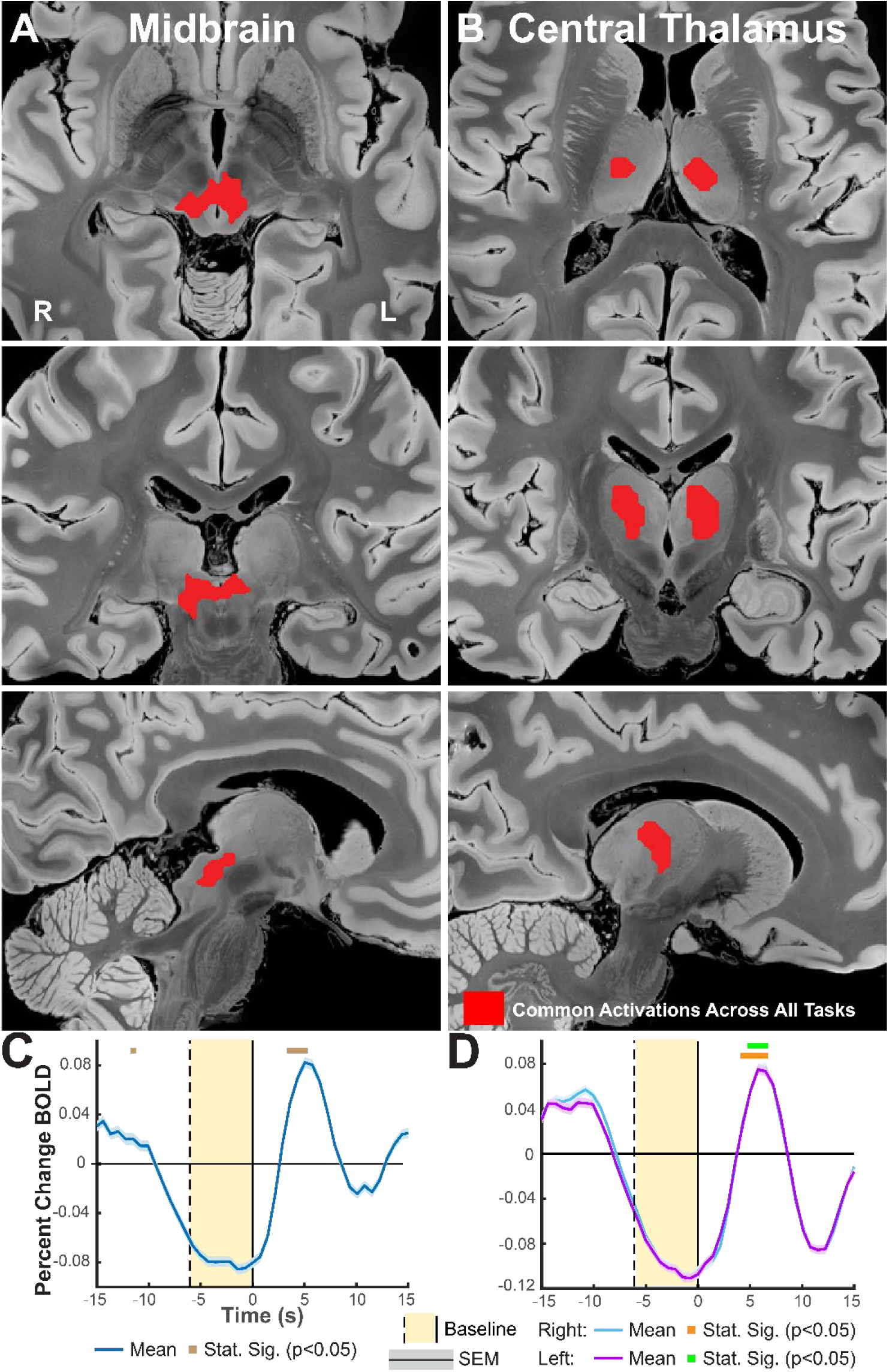
Midbrain and central thalamus show shared subcortical early activations (increases), observed in 11 tasks across four sensory modalities, including, vision, audition, taste, and touch. These shared activations reached statistical significance within four seconds from block/event onset. Cluster-based permutation testing (p < 0.05) was employed to identify the statistically significant changes in percentage change BOLD brain maps and time courses with respect to the baseline before block/event onset for each sensory task. Binary conjunction analysis was then applied across all tasks to identify subcortical regions and time points sharing activations/deactivations across tasks and sensory modalities. (A) Axial, coronal, and sagittal MRI slices in the midbrain showing the spatial extent of the observed shared activations 4s after stimulus onset, mainly centered on the midbrain reticular formation (MRF). No shared deactivations were seeen. (B) Axial, coronal, and sagittal MRI slices showing the spatial extent of the observed activations in the thalamus 4s after stimulus onset, centered on the intralaminar central lateral (CL) nucleus. (A, B) for additional brain slices and time points see Supplementary Presentation S1. (C, D) Mean percent change BOLD time courses across the 11 tasks from two anatomical ROIs, MRF (C) and CL (D), obtained from the Harvard Ascending Arousal Network atlas and the Morel atlas, respectively. The significant time points shared across tasks (permutation based statistics followed by conjunction analysis), marked on the top of the time courses, began 4 seconds after block/event onset in both the MRF and thalamic CL. Data are from 11 tasks obtained across a total of 1,561 participants.

To standardize the analysis TR for conjunction it was necessary to ensure the volumes were temporally aligned at each time point. As mentioned earlier, the tasks included in the analysis utilized four different TR values, including, 0.72 seconds (6 HCP tasks), 1 second (2 Yale tasks), 2 seconds (2 Glasgow and UCLA tasks), and 3 seconds for the tactile task, with only the latter (tactile) TR upsampled from 3 to 2 seconds by linear interpolation. To minimize the need for additional upsampling and creation of new data points that were not physically acquired in the scanner, we selected a common TR of 2 seconds for conjunction analysis across tasks. Specifically, permutation based spatiotemporal cluster-based statistical analysis was done at the original TR for each task, except for the tactile task, as already described, then for conjunction analysis, we used the volume closest in time to the 2 second TR time points for any tasks with higher (0.72 or 1 second) sampling rates.

#### Graded Conjunction Analysis

The binary conjunction approach had strict inclusion criteria, meaning a region would only be included in the conjunction if it was significant across all sensory modalities and tasks at the same time points. To identify regions that are statistically significant across most sensory modalities but not all of them, we introduced a graded conjunction method, to identify significant voxels in 1, 2, 3 or all 4 modalities. Graded conjunction analysis was implemented in the following two ways: 1. On a voxel-by-voxel basis, similar to binary conjunction; 2. In *a prior* defined anatomical ROIs.

For voxel-wise graded conjunction analysis, we began with binary conjunctions within each sensory modality to identify voxels sharing increases or decreases at the same time point, using the same binary approach already described (with the exception of the tactile modality which had only one task, so no within-modality conjunction was needed). This process yielded binary maps for statistically significant increases or decreases across tasks for each sensory modality (Supplementary Presentation S2). Subsequently, these maps were aggregated separately for increases and decreases, with voxel values indicating the number (1 to 4) of sensory modalities sharing statistically significant changes in the same direction (i.e., positive or negative) at each time point (Figure 2; Figure 3B, C; Supplementary Presentation S1).

**Figure 2.**
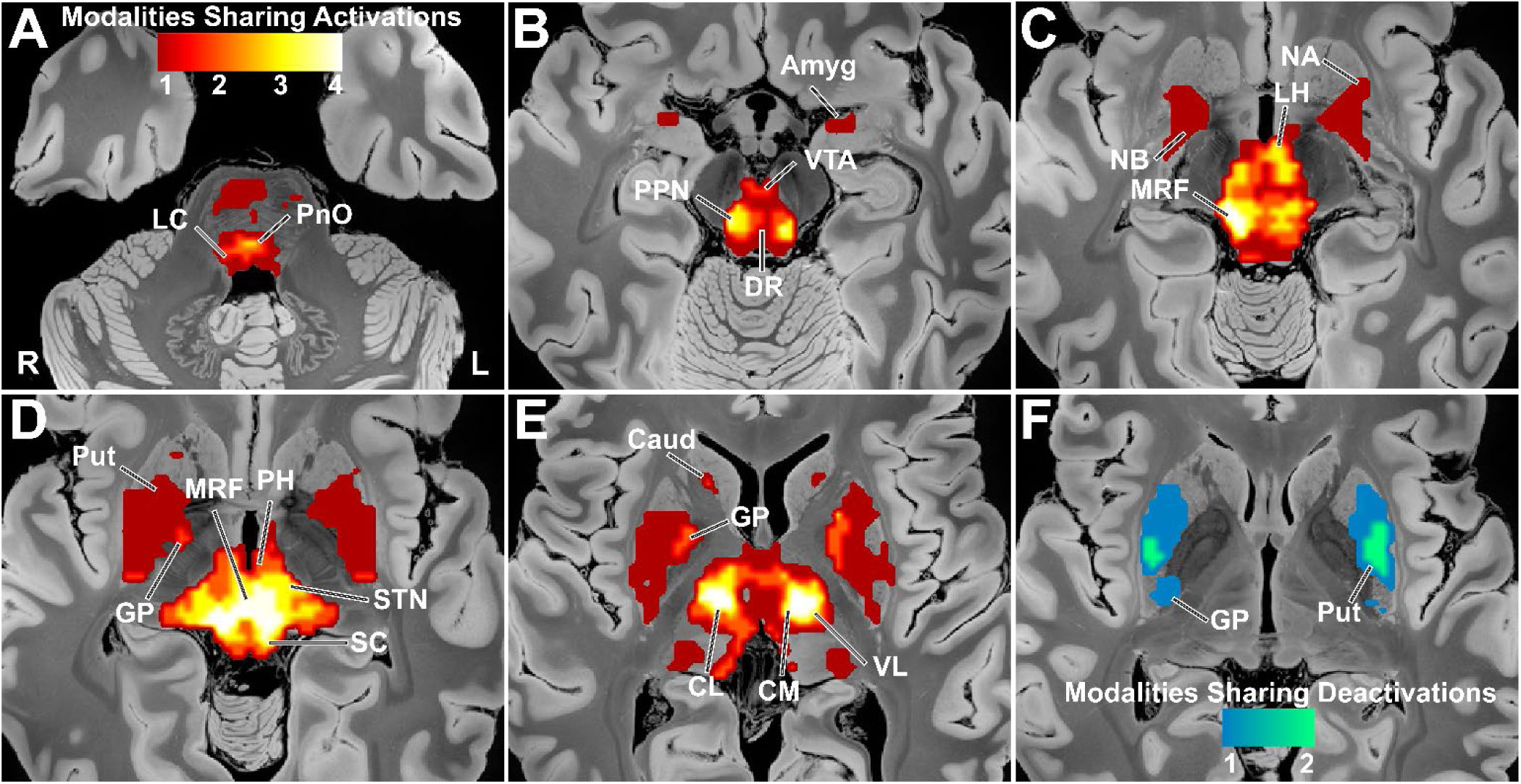
Graded conjunction analysis revealed additional subcortical changes shared less consistently across sensory modalities. This method, which is less stringent than binary conjunction, highlights shared activations (increases) and deactivations (decreases) even if they do not occur across all sensory modalities. The activations and deactivations shown are from four seconds after block/event onset. Spatiotemporal cluster-based permutation testing (p < 0.05) was employed to identify the statistically significant changes in percentage change BOLD brain maps with respect to the baseline before block/event onset for each task, and binary conjunction revealed shared changes within each of the four sensory modalities. Graded conjunction analysis was then applied across the four sensory modalities to identify subcortical regions with shared activations/deactivations, and shared changes were graded from 0 to 4. (A – E) shared subcortical activations; (F) shared subcortical deactivations. For additional brain slices and time points of the graded conjunction analysis please see Supplementary Presentation S1. Locus ceruleus (LC), pontine nucleus oralis (PnO), pedunculopontine tegmental nucleus (PPN), ventral tegmental area (VTA), dorsal raphe (DR), amygdala (Amyg), mibrain reticular formation (MRF), lateral hypothalamus (LH), nucleus basalis (NB), nucleus accumbens (NA), posterior hypothalamus (PH), subthalamic nucleus (STN), superior collicululus (SC), caudate nucleus (Caud), thalamic central lateral nucleus (CL), thalamic centromedian nucleus (CM), thalamic ventrolateral nucleus (VL). Same data and participants as in Figure 1.

**Figure 3.**
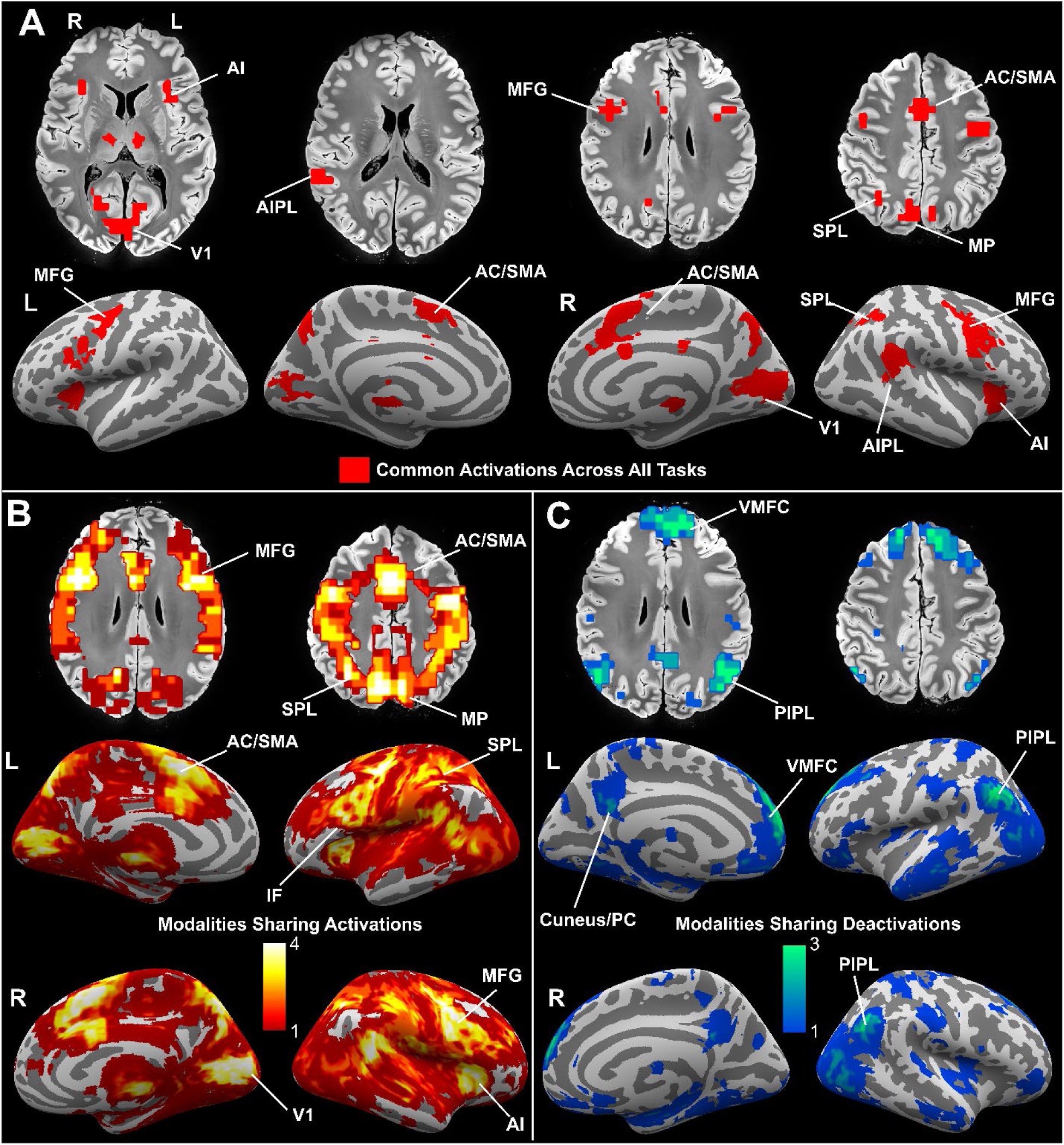
Shared cortical fMRI activations (increases) and deactivations (decreases) four seconds after block/event onset, obtained from whole brain analaysis. (A) Binary conjunction analysis required shared activations/deactivations across the 11 tasks and four sensory modalities. Top row, axial slices; bottom row, surface views. (B) Graded conjunction analysis showing number of modalities sharing cortical activations (significant fMRI increases) across the four sensory modalities. Top row, axial views; bottom rows surface views. (C) Graded conjunction analysis for deactivations across modalities. Top row, axial views; bottom rows surface views. For additional brain slices and time points of the binary and graded conjunction analyses please see Supplementary Presentation S1. Anterior insula (AI), anterior cingulate/supplementary motor area (AC/SMA), primary visual cortex (V1), anterior inferior parietal lobule (AIPL), superior parietal lobule (SPL), medial parietal cortex (MP), middle frontal gyrus (MFG), inferior frontal gyrus/frontal operculum (IF), ventral medial frontal cortex (VMFC), posterior cingulate (PC), and posterior inferior parietal lobule (PIPL). Same data and participants as in Figure 1.

For ROI-based graded conjunction analysis, we used the defined anatomical ROIs based on atlases listed above (see *Anatomical Localization* section). We evaluated each subcortical ROI to determine whether it overlapped with increases or decreases in the binary conjunction maps for each modality (Table 2, left four columns). This was done using a criterion where if more than 50% of the ROI overlapped with significant changes in a given modality, this was counted as an increase or decrease for that ROI. To report the graded conjunction of changes across modalities, we then listed the number of modalities sharing increases or decreases for each ROI (Table 2, right two columns).

#### Disjunction Analyses

We performed exclusive disjunction analyses to identify subcortical and cortical regions unique for each sensory modality. Similar to the conjunction analyses, we performed the disjunction analyses separately on the high-resolution subcortical and lower-resolution whole-brain statistical maps (Figure 4; Figure 5; Supplementary Presentation S3). We first obtained binary conjunction maps across tasks within each sensory modality to identify voxels sharing increases or decreases at the same time point, as already described. We then performed disjunction analysis comparing each modality to the other three. For a voxel to be included in the disjunction of a specific sensory modality, it had to show statistically significant changes in a specific direction (i.e., positive or negative) only in the binary conjunction maps of that sensory modality but not in the binary conjunction maps of any of the other three modalities at the same time point.

**Figure 4.**
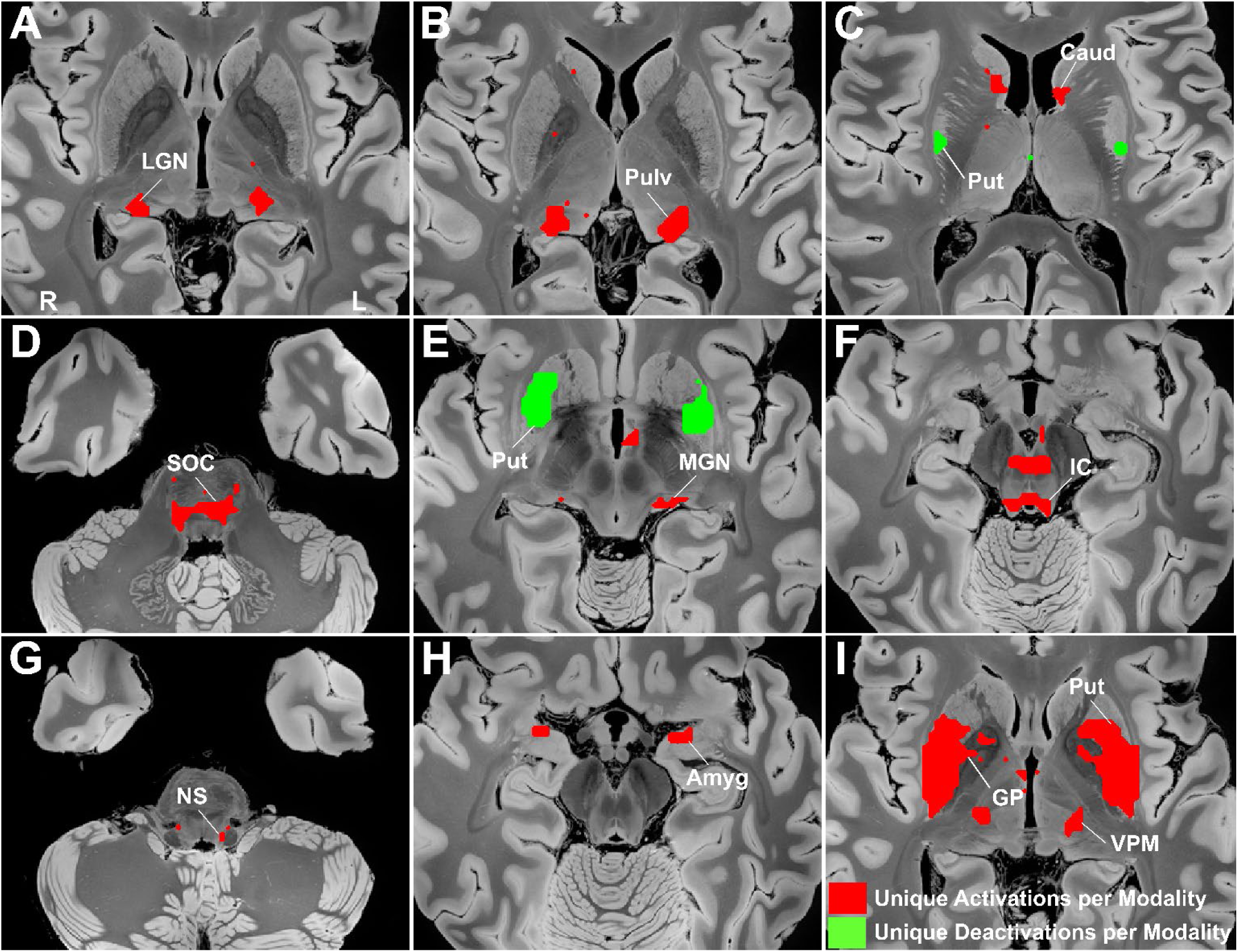
Unique subcortical activations (increases) and deactivations (decreases) for each of the four sensory modalities (vision, audition, taste, and touch) observed four seconds after block/event onset. Exclusive disjunction analysis identified statistically significant subcortical changes present in each modality alone but in none of the other sensory modalities. (A, B) Visual disjunction analysis. (C) Tactile disjunction analysis. (D – F) Auditory disjunction analysis. (G – I) Taste disjunction analysis. For additional brain slices and time points of the disjunction analyses for each modality please see Supplementary Presentation S3. Lateral geniculate nucleus (LGN), pulvinar (Pulv), putamen (Put), caudate nucleus (Caud), superior olivary nuclear complex (SOC), medial geniculate nucleus (MGN), inferior colliculi (IC), nucleus solitarius (NS), amygdala (Amyg), globus pallidus (GP), ventral posterior medial nucleus (VPM). Same data and participants as in Figure 1.

**Figure 5.**
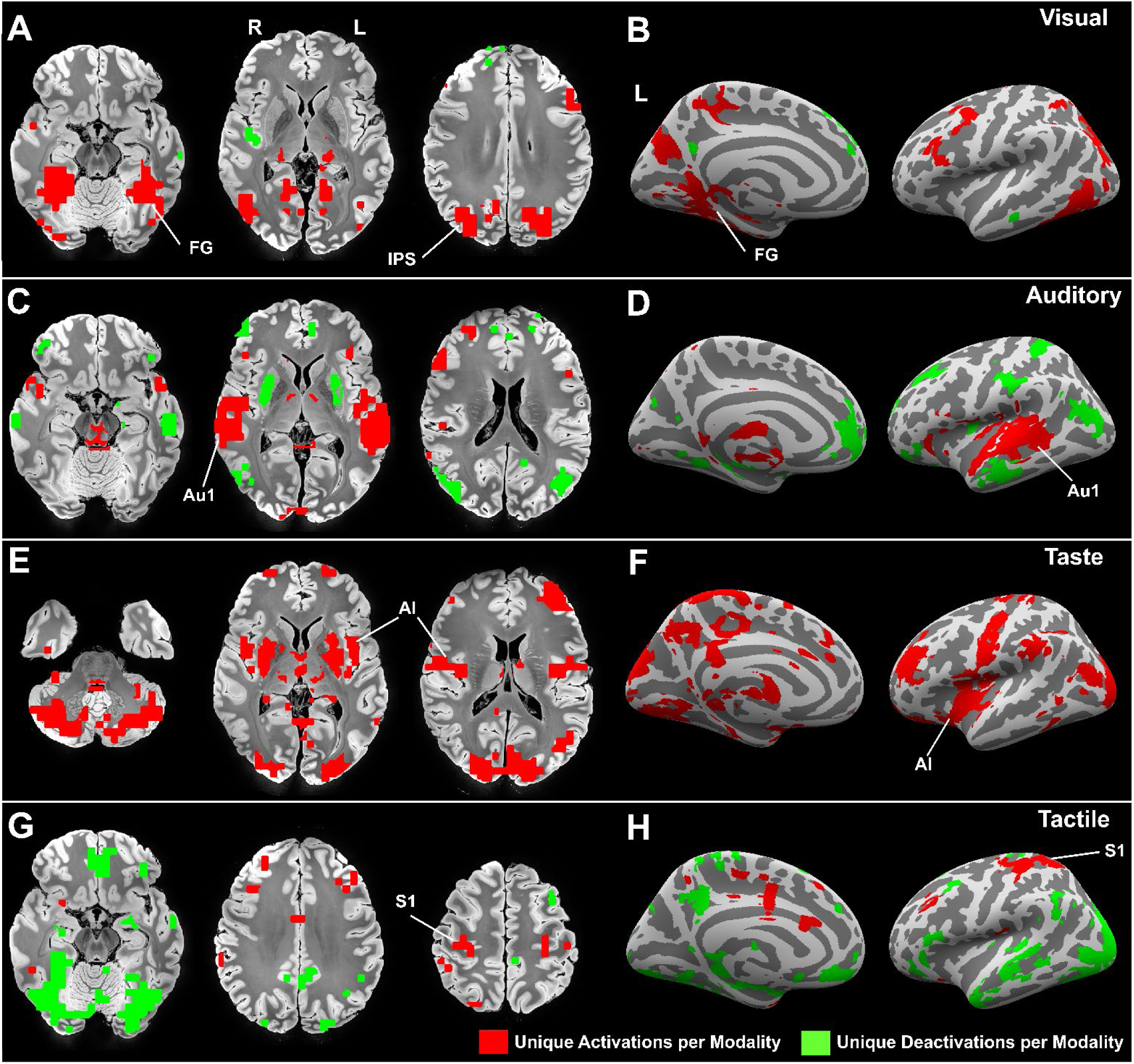
Unique cortical activations (increases) and deactivations (decreases) for each of the four sensory modalities (vision, audition, taste, and touch) observed four seconds after block/event onset. Exclusive disjunction analysis identified statistically significant cortical changes present in each modality alone but in none of the other sensory modalities. (A, B) Visual disjunction analysis. (C, D) Auditory disjunction analysis. (E, F) Taste disjunction analysis. (G, H) Tactile disjunction analysis. (A, C, E, G) Axial brain slices. (B, D, F, H) Left hemisphere surface views. For additional brain slices and time points of the disjunction analyses for each modality please see Supplementary Presentation S3. Fusiform gyrus (FG), intraparietal sulcus (IPS), primary auditory cortex (Au1), anterior insula (AI), primary somatosensory cortex (S1). Same data and participants as in Figure 1.

#### Brain Map Visualization

The conjunction and disjunction maps we obtained were overlaid on a 100 micron 7T MRI structural scan of an ex vivo human brain^71^ for improved visualization and localization of the subcortical structures of interest. These maps were also plotted on the fsaverage FreeSurfer (https://surfer.nmr.mgh.harvard.edu/) inflated brain surface (left hemisphere: lh.inflated surface; right hemisphere: rh.inflated surface) to show the spatial extent of the shared cortical networks as well as the cortical networks unique to each sensory modality.

### Time-Course Conjunction Analysis

We performed a binary conjunction analysis on the ROI time courses to identify the time points sharing common significant changes across all modalities and tasks relative to the baseline before block/event onset for each subcortical ROI (see *Temporal Analyses* section above). Similar to the conjunction analysis applied to brain maps, for a time point to be included in the conjunction, it had to show statistically significant changes in the same direction (i.e., positive, or negative) across all modalities and all 11 tasks at the same time point. The binary conjunction was performed at each time point spanning from −15 to 15 seconds relative to the block/event onset. For display purposes (Figure 1C, D), the mean percent change time course for each ROI was calculated by first averaging across subjects within each task, and these time courses were then averaged across all 11 tasks.

## Results

Previous studies have mainly focused on the role of cortical networks in top-down and bottom-up dynamic modulation of attention^9, 31, 72, 73^. Meanwhile, subcortical arousal structures are mainly known for their role in controlling long-lasting states such as sleep-wake cycles^1^, but their role in dynamic modulation of attention has been increasingly studied recently^5–7^. In the current study, we aim to investigate a shared transient pulse of activity in subcortical arousal systems that occurs with modulation of attention across 11 different tasks spanning four sensory modalities, including, vision, audition, taste, and touch with large sample sizes to ensure robustness of the results and that the observed networks are independent of the task design, type, or demands. This approach allows better isolation of brain activity due to dynamic transitions in attention from the activity due to particular stimuli/tasks. We performed a model-free fMRI analysis by calculating percent change in BOLD fMRI signals with respect to the mean of each fMRI run. To identify the statistically significant changes in percent change BOLD brain maps and time courses with respect to the baseline just prior to transitions in attention, we employed cluster-based permutation testing (p < 0.05). Binary and graded conjunctions were performed on the statistical brain maps and time courses to identify the shared subcortical and cortical regions across sensory modalities. Disjunction analyses were applied to the statistical brain maps to identify unique cortical and subcortical regions for each sensory modality.

Binary conjunction analysis showed a shared transient pulse of subcortical fMRI increases across all sensory modalities and tasks in the midbrain and central thalamus within four seconds from the stimulus onset (Fig. 1.A, 1.B). These increases were centered mainly on the midbrain reticular formation (MRF) and thalamic intralaminar central lateral nucleus (CL) which are key subcortical structures for arousal and attention modulation^1, 6, 23, 25, 62^. Shared fMRI increases across modalities extended into adjacent anatomical regions of the midbrain tegmentum and into other nearby thalamic nuclei such as the mediodorsal nucleus and ventrolateral nucleus bordering thalamic CL (See Supplementary Presentation S1 for binary conjunction maps in additional brain slices and time points). To investigate the timing of these changes, we performed a conjunction analysis of the mean time course of percent change fMRI signals in the MRF and thalamic CL nucleus across all tasks (Fig 1.C, D). This demonstrated a shared significant transient increase in both regions across all sensory modalities and tasks within four seconds from the stimulus onset, which remained significant for an additional 2-4 seconds before returning towards baseline. Thus, a transient pulse of fMRI activation was seen most consistently in the midbrain reticular formation and central thalamus during transitions of attention in a large data set across perceptual modalities and tasks.

Our binary conjunction approach employed stringent inclusion criteria where a region could be included in the conjunction only if it showed significance across all sensory modalities at the same time point. To pinpoint regions statistically significant across some sensory modalities, but not necessarily all of them, we conducted graded conjunction analyses. The graded conjunction analysis revealed subcortical increases and decreases less consistently shared across modalities, not detectable through the strict binary conjunction that required the activity to be shared across all tasks. Through the graded conjunction analysis, we found early fMRI changes after stimulus onset overlapping several subcortical structures, including those in the pons, midbrain, hypothalamus, basal forebrain, amygdala, thalamus and basal ganglia (Fig. 2 and Table 2; see also Supplementary Presentation S1 for graded conjunction maps in additional brain slices and time points). In the pons, shared increases were noted in at least two sensory modalities 4 s after stimulus onset in the locus coeruleus, parabrachial nucleus, and pontine nucleus oralis. In the midbrain, in addition to the MRF, early shared increases were observed in the dorsal raphe, pedunculopontine tegmental nucleus, superior colliculi, and ventral tegmental area. In the thalamus, in addition to CL, consistent increases were seen in all modalities in adjacent central thalamic regions of the mediodorsal and ventrolateral nuclei. The nearby centromedian and ventral medial nuclei also showed increases in three or four modalities. Increases in at least two modalities were also seen in the lateral and posterior hypothalamus, as well as in the basal ganglia caudate, globus pallidus and subthalamic nucleus. Increases in only one modality were seen in the amygdala, nucleus basalis, nucleus accumbens and putamen. fMRI decreases were less consistently seen in subcortical structures at early times, with shared decreases seen across two sensory modalities in the amygdala, putamen and globus pallidus; and in one modality in the nucleus basalis.

To comprehensively delineate the brain networks outside subcortical regions that participate during transitions in attention across sensory modalities, we performed a whole-brain binary conjunction analysis to identify the involved cortical networks. The whole-brain conjunction showed transient cortical increases at early times in detection, arousal and salience networks, including bilateral visual cortex, bilateral anterior insula and bilateral anterior cingulate/supplementary motor area (Fig. 3A). Early increases were also observed in attention and executive control networks, including the right anterior inferior parietal lobule, right superior parietal lobule, bilateral medial parietal cortex, and bilateral middle frontal gyrus (Fig. 3A). For cortical regions, we also conducted a graded conjunction analysis to identify fMRI changes present in some but not all sensory modalities. The graded conjunction analysis enabled identification of additional bilateral cortical regions showing less consistent increases across modalities at early times including the opercular part of the inferior frontal gyrus (Fig. 3B). In addition, although no early shared cortical fMRI decreases were observed across all sensory modalities, the graded conjunction analysis revealed early decreases in at least three modalities in default mode network areas, including the ventral medial prefrontal cortex, posterior cingulate/precuneus, and posterior inferior parietal lobule (Fig. 3C; see also Supplementary Presentation S1 for binary and graded conjunction maps of shared cortical changes in additional brain slices and time points).

To further validate our approach investigating shared changes across sensory modalities, we also analyzed changes specific to each modality. As already described for the binary and graded conjunctions analyses, we began by constructing binary conjunction maps across tasks within each modality to obtain changes for the four modalities (see Supplementary Presentation S2). We then used exclusive disjunction analyses to identify changes unique for each sensory modality. This approach retained only voxels that showed statistically significant increases or decreases for one modality but no others at each location in the brain. We found expected sensory modality-specific changes at early times after stimulus onset in both subcortical and cortical regions. Thus, the subcortical disjunction analysis revealed fMRI increases in the lateral geniculate nucleus and pulvinar exclusively for visual tasks (Fig. 4A, B); increases in the superior olivary complex, medial geniculate nucleus, and inferior colliculus as well as decreases in the putamen exclusively for auditory tasks (Fig. 4D - F); increases in the nucleus solitarius, ventral posterior medial nucleus, amygdala and regions of the basal ganglia exclusively for the taste tasks (Fig. 4 G – I); and increases in the caudate nucleus as well as decreases in a portion of the putamen for tactile tasks (Fig. 4C). Cortical disjunction analyses likewise showed mainly expected changes unique to each sensory modality at early times after stimulus onset. These included increases in the fusiform gyrus and intraparietal sulcus for visual tasks (Figure 5A, B); increases in primary auditory cortex for auditory tasks (Fig. 5C, D); increases in the anterior insula and other regions for taste tasks (Fig. 5 E, F); and increases in primary somatosensory cortex along with changes in several other cortical regions for the tactile tasks (Fig. 5G, H; see also Supplementary Presentation S3 for disjunction maps for each sensory modality in additional brain slices and time points).

## Discussion

We identified a shared subcortical arousal network across four sensory modalities – vision, audition, taste, and touch. The regions belonging to this network showed an early transient pulse of fMRI increases across 11 tasks within four seconds from the onset of task blocks and individual events. These increases were centered mainly on the MRF and thalamic intralaminar CL, structures pivotal for arousal and attention modulation. The time courses of percent change BOLD signals in the MRF and CL demonstrated a shared significant transient increase within four seconds from the stimulus onset. Besides CL, other nearby central thalamic nuclei overlapped with the observed increases, including, mediodorsal, ventrolateral, centromedian, and ventral medial nuclei. In addition to the identified subcortical network, a shared cortical network was activated at the same time frame in regions important for signal detection, attentional salience and top-down control such as the visual cortex, anterior insula, anterior cingulate/supplementary motor area, anterior inferior parietal lobule, superior parietal lobule, medial parietal cortex, and middle frontal gyrus. At the same time frame (four seconds from stimulus onset), less consistent increases and decreases were observed in multiple arousal and/or attention-related subcortical areas in the pons, midbrain, hypothalamus, basal forebrain, basal ganglia, and amygdala. Cortically, less consistent increases were observed in several regions such as the opercular part of the inferior frontal gyrus associated with attention control, and decreases were observed in the default mode network. Collectively, these observations provide new insights into brain mechanisms of arousal and attention irrespective of sensory modality, presented stimuli, or task demands and could lead to improved targeted therapies for disorders of arousal, attention and consciousness.

Several models of attention suggested a potential role of subcortical networks in attention modulation^74–77^, however, previous studies have mainly focused on the role of cortical large-scale networks in top-down and bottom-up attention regulation^9, 31, 72, 73^. Meanwhile, subcortical arousal networks have been mainly investigated for their involvement in controlling sustained changes of attention and state such as sleep-wake cycles^1, 62, 78^. Recently, the role of these subcortical networks in dynamic modulation of attention has been increasingly recognized. Previous studies suggested that arousal systems in the thalamus, upper brainstem and basal forebrain may contribute to dynamic modulation of attention and conscious perception^5, 6, 23–25^. This is further supported by lesion studies in the brainstem and thalamus identifying a key role of these regions in conscious perception and attention modulation ^79, 80^. Although findings of several recent studies highlighted the involvement of some subcortical structures in dynamic attention control^7, 81^, the potential role of subcortical networks in modulating attention across sensory modalities has not been investigated.

An early bilateral pulse of increases was observed within the midbrain and central thalamus within four seconds from the stimulus onset. Notably, this observation is very early given the relatively low temporal resolution of fMRI, but represents the earliest time at which the rising phase of these increases reach statistical significance, whereas the peak occurs 1-2 seconds later. The midbrain and central thalamic increases were common across the four sensory modalities, including vision, audition, taste, and touch. This may reflect the common role of these regions in attention modulation irrespective of the sensory modality or the specific tasks/stimuli presented to the participants. Previous studies on healthy participants and patients with impaired consciousness suggested that the MRF and central thalamus are key subcortical structures in the modulation of attention^2–4, 13, 14, 16, 17^. Additionally, deep brain stimulation studies in human and animal models showed that stimulation of the central thalamus significantly improves arousal and restores consciouness^18, 19, 21, 22, 82, 83^. The early bilateral pulse of increases we identified within the MRF and central thalamus aligns with the findings from previous intracranial EEG and fMRI studies that investigated the role of these regions in conscious perception and dynamic modulation of attention across visual tasks requiring varying degrees of attention^6, 7^. Furthermore, our findings are consistent with a seminal positron emission tomography study that reported early cerebral blood flow increases in MRF and intralaminar thalamus while participants performed an attention-demanding reaction-time task ^84^.

Several neurotransmitters play an important role in attention and/or arousal modulation, including acetylcholine, glutamate, dopamine, noradrenaline, histamine, and orexin ^85–87^ ^88^. The current study identified various subcortical regions associated with attention modulation, each predominantly utilizing one or more of these neurotransmitters. Among the identified regions, pontine nucleus oralis, midbrain reticular formation, and central thalamus, primarily employ glutamate for attention control^2, 87, 89–92^. Other subcortical structures that we visualized with some involvement at early times, such as the parabrachial complex, pedunculopontine tegmental nucleus, and nucleus basalis utilize primarily acetylcholine, along with glutamate and GABA^87, 89, 93^. Meanwhile, the locus coeruleus, ventral tegmental area, and dorsal raphe use primarily noradrenaline^94, 95^, dopamine^96^, and serotonin^93, 97^, respectively, although each contain other neurotransmitters as well. Furthermore, the posterior hypothalamus including the tuberomammillary nucleus and the lateral hypothalamus are recognized for their roles in releasing histamine and orexin, respectively, to modulate arousal ^78, 88, 98, 99^. Additional subcortical structures showed significant changes with respect to baseline across some sensory modalities including the superior colliculi, caudate, putamen, globus pallidus, nucleus accumbens, and amygdala. Previous studies indicated that the amygdala plays key roles in attention, arousal, and decision making^100^. The superior colliculus is mainly known for its role in stimulus detection and modulation of spatial attention^28, 101–104^. Basal ganglia structures including the caudate, putamen, globus pallidus, and nucleus accumbens were found to control sleep-wake transitions ^105^ and play a role in recovery of consciousness after a brain injury^106, 107^. Although we identified BOLD decreases in basal ganglia for some modalities, these decreases do not necessarily reflect decreases in neural activity^108, 109^.

We identified a shared cortical network that includes regions involved in event detection, bottom-up attentional salience, top-down attentional control, conscious perception, and motor preparation^6–11, 110, 111^. The identified regions included anterior insula and anterior cingulate/supplementary motor area which are key structures in the salience network as well as additional regions belonging to the attention and executive control networks, including regions of the parietal lobe and lateral frontal cortex^6–11^. Interestingly, consistent early cortical increases across modalities also included the primary visual cortex, which may speak to the potential cross-modal function of some primary cortical regions in sensory processing^112^. Less consistent increases were observed in the opercular part of the inferior frontal gyrus, known to play a role in attention control^113–115^, and less consistent decreases were observed in well-known default mode areas, including ventromedial frontal cortex, precuneus, and the posterior inferior parietal lobule^6, 7, 116–119^.

Our findings support a data-driven hypothesis we introduced previously to describe the sequence of neural mechanisms required to produce conscious perceptual awareness of external sensory stimuli^26^. In particular, the transient pulse of activation in subcortical arousal systems observed across sensory modalities in the current study fits in this framework. We hypothesize that for a sensory stimulus to be consciously perceived, it has to be first detected by the primary cortex and other cortical and subcortical signal detection circuits. Next, a dynamic transient pulse of activity in subcortical and cortical arousal systems modulates attention and facilitates subsequent widespread signal processing necessary for conscious perception. Then, potentially competing activity in the default mode network is switched off. Finally, a broad wave of hierarchical processing progresses through association cortical areas to fully process the event before it is encoded in memory systems. Our present findings strengthen this hypothesis^26^ by identifying a highly consistent transient pulse of increased fMRI activity in midbrain and central thalamus shared across visual, tactile, auditory and taste stimuli, associated with transitions of attention in tasks requiring sensory perception.

The disjunction analysis helped to validate our approach by showing cortical and subcortical regions that are well-known to be associated with each sensory modality. For instance, visual tasks showed unique activations in lateral geniculate nucleus, pulvinar, fusiform gyrus and the intraparietal sulcus^120, 121^, while auditory tasks showed unique activations in superior olivary nuclear complex, inferior colliculus, medial geniculate nucleus, and primary auditory cortex^122^. Unique activations for taste included the nucleus solitarius, ventral posterior medial nucleus, amygdala, and anterior insular cortex^123, 124^. Additionally, unique increases were observed for touch in the caudate nucleus and primary somatosensory cortex^125, 126^. Unique decreases in different parts of the putamen were found in audition and touch. Previous studies have shown that the putamen is involved in attentive processing of auditory or tactile stimuli, but with increased BOLD activity^127–129^. Thus, the decreases observed in the current study need to be further investigated^109^.

Our study has several limitations that should be addressed in future work. Techniques have been proposed to improve inter-subject subcortical co-registration, but are so far not widely used ^130, 131^. These approaches typically require high computational costs, rendering their application impractical in our present study due to the substantial sample size, which exceeded 1,500 participants. Because we did not use such approaches in the current study, we were cautious to avoid making strong conclusions on the voxel level, particularly if the activations/deactivations were not centered on anatomically known structures. Although we included large sample sizes to identify the shared subcortical and cortical networks, the analyzed datasets were not balanced across sensory modalities due to the limited availability of tasks from certain sensory modalities such as taste and touch. No olfaction tasks suiting our analysis purposes were available. Future studies should aim to balance the sample sizes across sensory modalities, and should include more taste and tactile tasks if available. Inclusion of olfaction tasks is an important future direction, particularly because some olfactory signaling pathways bypass the thalamus. This will help to further identify the shared changes across all senses. Although fMRI provides comprehensive anatomical mapping of cortical and subcortical structures not available with more spatially limited human electrophysiological methods, it has lower temporal resolution, and therefore may provide limited information about the sequence of activations/deactivations within the observed networks. Further investigation of these networks could be performed in animal models through direct electrophysiological recordings, or in human studies with availability of subcortical depth electrodes to identify the temporal dynamics of these networks. In addition, regions that are common to some but not all sensory modalities need to be investigated further to identify why they are specific to certain sensory modalities but not others.

In summary, although previous work in conscious perception and attention modulation has recognized the regions we found, prior studies were conducted predominantly in individual sensory modalities. Our approach of analyzing different tasks spanning multiple sensory modalities and with overall large sample size, enabled us to identify changes independent of the task design, demands or stimulus type. We found the most consistent subcortical change associated with transitions in attention was a transient increase in activity in the MRF and central thalamus. These subcortical changes were accompanied by consistent increases in activity in cortical detection, arousal and salience networks, as well as by less consistent changes in multiple other subcortical and cortical regions. Further investigation of the shared subcortical arousal systems participating across sensory modalities could lead to improved targeted therapies for disorders of arousal, attention and consciousness^82, 132–134^ and a better understanding of the complex spatiotemporal mechanisms of normal brain function.

## Supporting information

Supplementary Presentation S1

Supplementary Presentation S2

Supplementary Presentation S3

Supplementary Data Guide

## Acknowledgements

This work was supported by NIH R01 NS134655 (to H.B.), the Mark Loughridge and Michele Williams Foundation, and the Betsy and Jonathan Blattmachr family. A.H. was supported by the German Research Foundation (Deutsche Forschungsgemeinschaft, 424778381 – TRR 295), Deutsches Zentrum für Luft- und Raumfahrt (DynaSti grant within the EU Joint Programme Neurodegenerative Disease Research, JPND), the National Institutes of Health (R01MH130666, 1R01NS127892-01, 2R01 MH113929 & UM1NS132358) as well as the New Venture Fund (FFOR Seed Grant). A.H. reports lecture fees for Boston Scientific and is a consultant for FxNeuromodulation and Abbott.

## Data Availability

Datasets included in the study are publicly available through OpenNeuro and Human Connectome Project websites.

## Code Availability

Analysis codes for this study will be publicly available upon publication here: https://github.com/BlumenfeldLab/Khalaf-et-al_2024

