## Supplementary Presentation S1 for "Shared subcortical arousal systems across sensory modalities during transient modulation of attention"

### Binary and Graded Conjunction Across Modalities

### Binary Conjunction Across All Modalities

### Binary Conjunction Across All Modalities

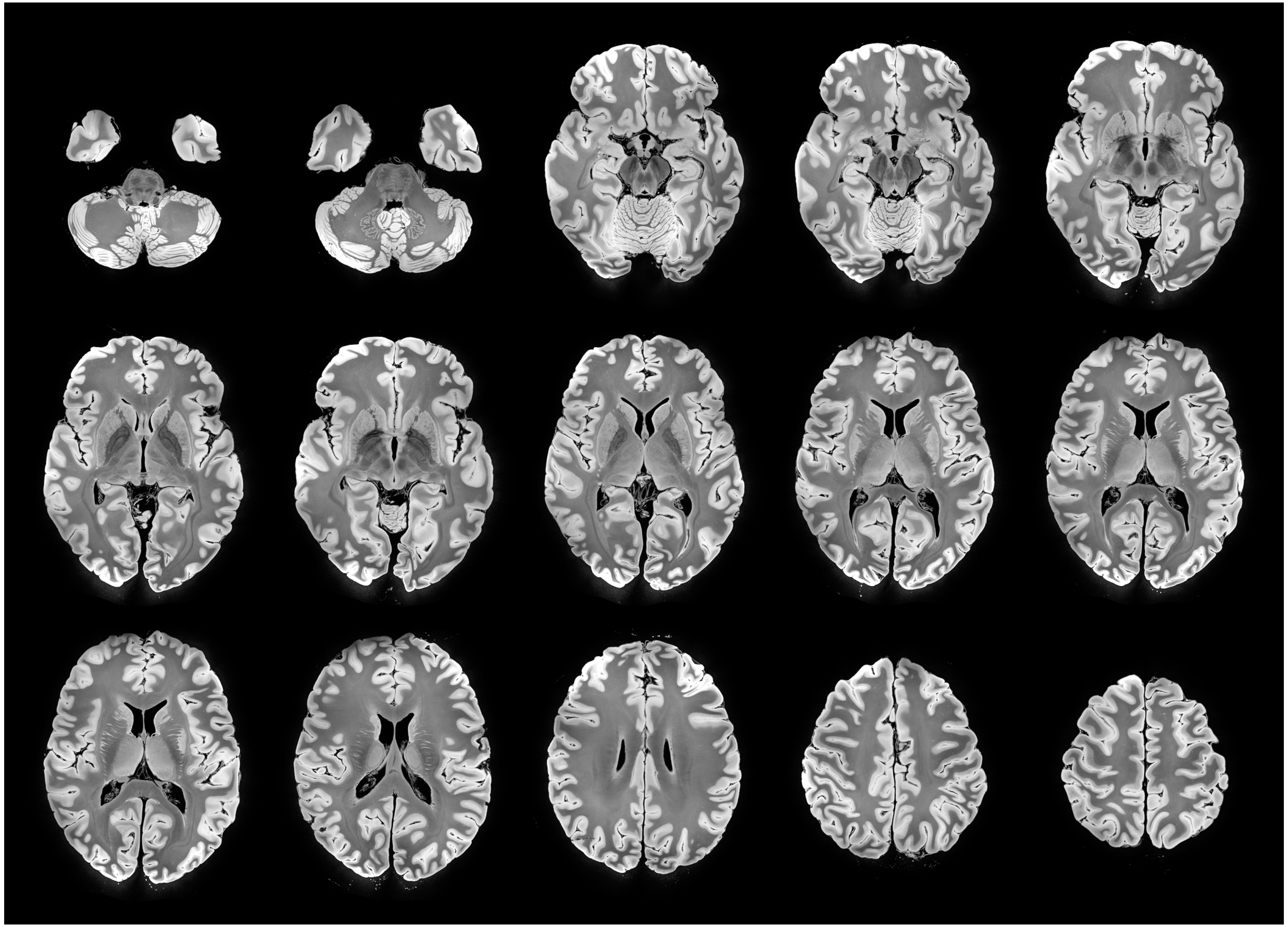

Common activation across all tasks

Common deactivation across all tasks

-4 sec

### Binary Conjunction Across All Modalities

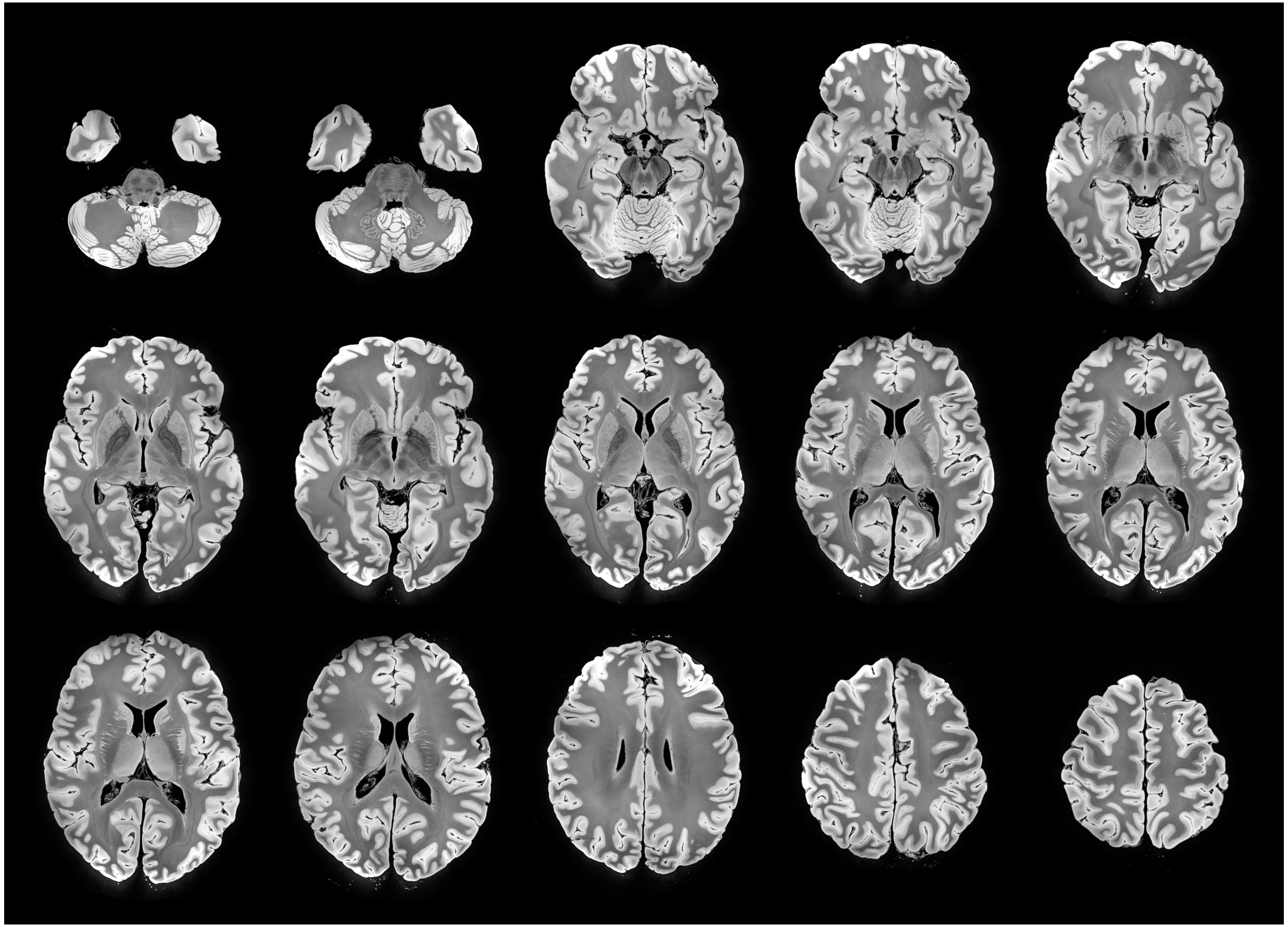

Common activation across all tasks    Common deactivation across all tasks

-2 sec

### Binary Conjunction Across All Modalities

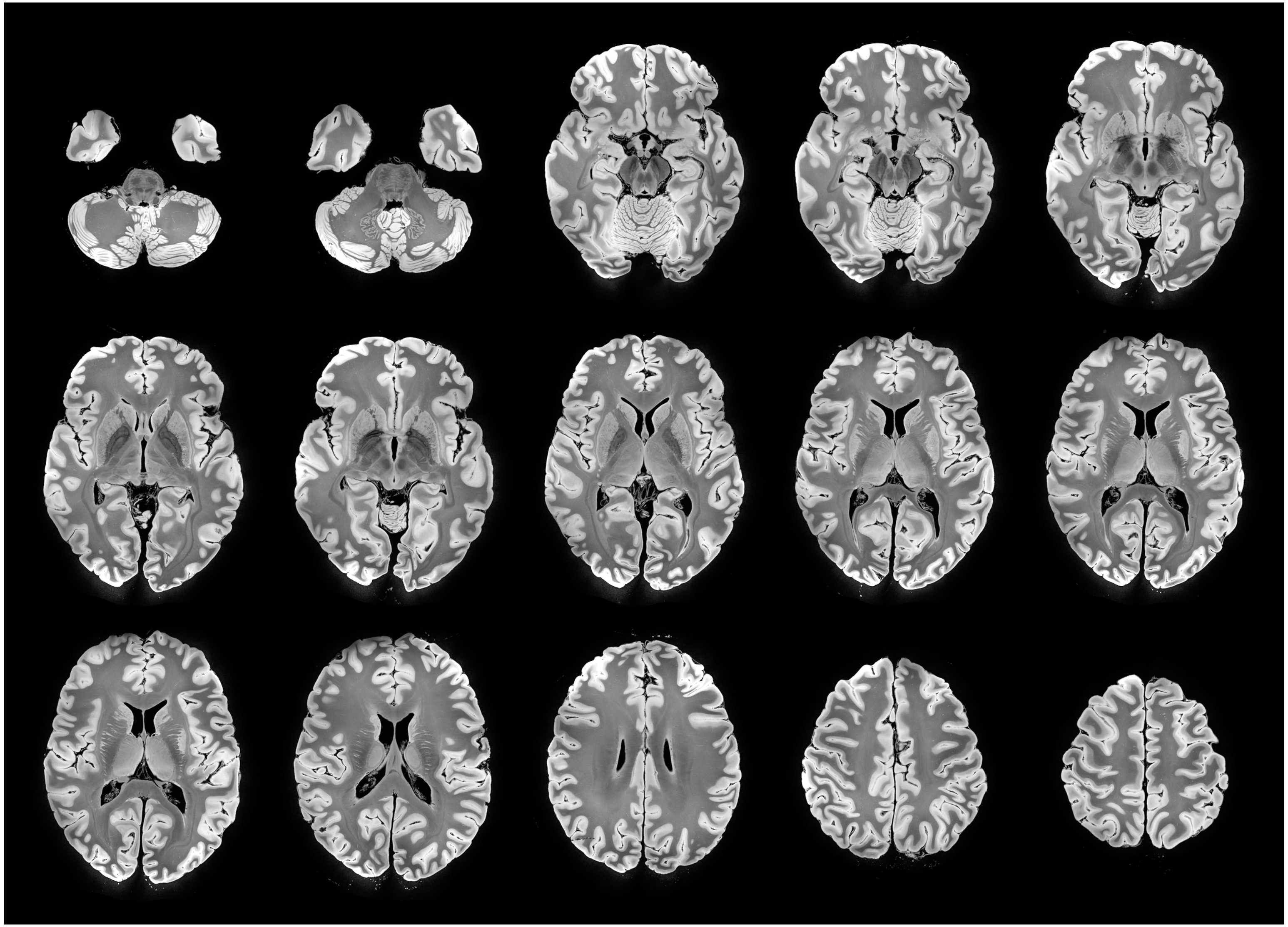

Common activation across all tasks

Common deactivation across all tasks

0 sec

### Binary Conjunction Across All Modalities

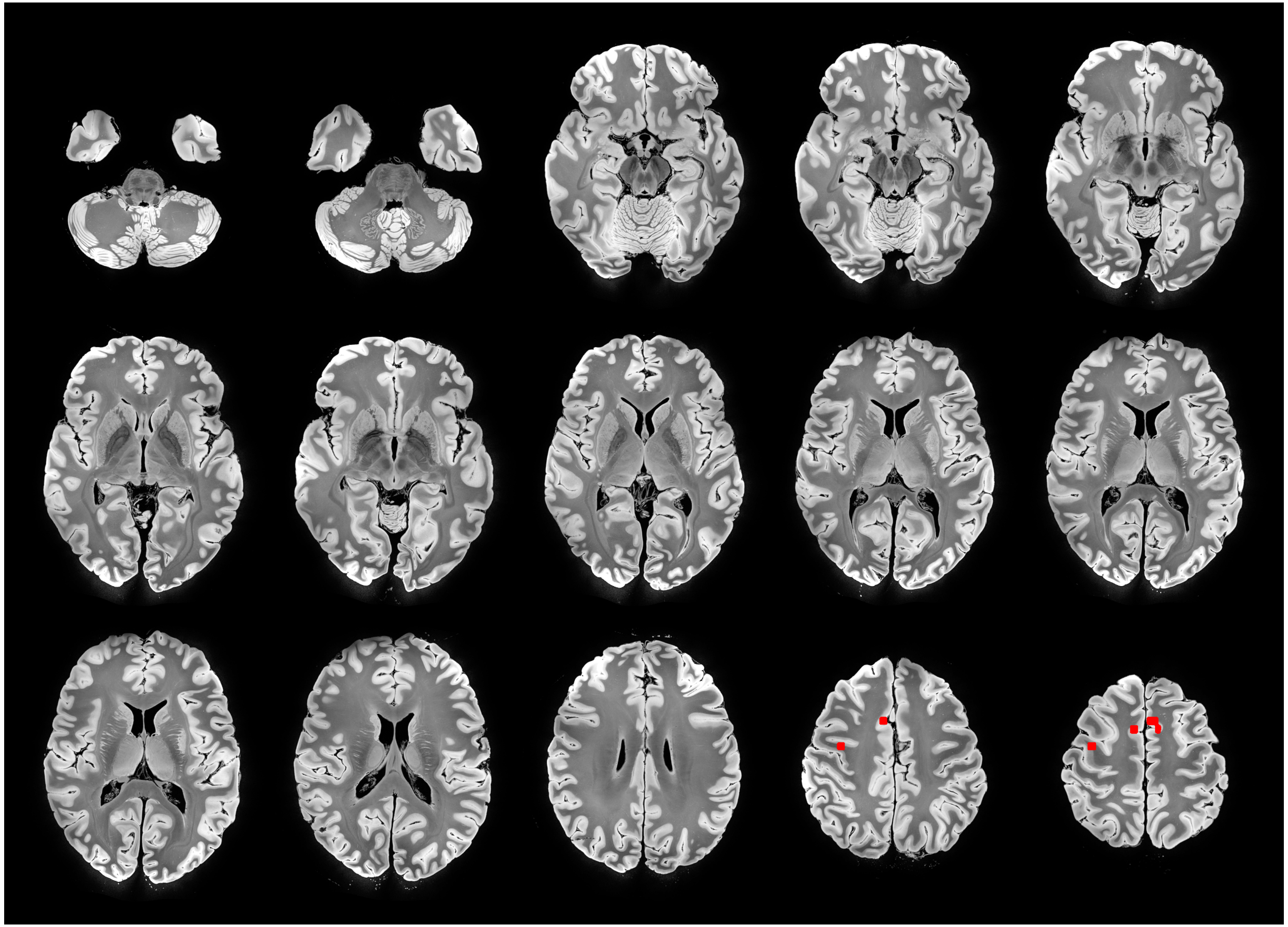

Common activation across all tasks

Common deactivation across all tasks

+2 sec

### Binary Conjunction Across All Modalities

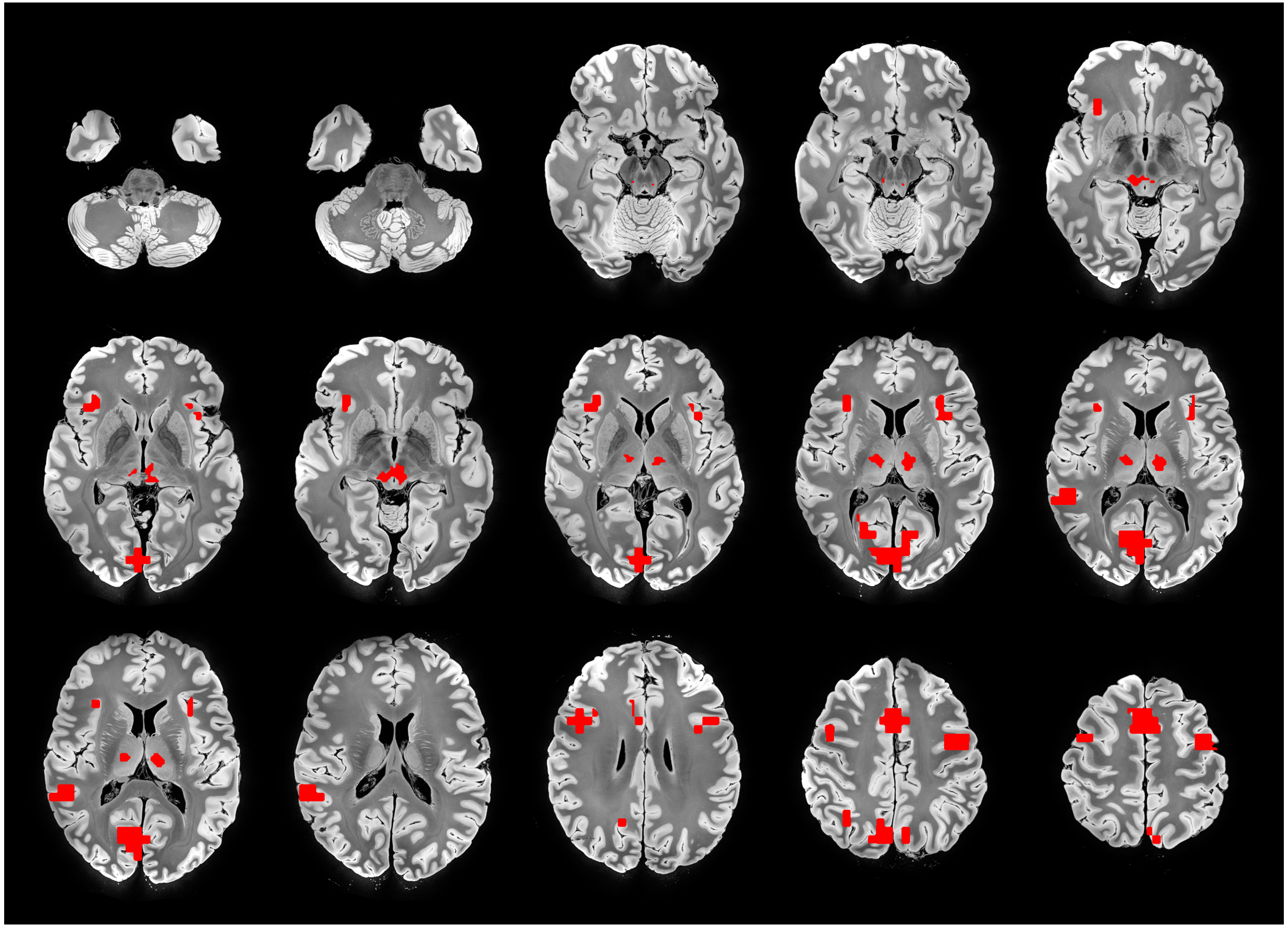

Common activation across all tasks

Common deactivation across all tasks

+4 sec

### Binary Conjunction Across All Modalities

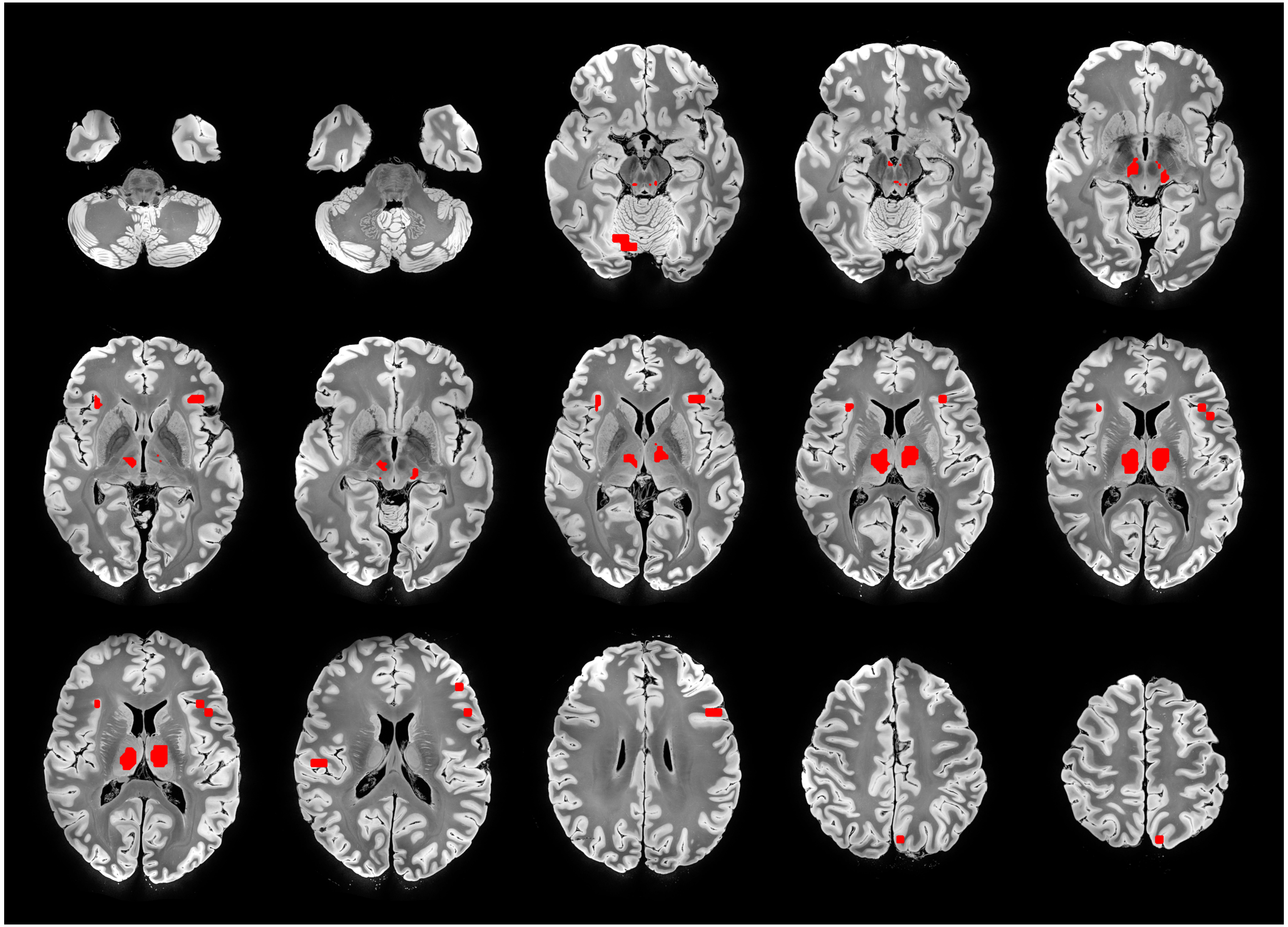

Common activation across all tasks

Common deactivation across all tasks

+6 sec

### Binary Conjunction Across All Modalities

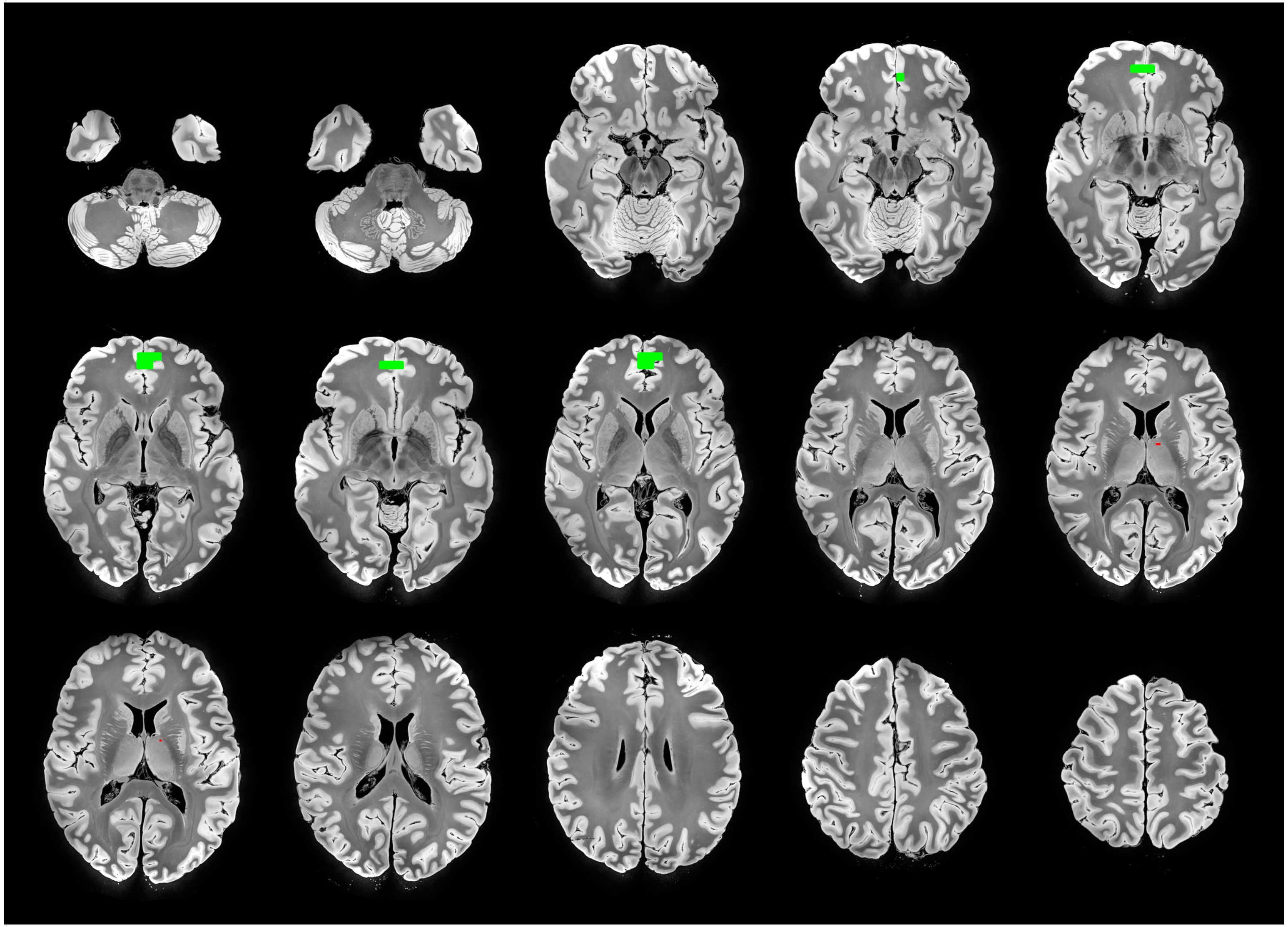

Common activation across all tasks

Common deactivation across all tasks

+8 sec

### Graded Conjunction Across All Modalities

### Graded Conjunction Across All Modalities: Increases

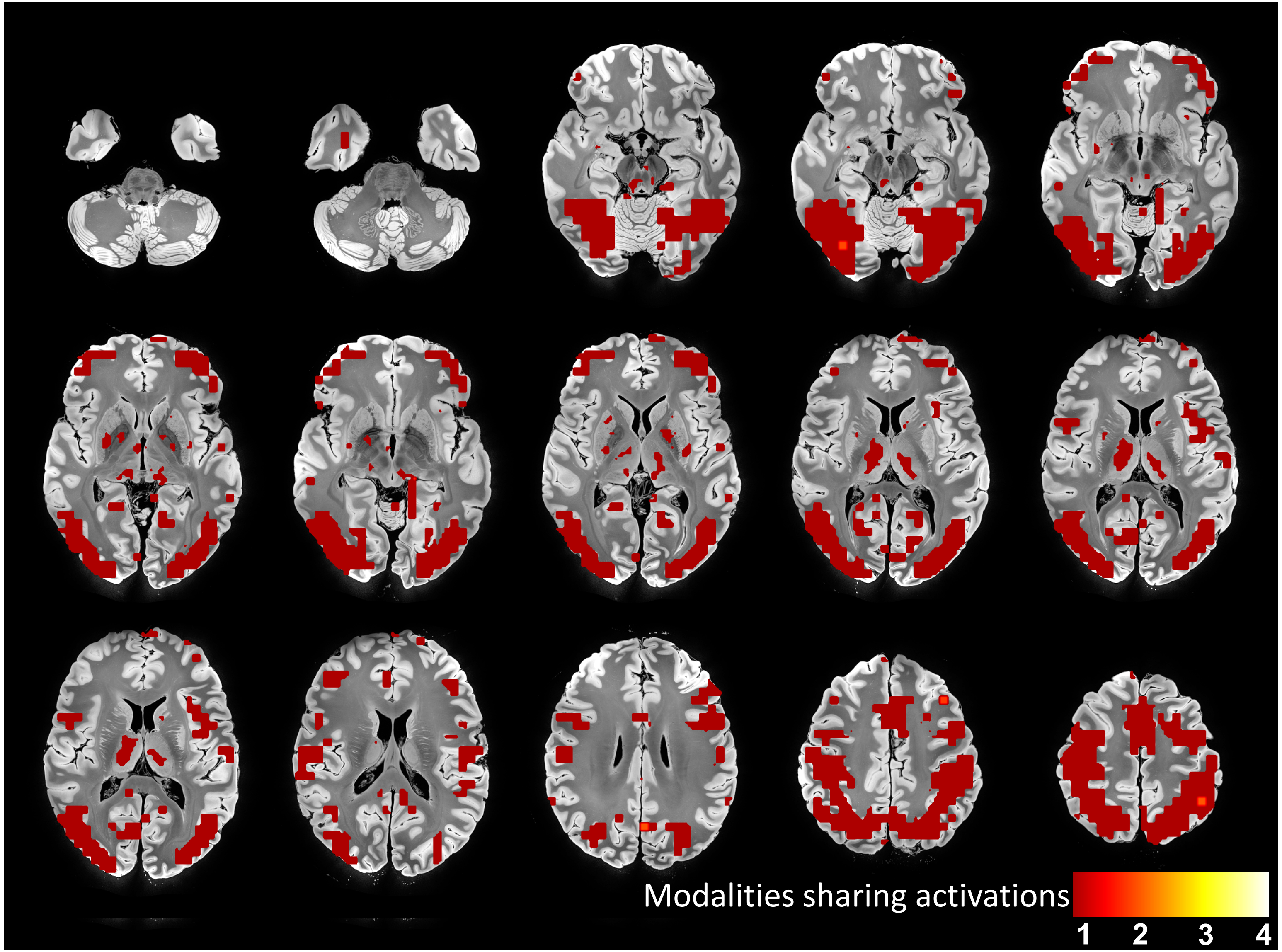


-4 sec

### Graded Conjunction Across All Modalities: Increases

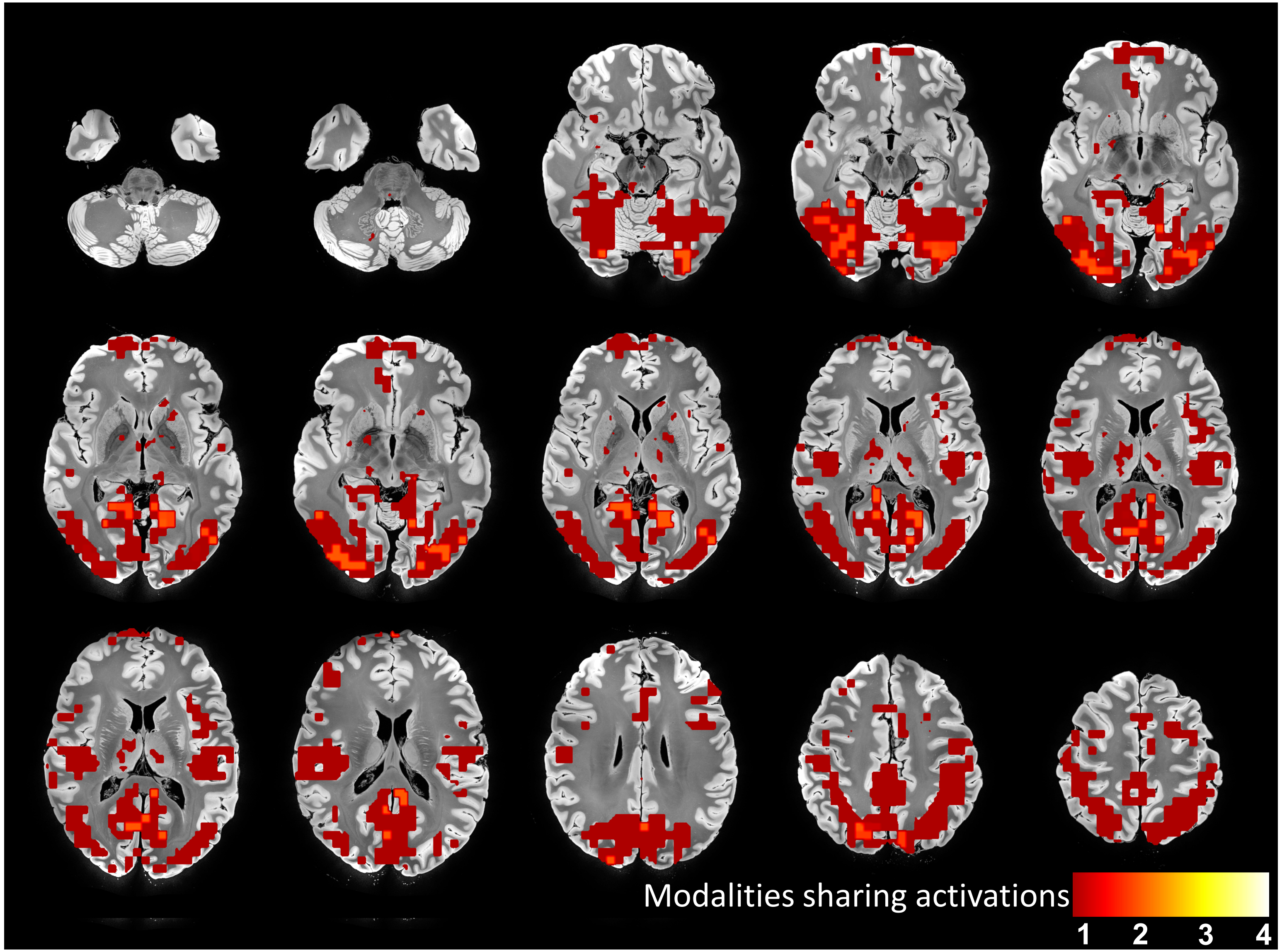


-2 sec

### Graded Conjunction Across All Modalities: Increases

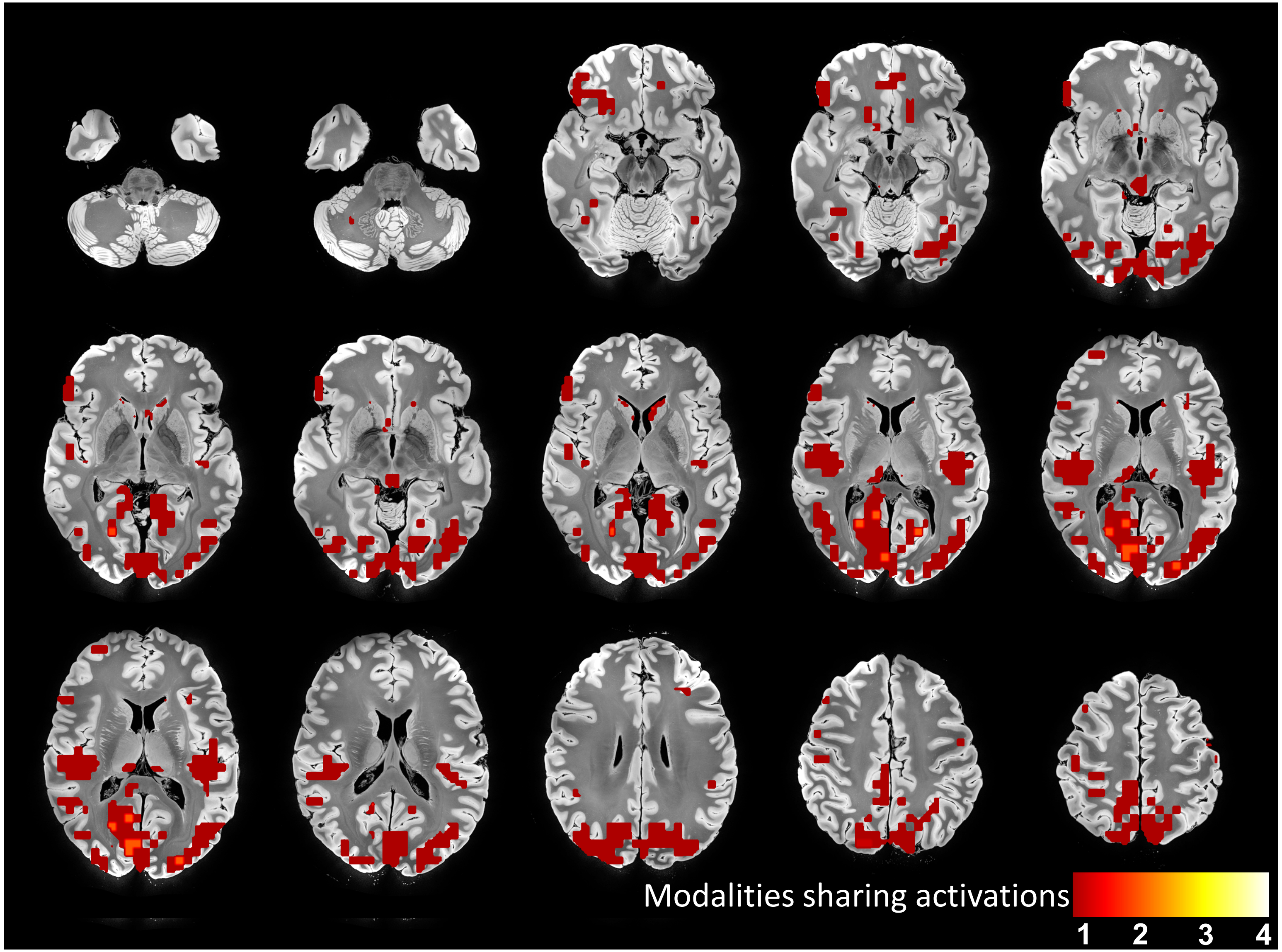


0 sec

### Graded Conjunction Across All Modalities: Increases

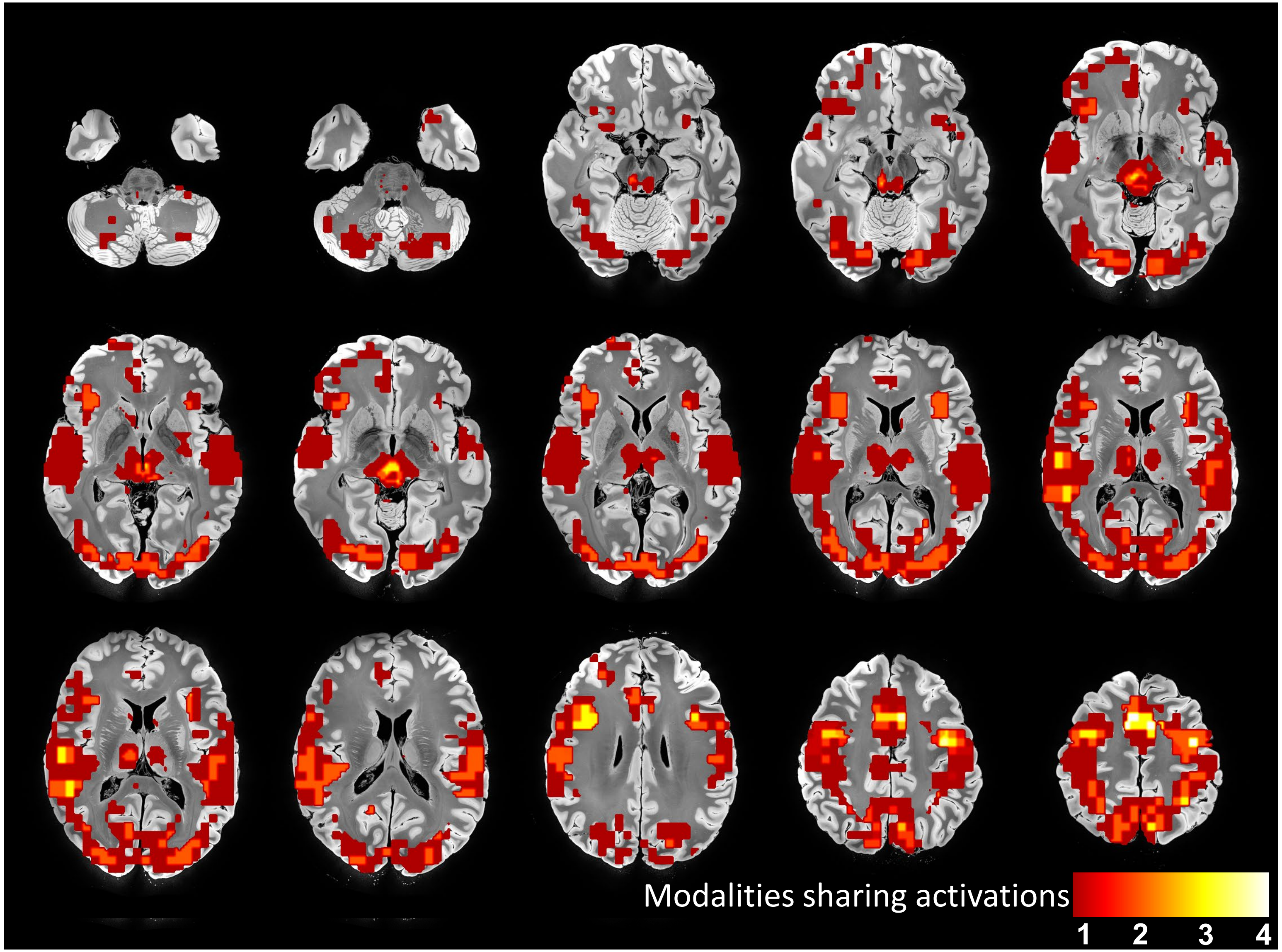


+2 sec

### Graded Conjunction Across All Modalities: Increases

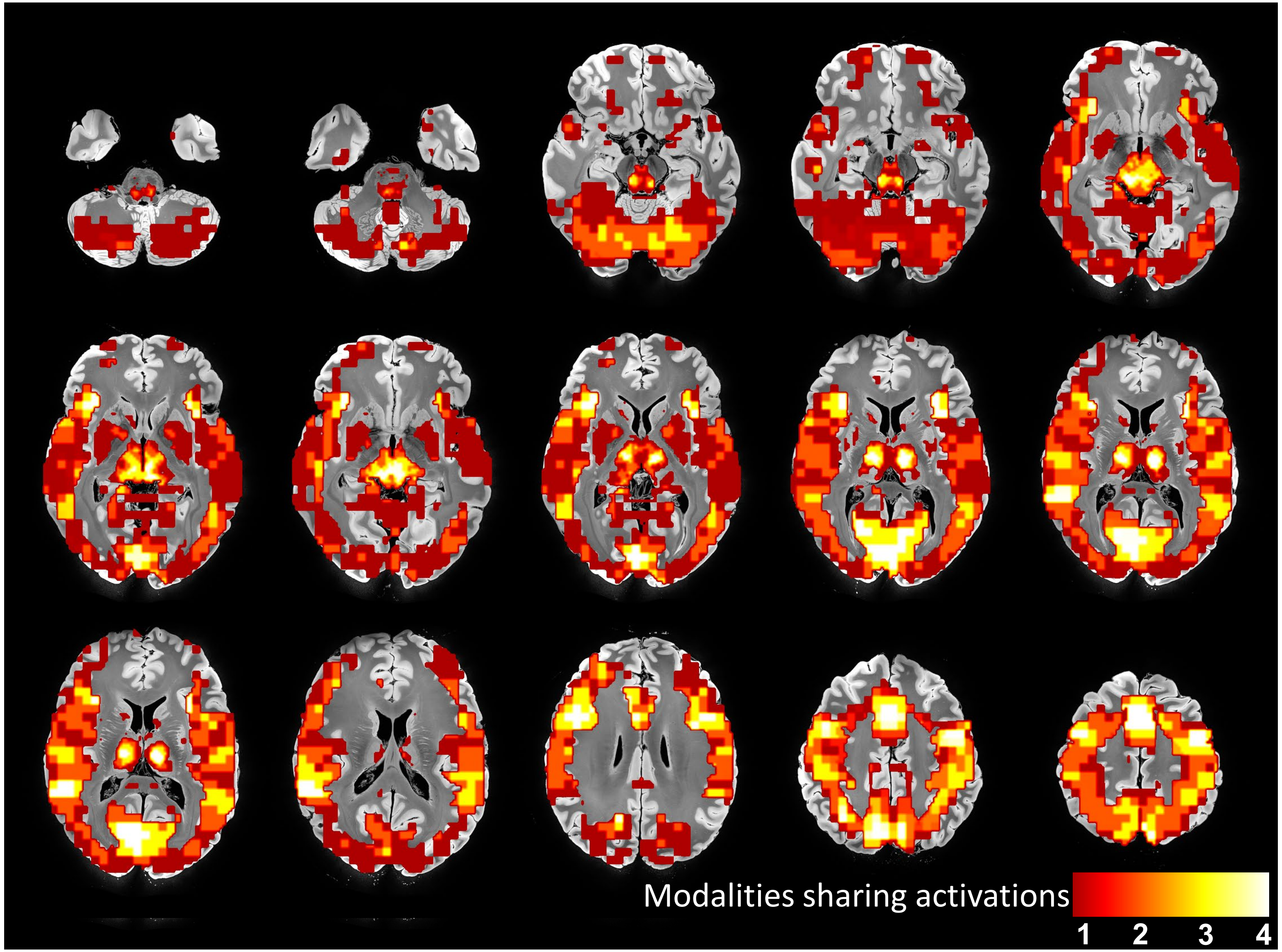


+4 sec

### Graded Conjunction Across All Modalities: Increases

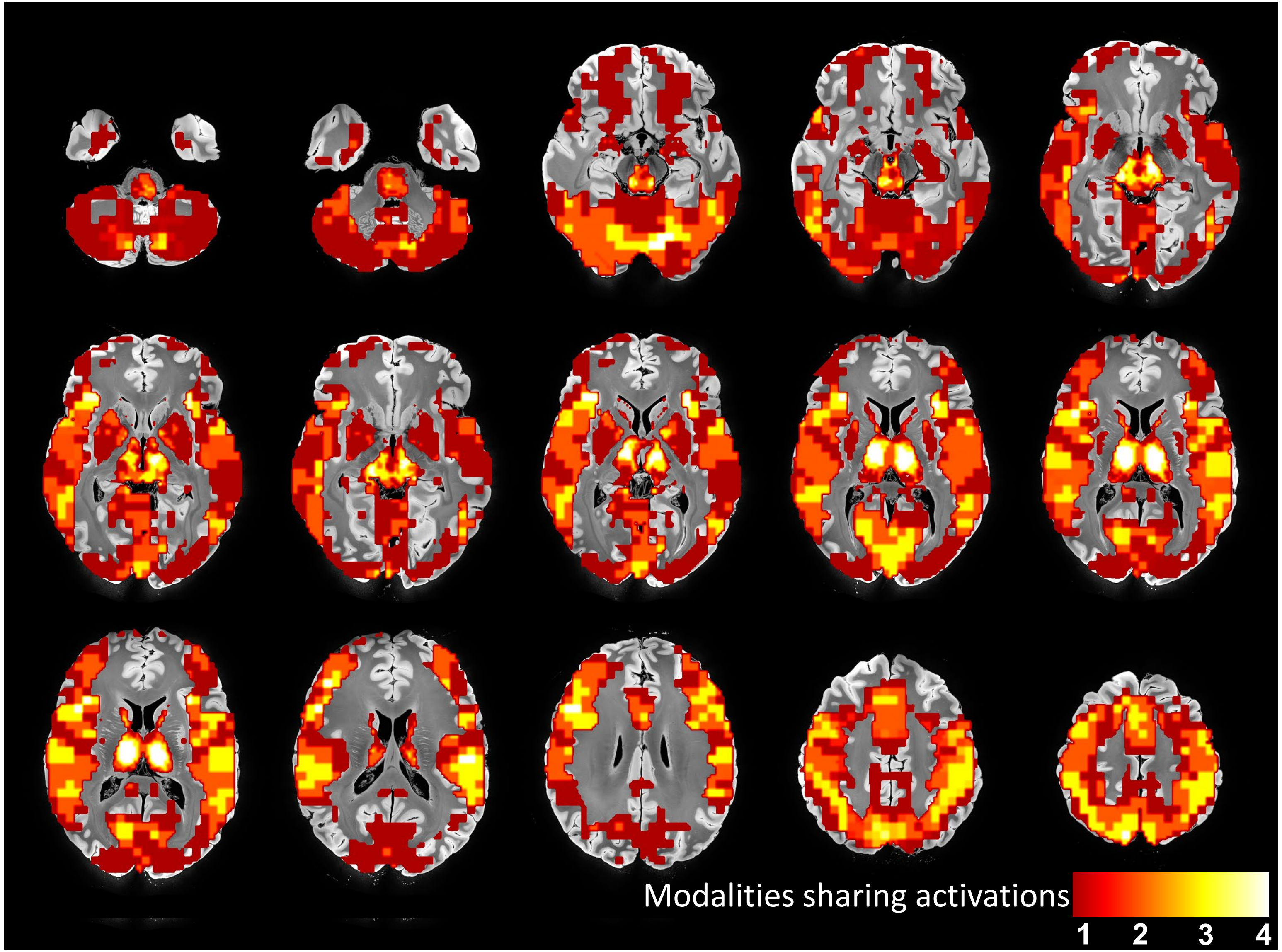


+6 sec

### Graded Conjunction Across All Modalities: Increases

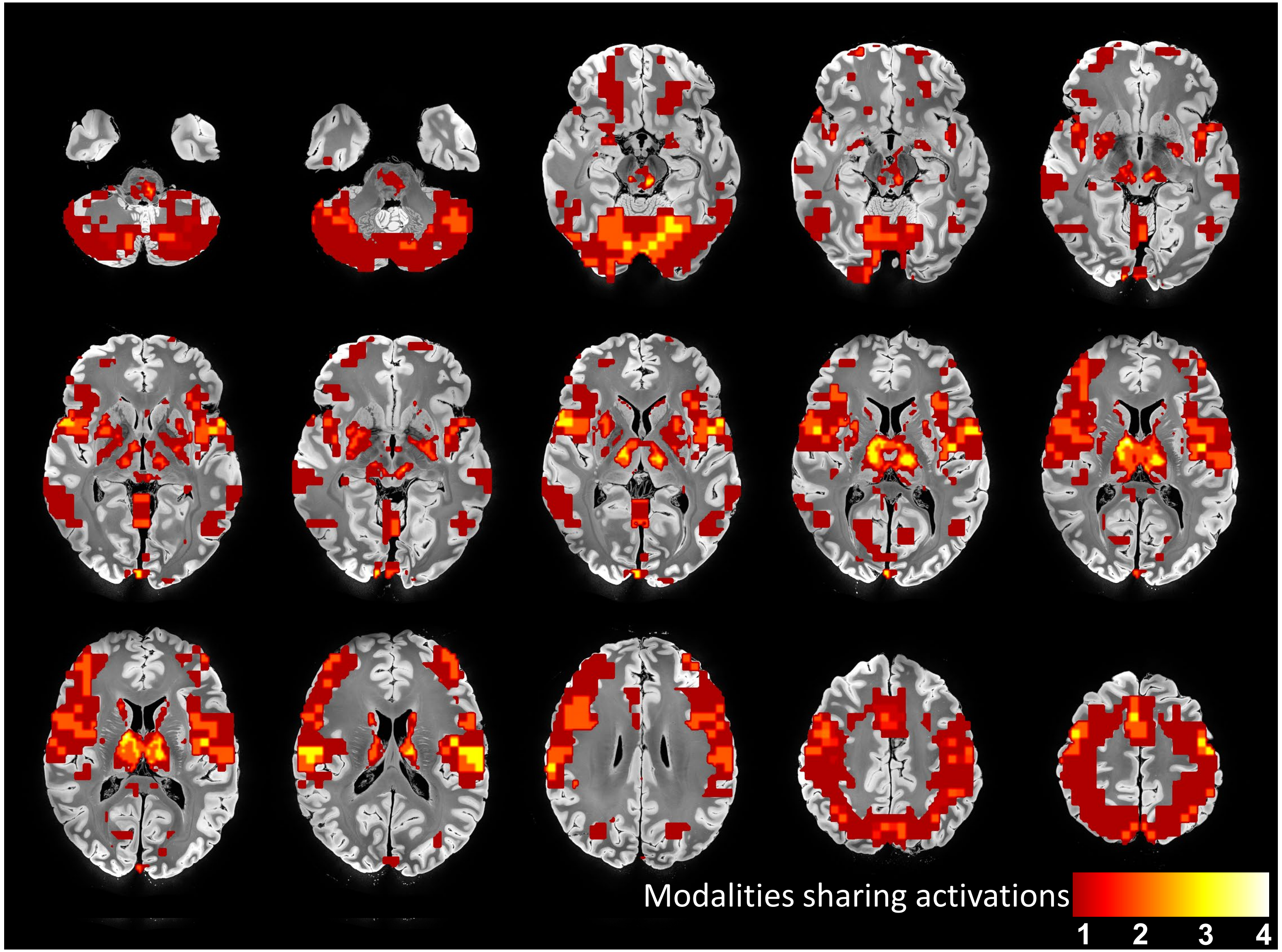


+8 sec

### Graded Conjunction Across All Modalities: Decreases

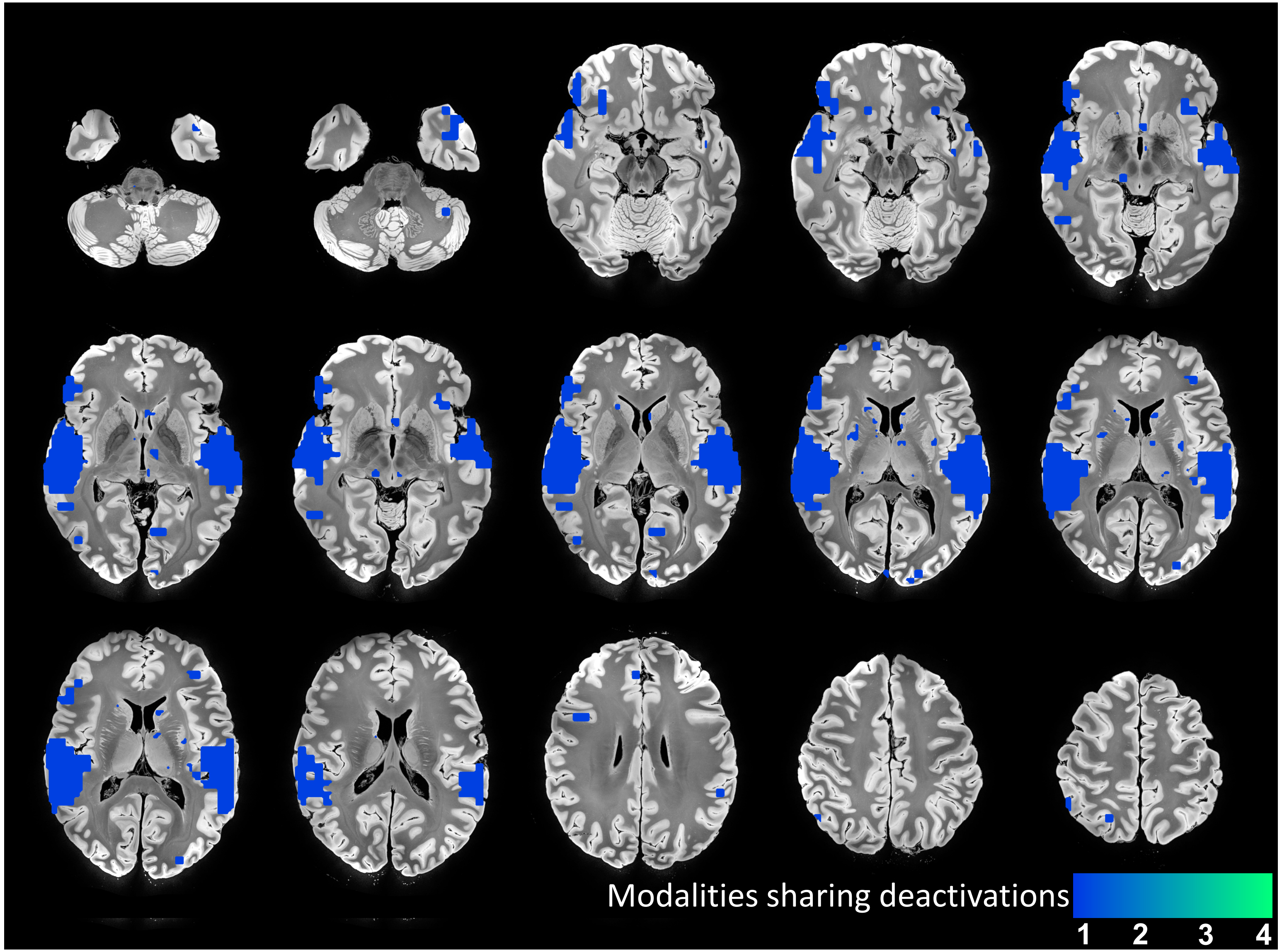


-4 sec

### Graded Conjunction Across All Modalities: Decreases

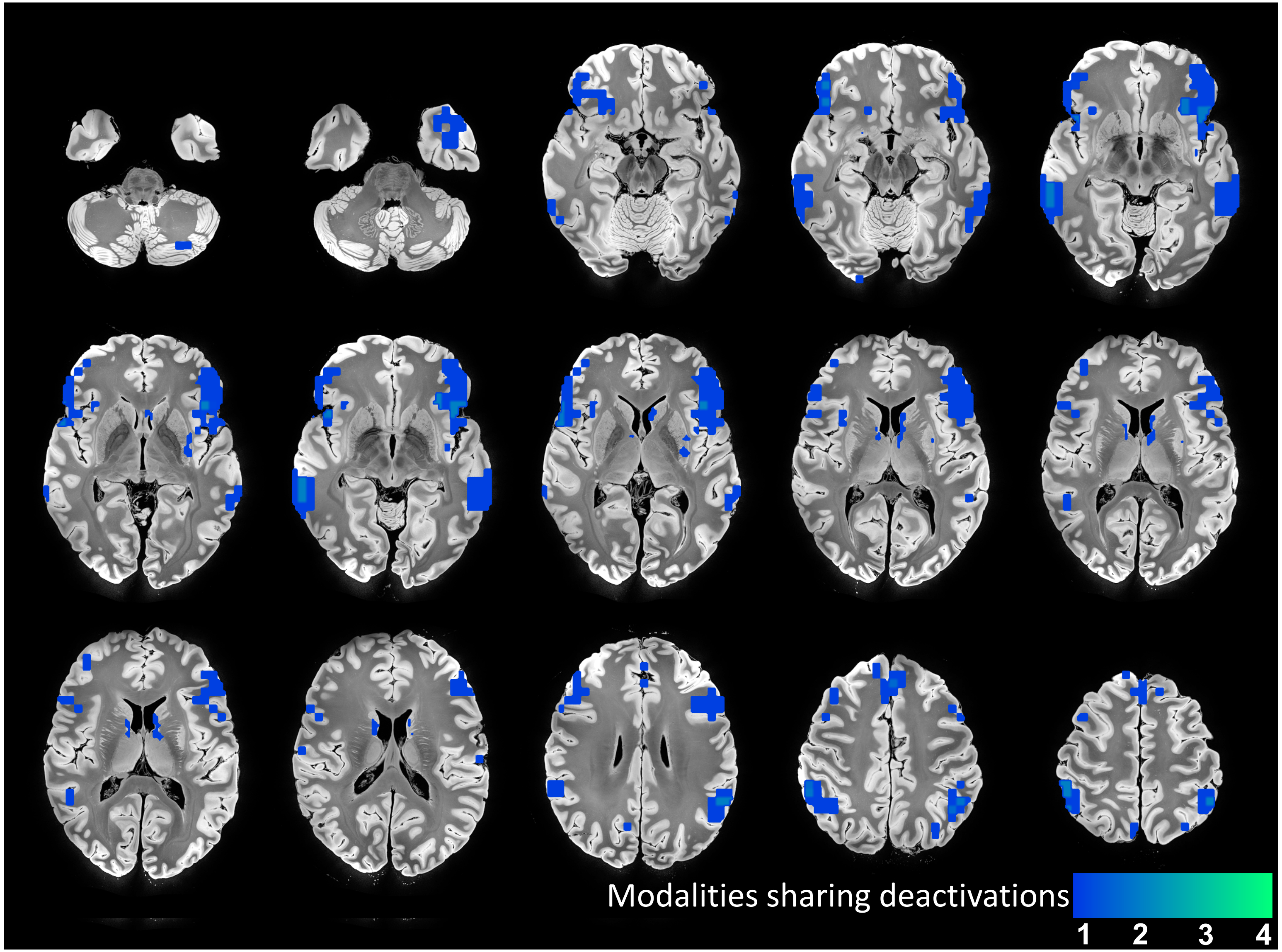


-2 sec

### Graded Conjunction Across All Modalities: Decreases

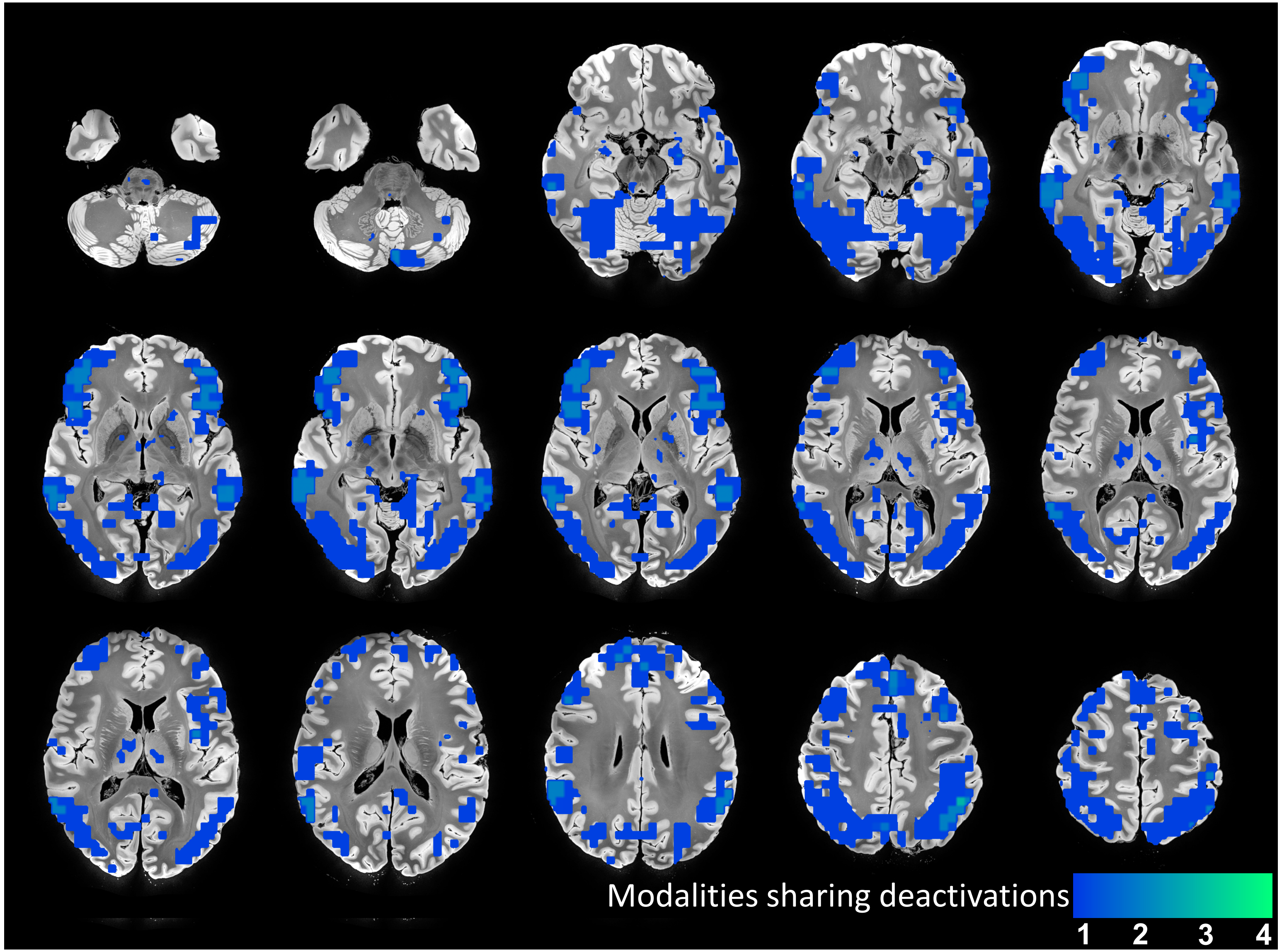


0 sec

### Graded Conjunction Across All Modalities: Decreases

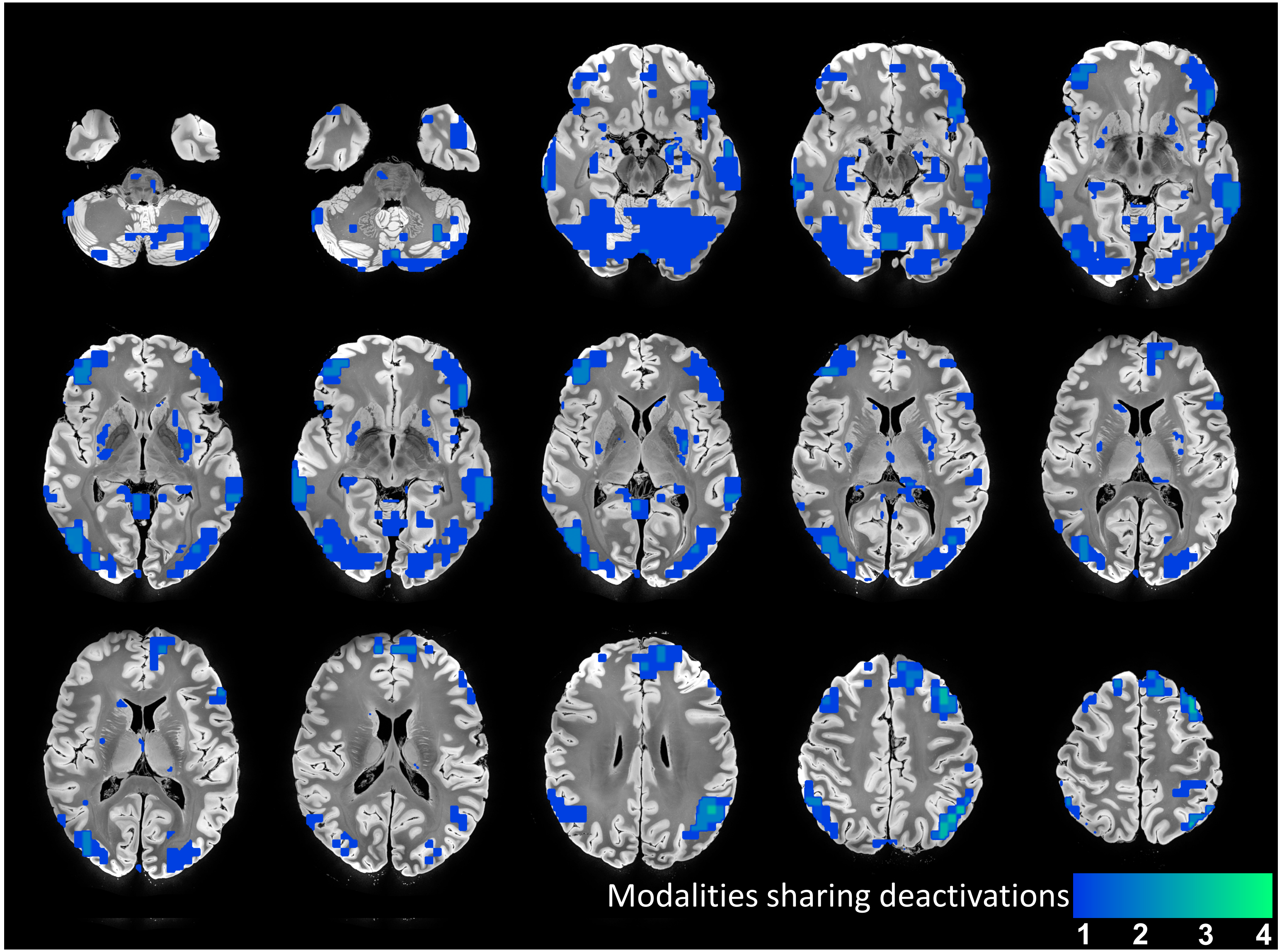


+2 sec

### Graded Conjunction Across All Modalities: Decreases

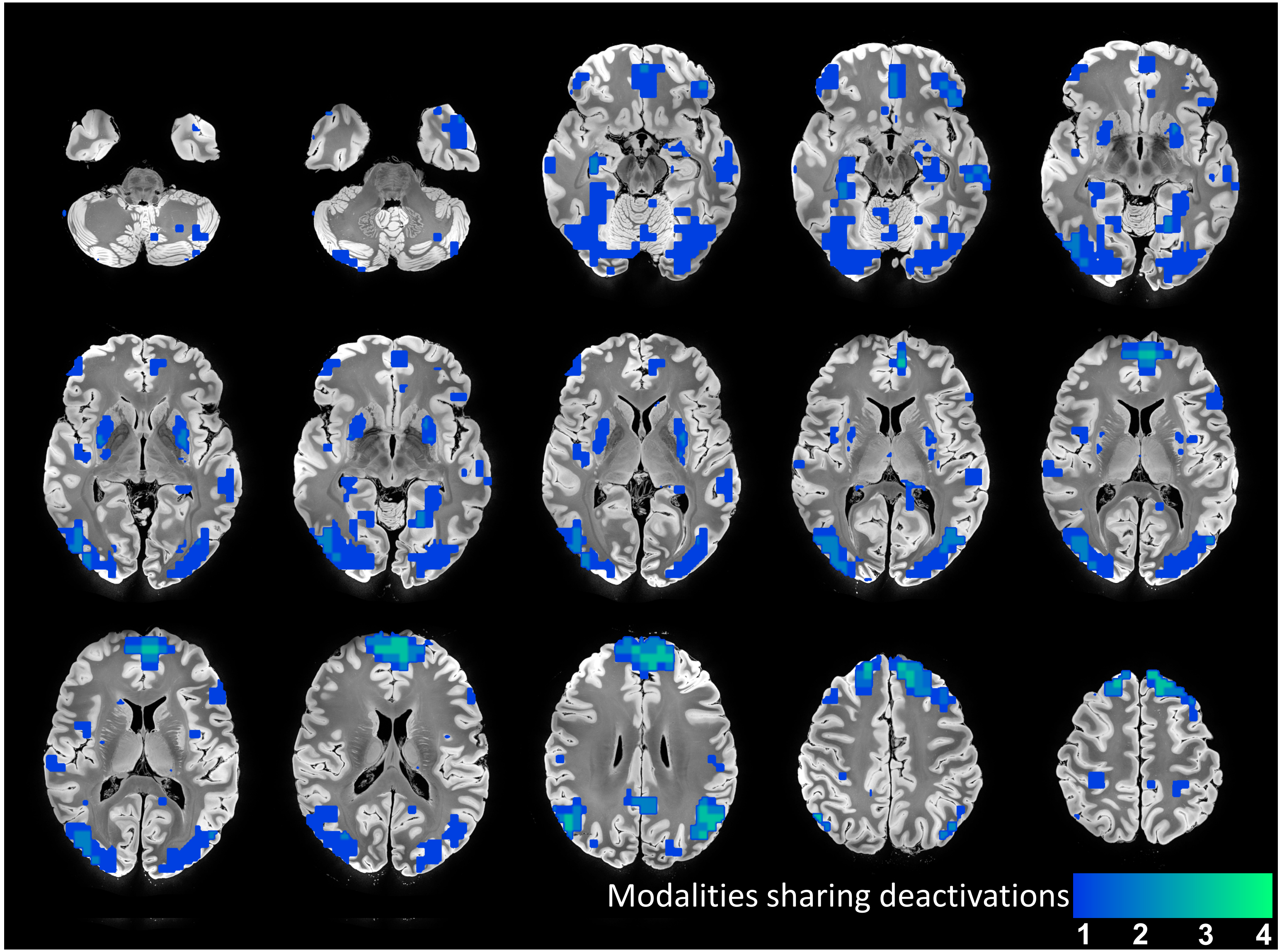


+4 sec

### Graded Conjunction Across All Modalities: Decreases

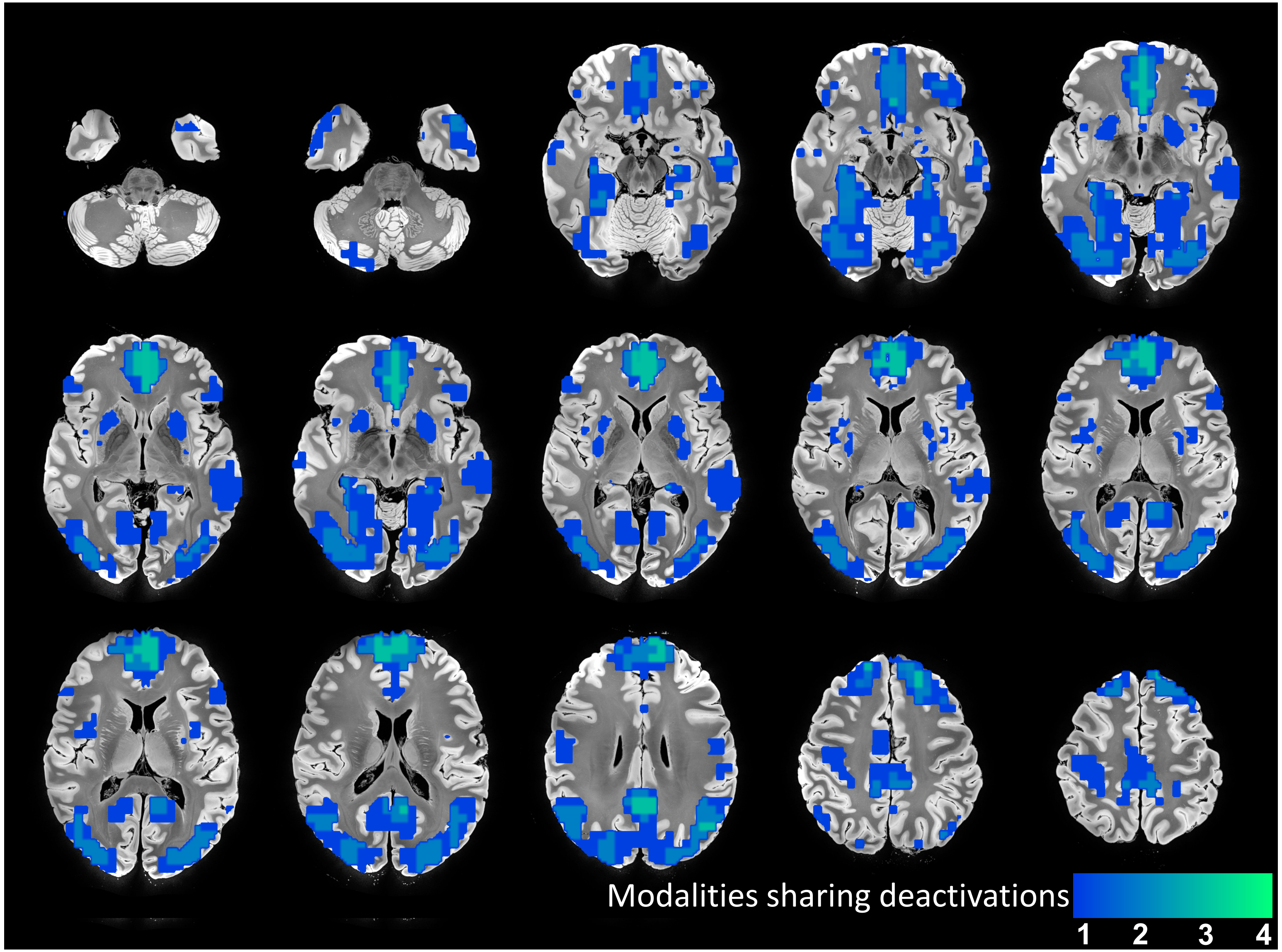


+6 sec

### Graded Conjunction Across All Modalities: Decreases

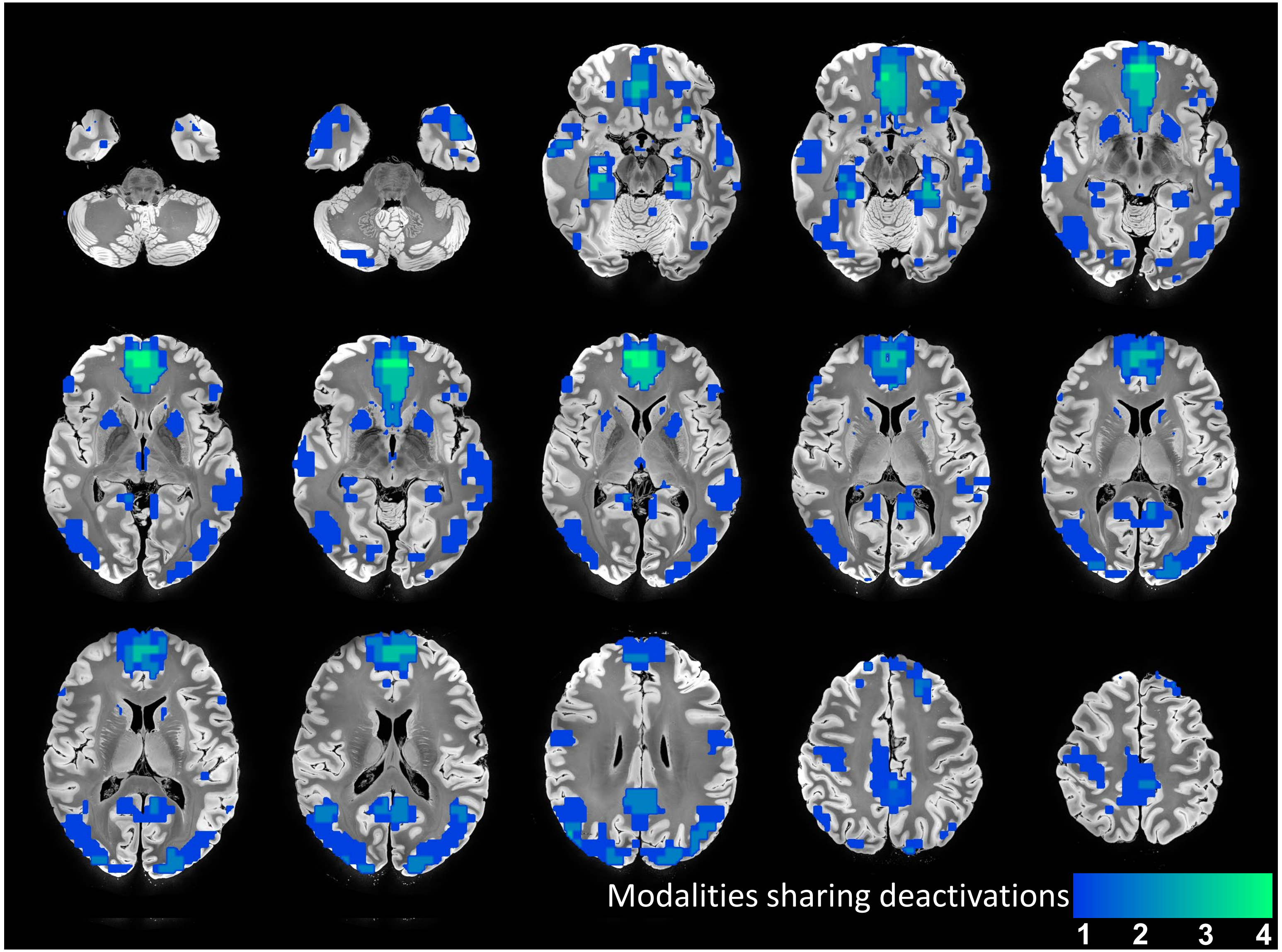


+8 sec
