## Supplementary Presentation S2 for "Shared subcortical arousal systems across sensory modalities during transient modulation of attention"

### Binary Conjunction Within Each Modality

### Visual Conjunction

### Visual Conjunction

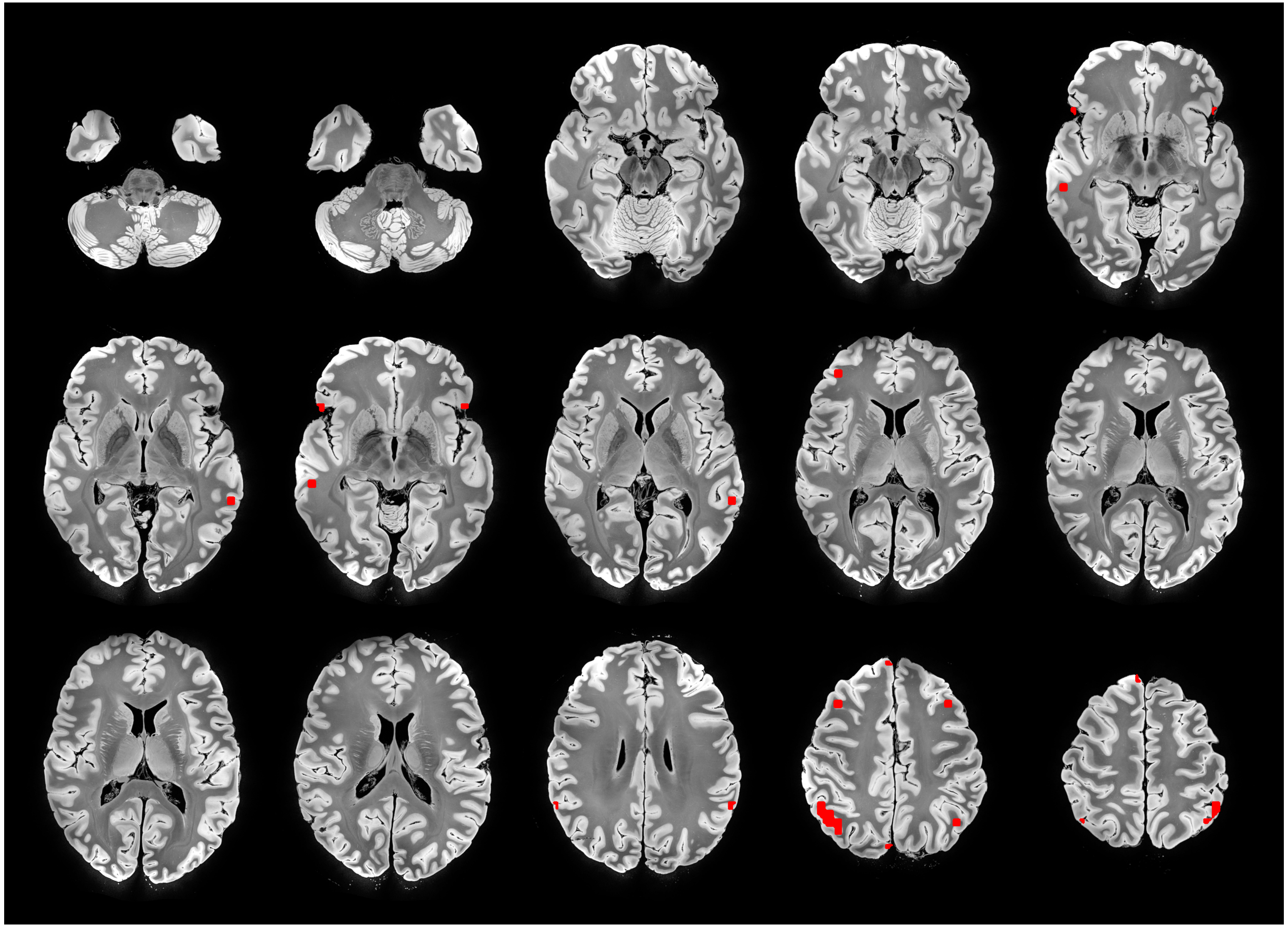

Common activation across all tasks

Common deactivation across all tasks

-4 sec

### Visual Conjunction

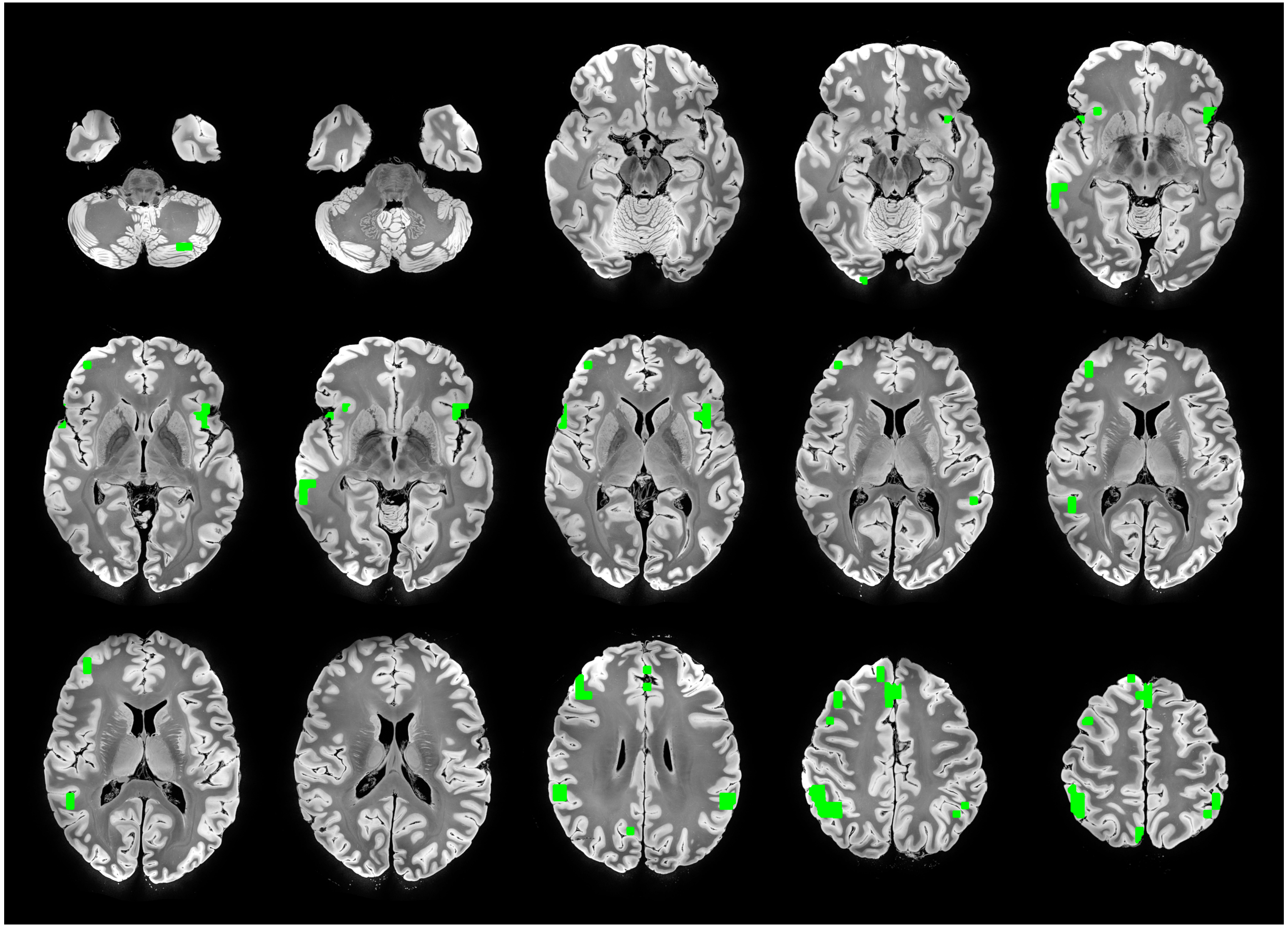

Common activation across all tasks

Common deactivation across all tasks

-2 sec

### Visual Conjunction

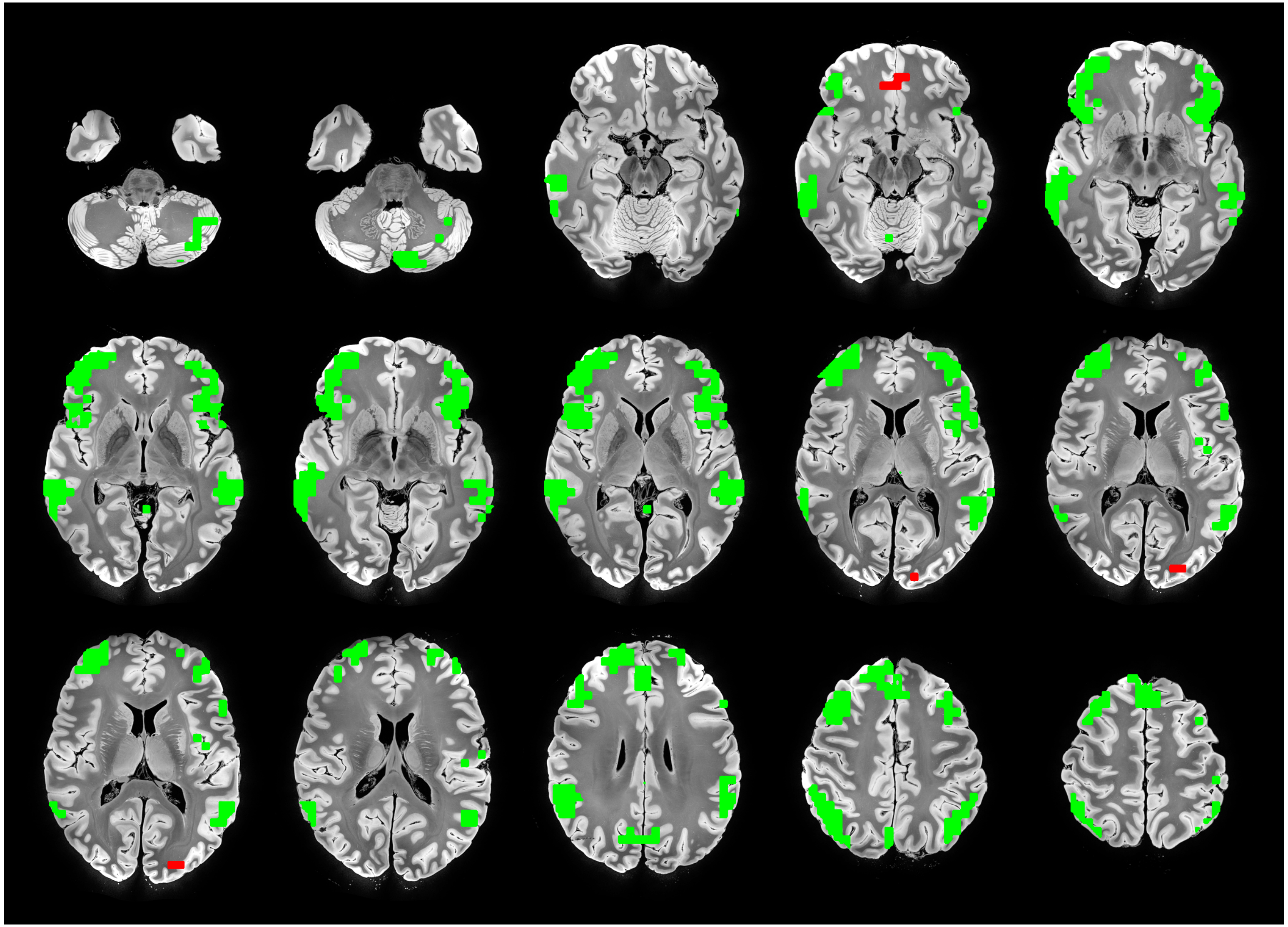

Common activation across all tasks    Common deactivation across all tasks

0 sec

### Visual Conjunction

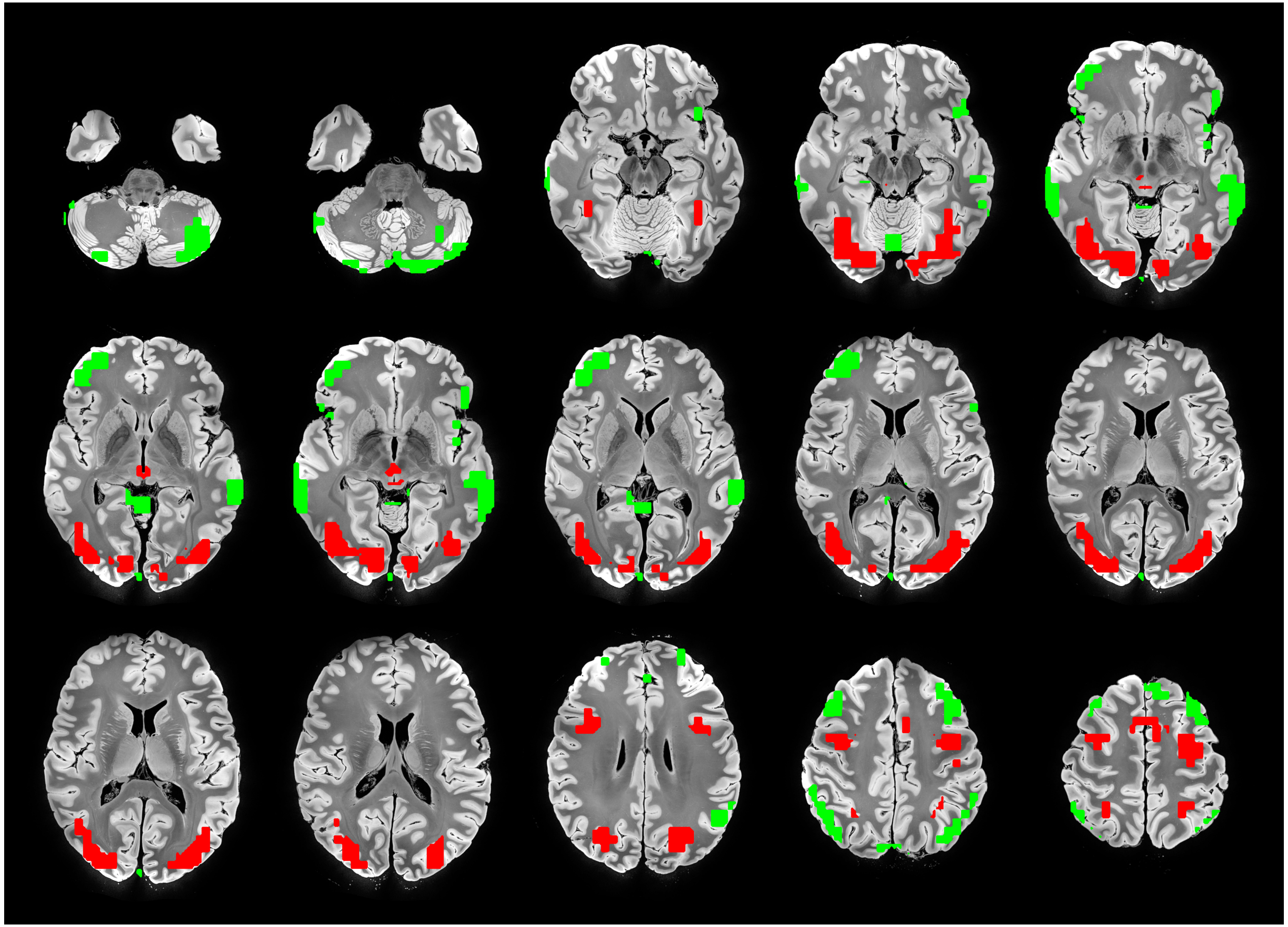

Common activation across all tasks

Common deactivation across all tasks

+2 sec

### Visual Conjunction

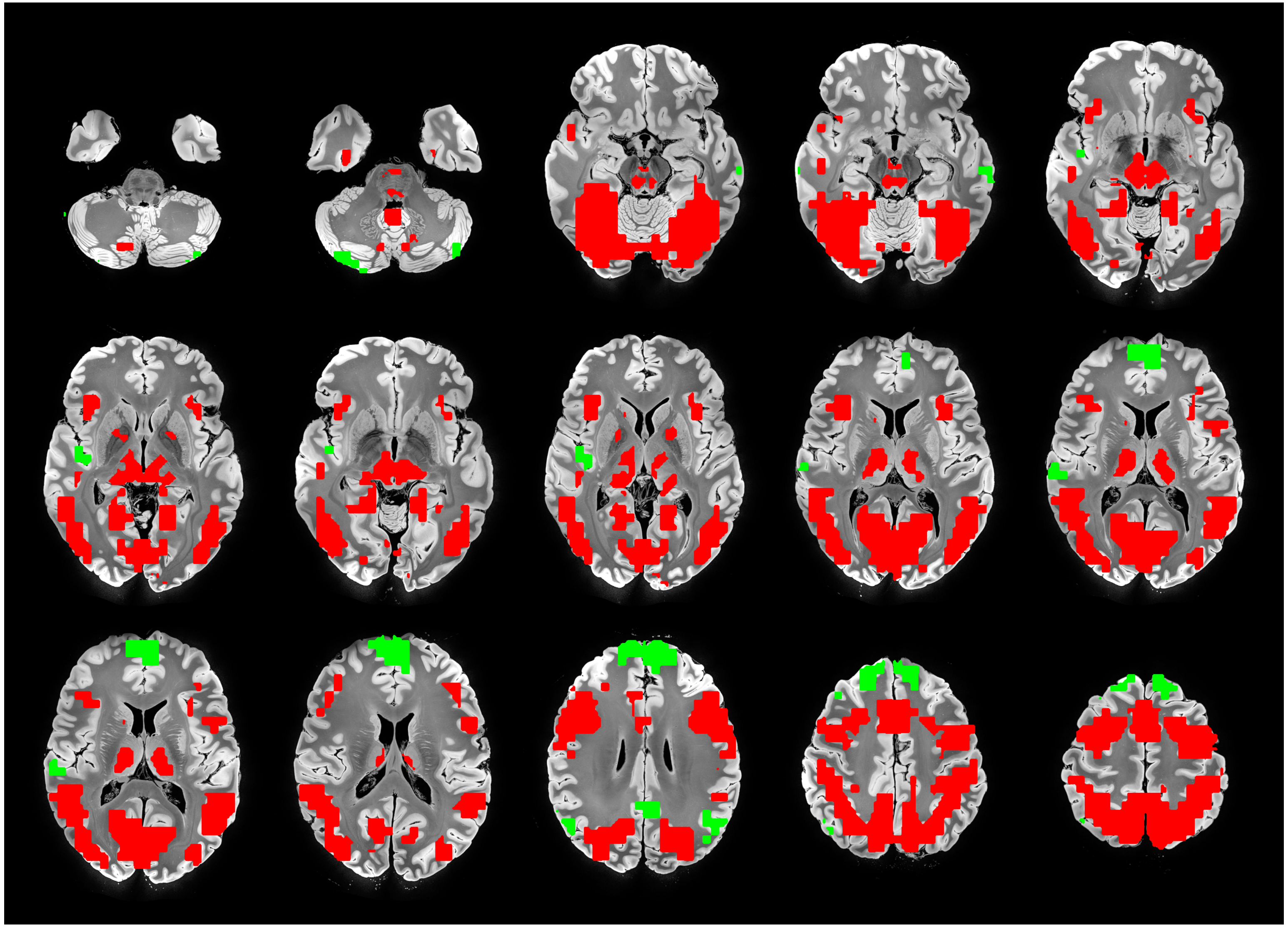

Common activation across all tasks

Common deactivation across all tasks

+4 sec

### Visual Conjunction

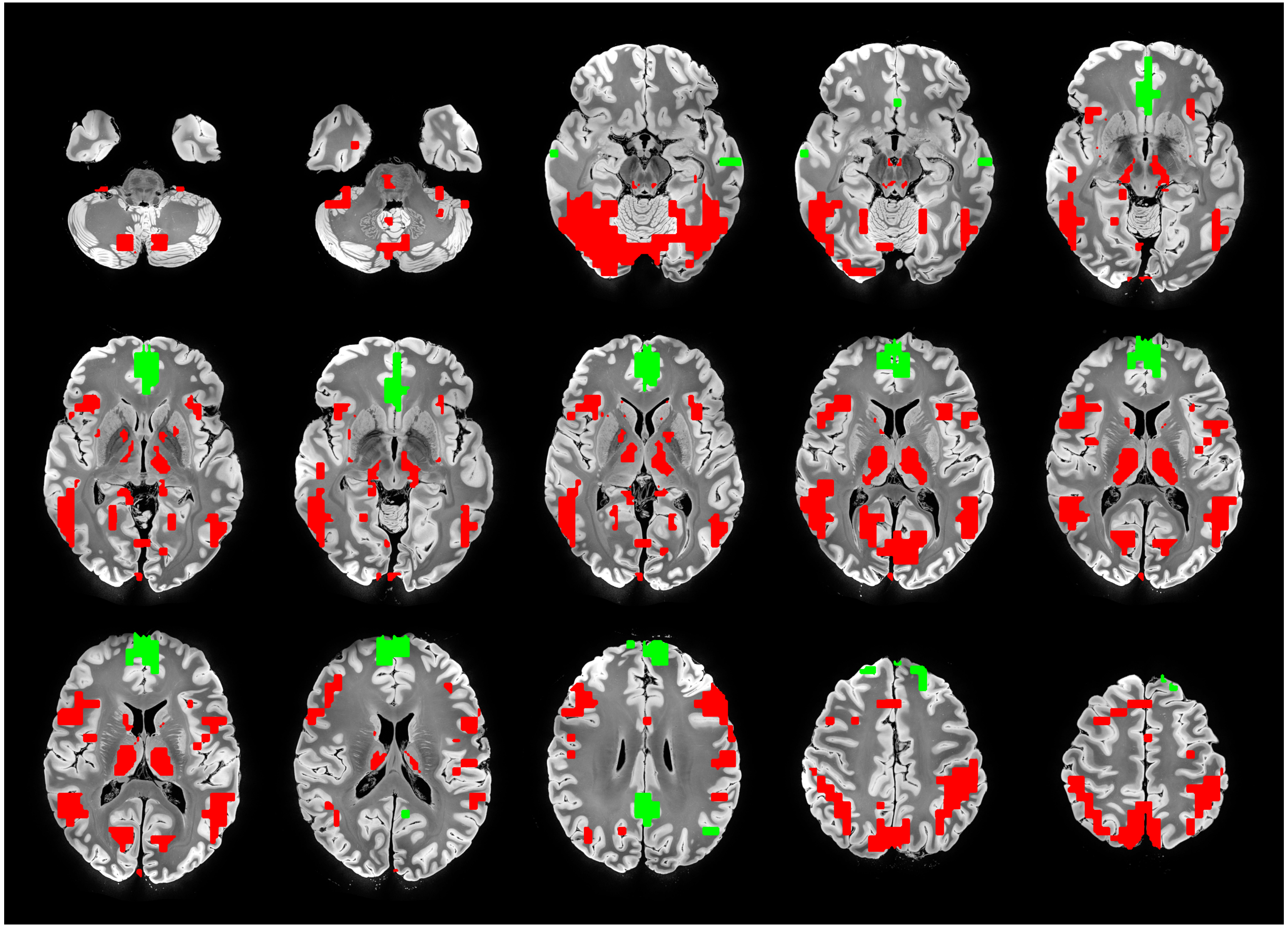

Common activation across all tasks   Common deactivation across all tasks

+6 sec

### Visual Conjunction

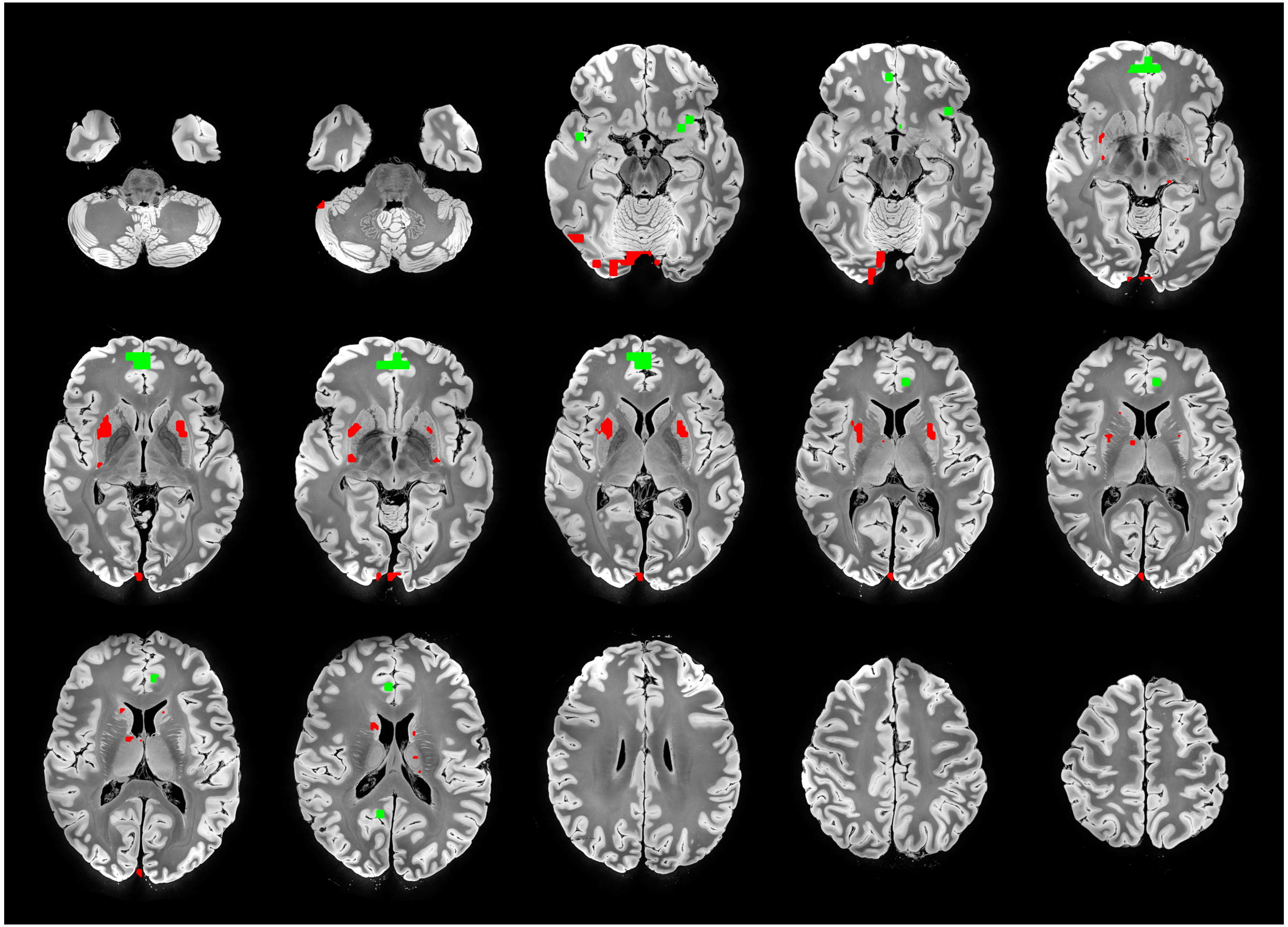

Common activation across all tasks

Common deactivation across all tasks

+8 sec

### Auditory Conjunction

### Auditory Conjunction

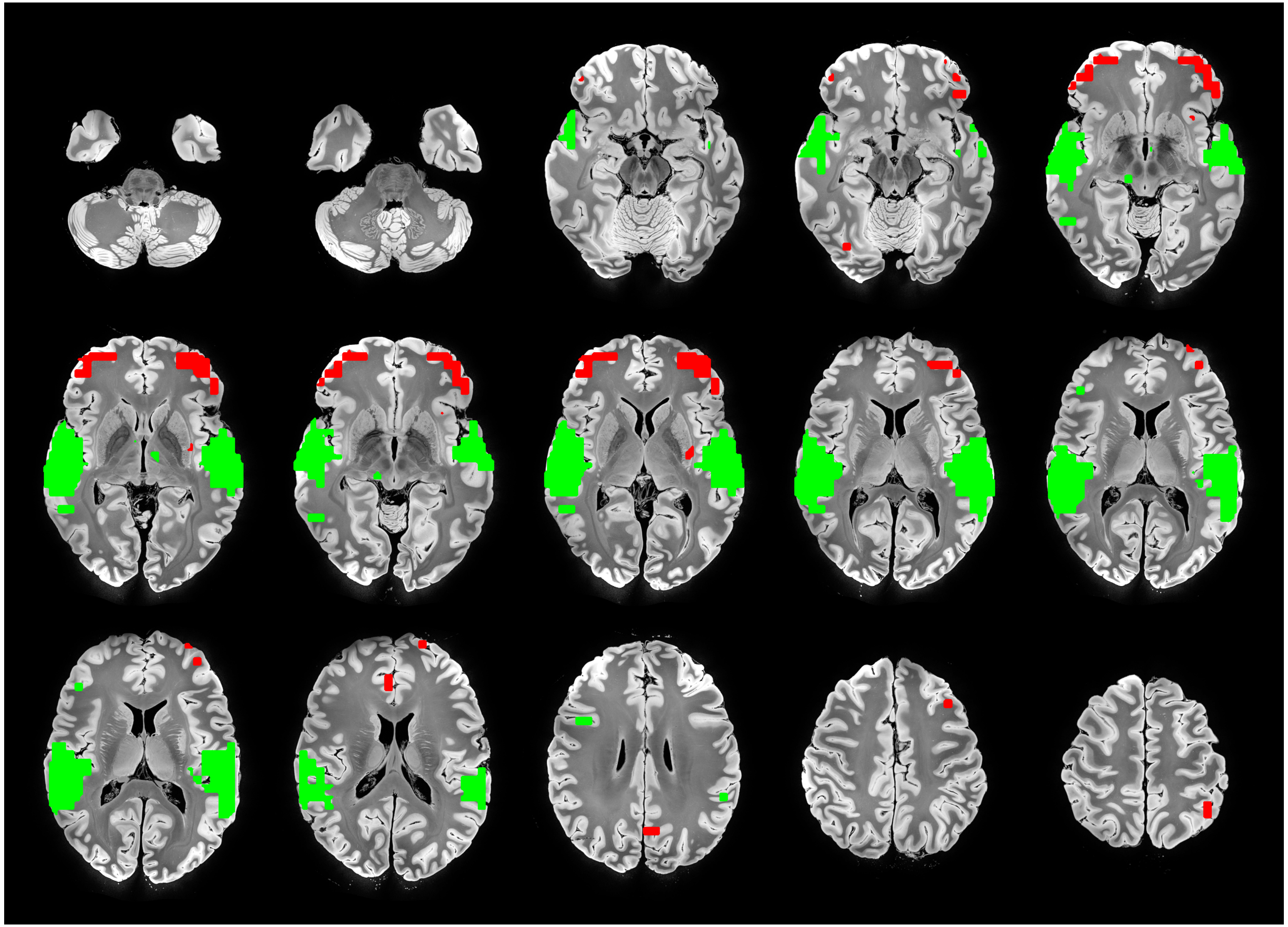

Common activation across all tasks

Common deactivation across all tasks

-4 sec

### Auditory Conjunction

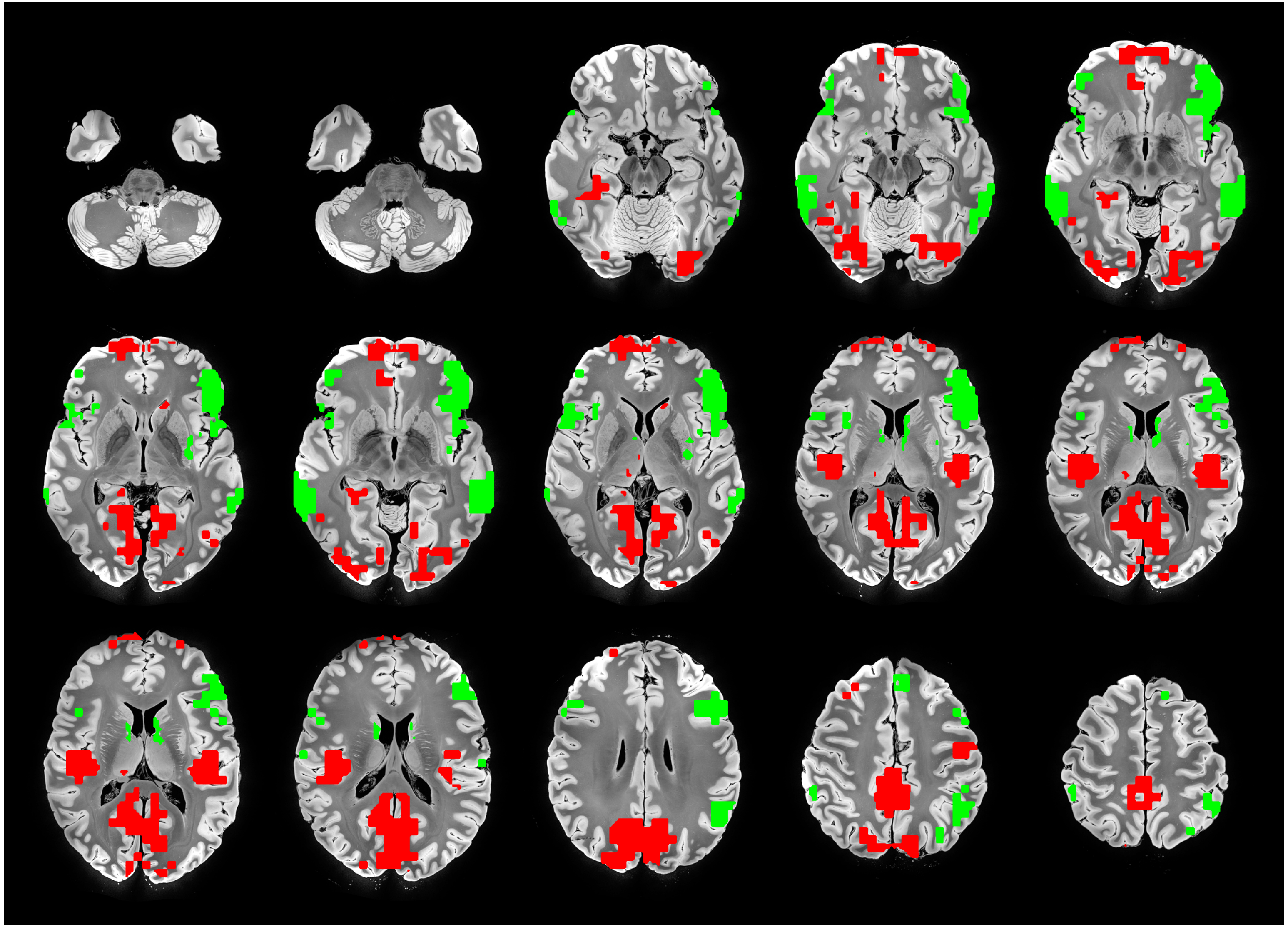

Common activation across all tasks

Common deactivation across all tasks

-2 sec

### Auditory Conjunction

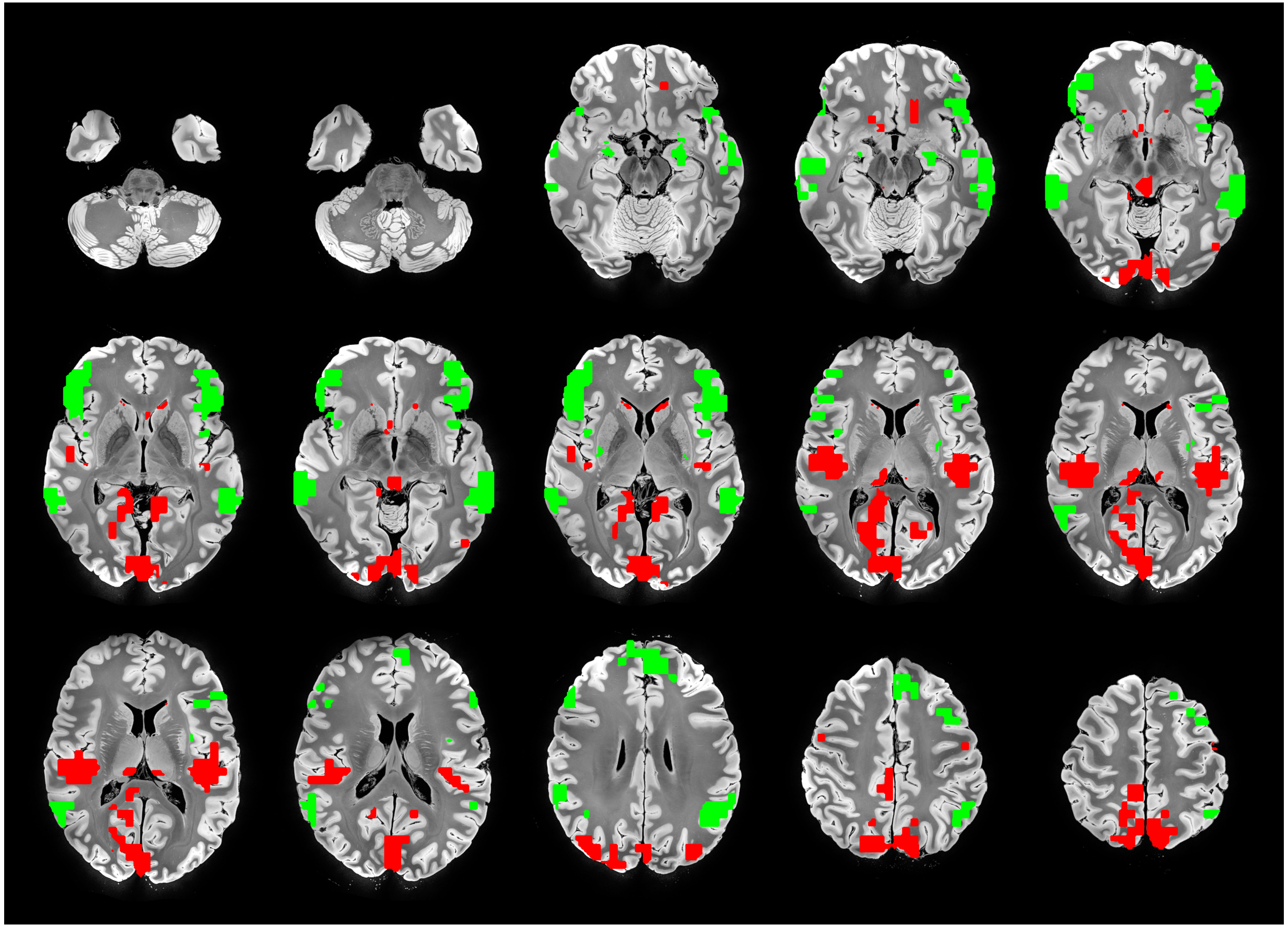

Common activation across all tasks

Common deactivation across all tasks

0 sec

### Auditory Conjunction

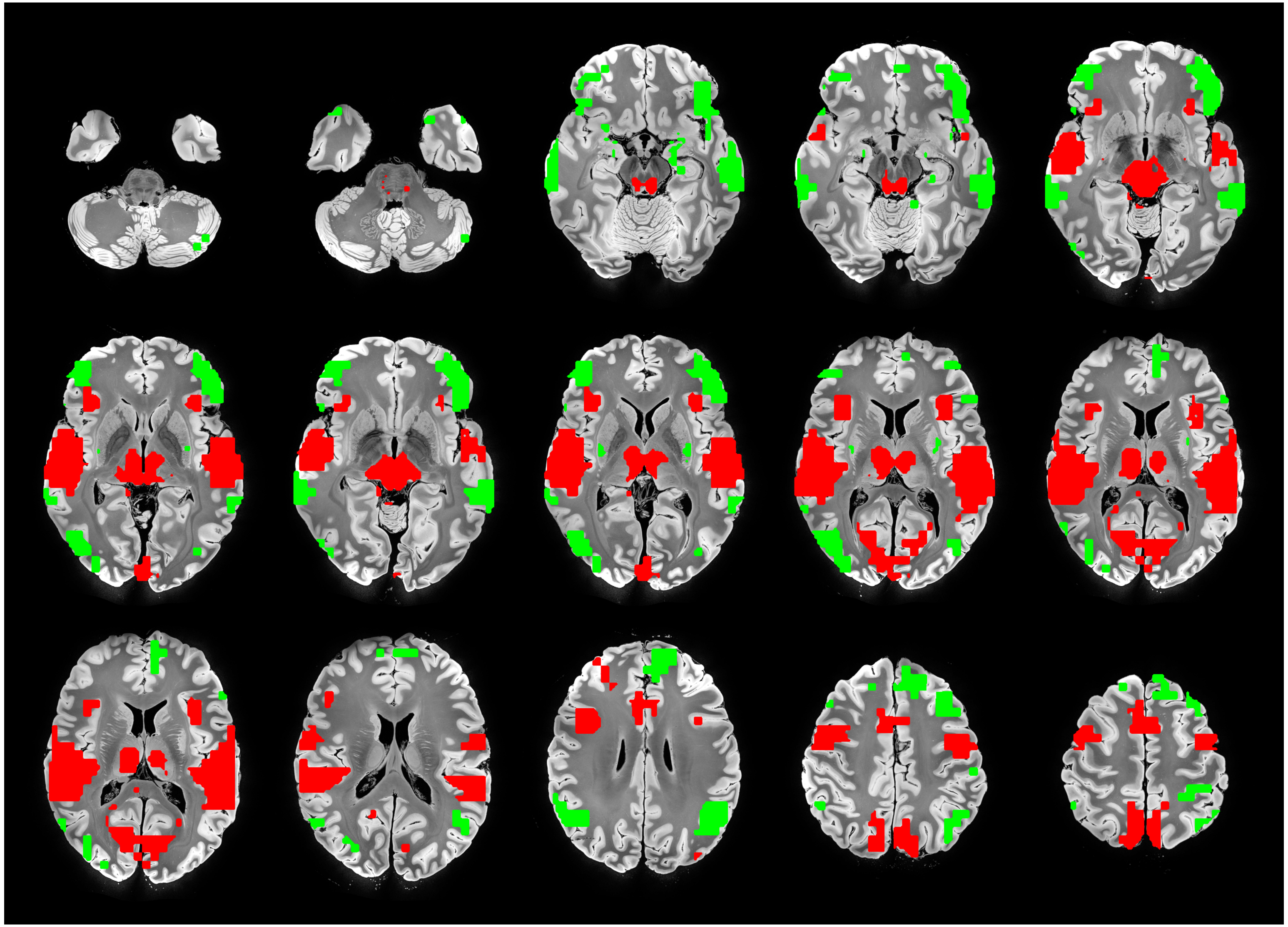

Common activation across all tasks

Common deactivation across all tasks

+2 sec

### Auditory Conjunction

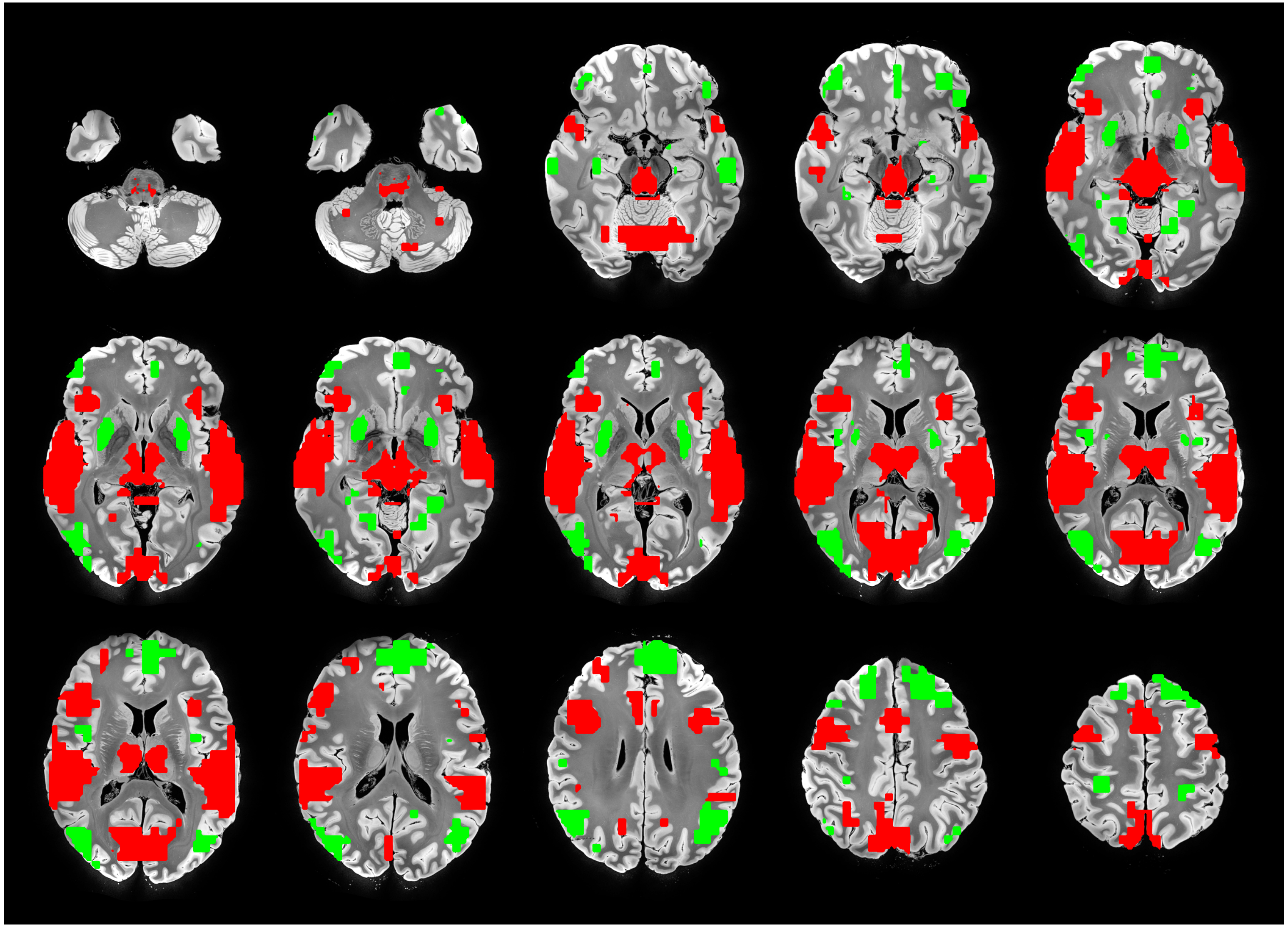

Common activation across all tasks    Common deactivation across all tasks

+4 sec

### Auditory Conjunction

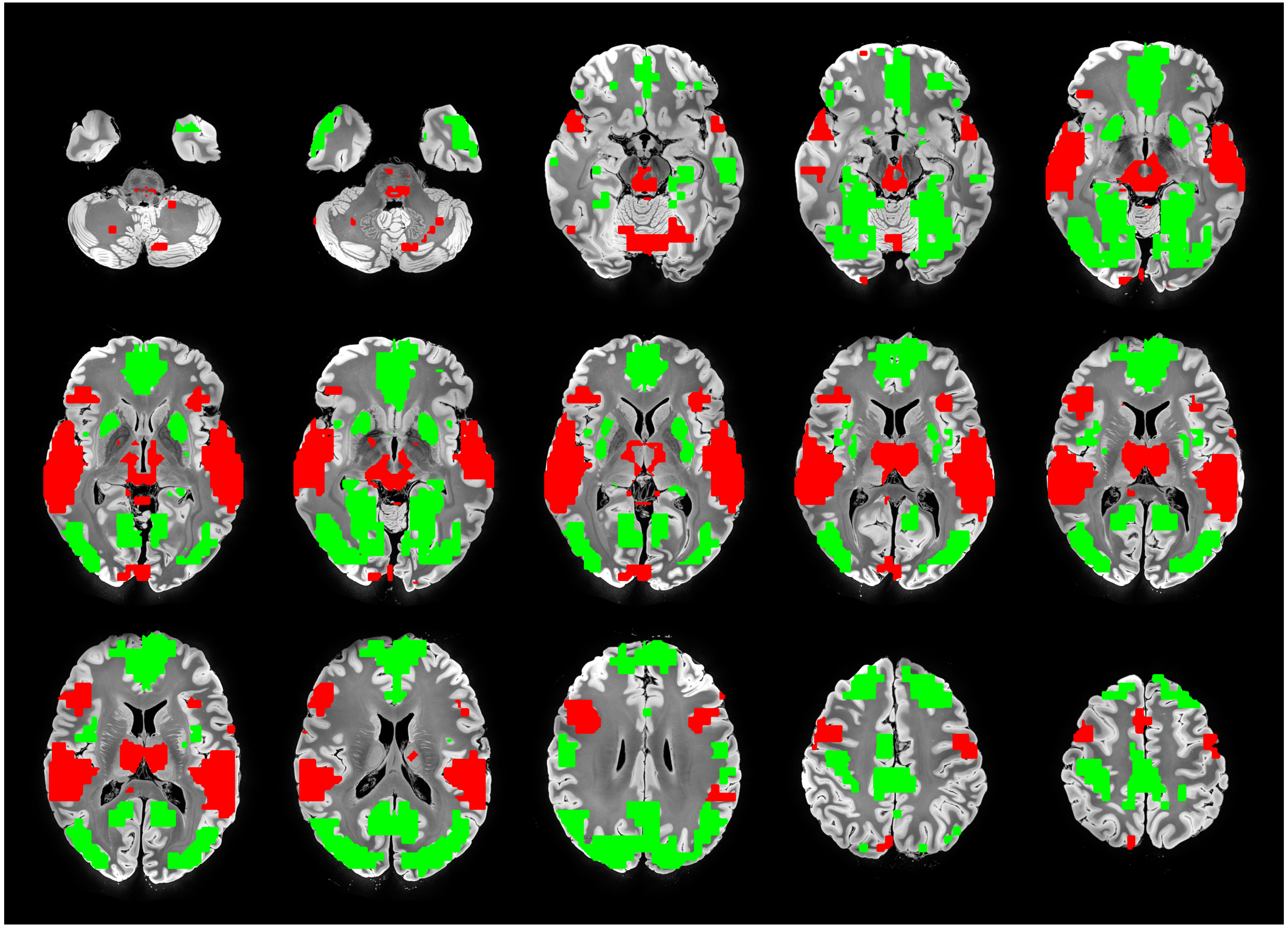

Common activation across all tasks    Common deactivation across all tasks

+6 sec

### Auditory Conjunction

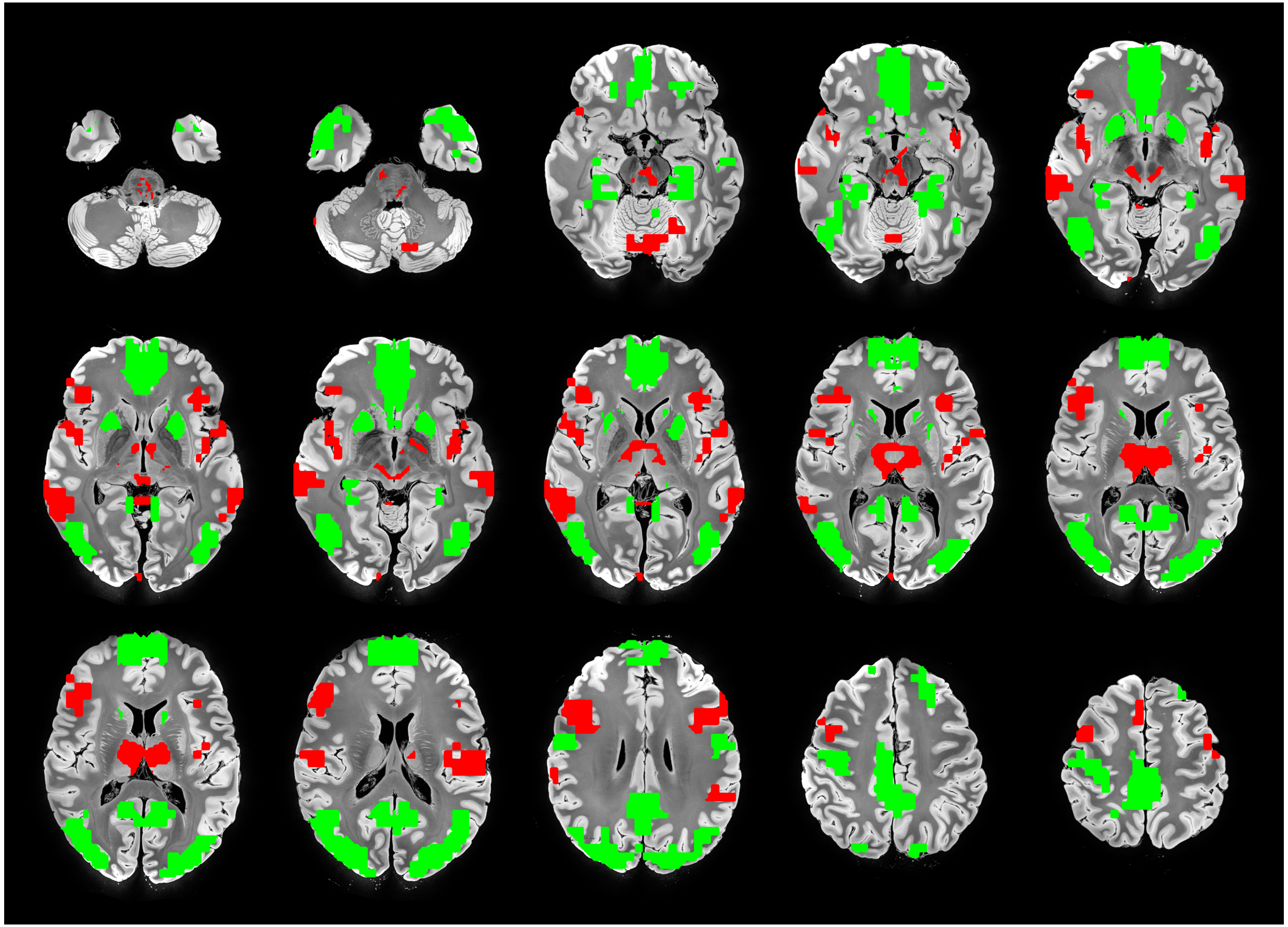

Common activation across all tasks

Common deactivation across all tasks

+8 sec

### Taste Conjunction

### Taste Conjunction

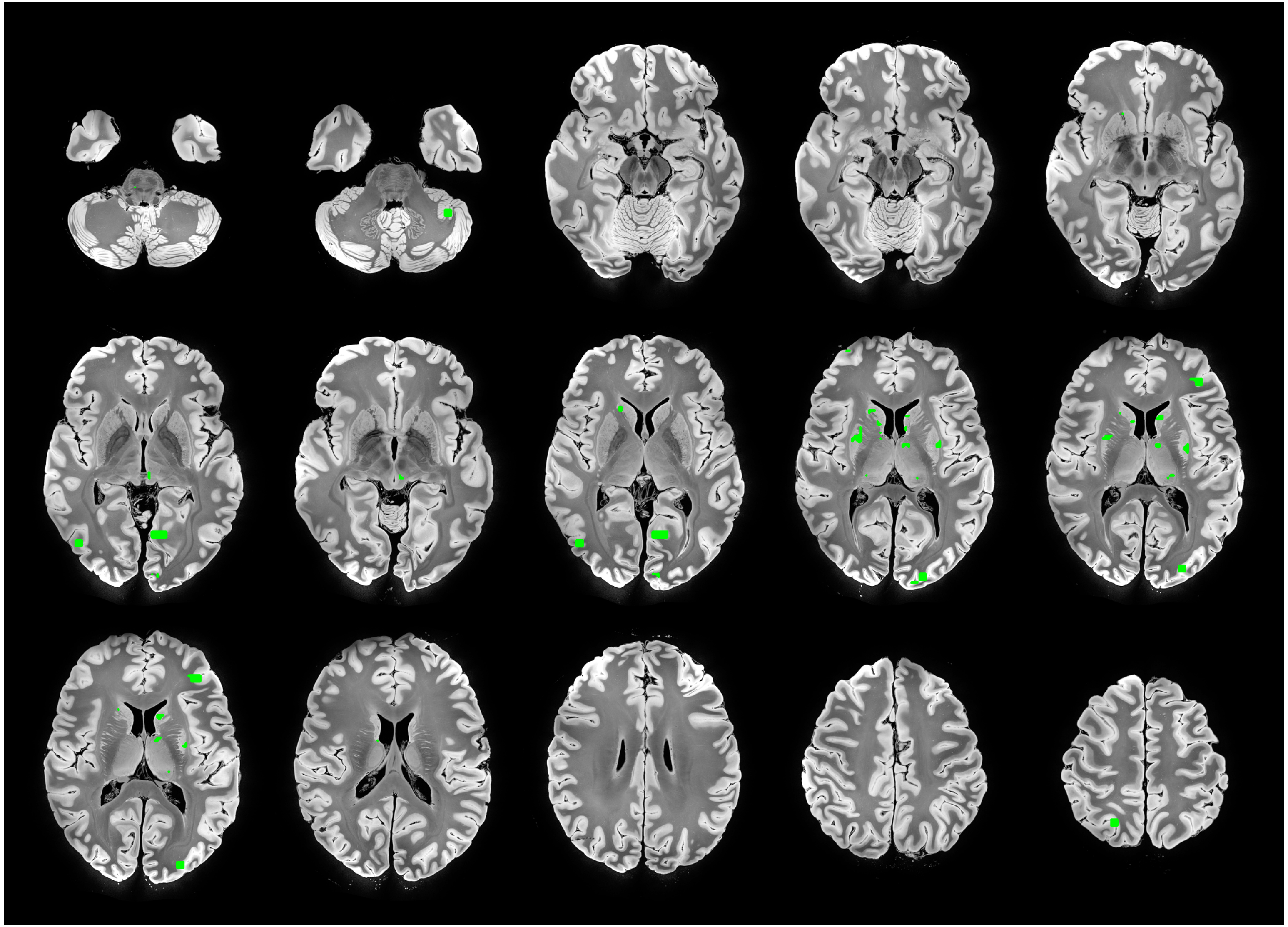

Common activation across all tasks

Common deactivation across all tasks

-4 sec

### Taste Conjunction

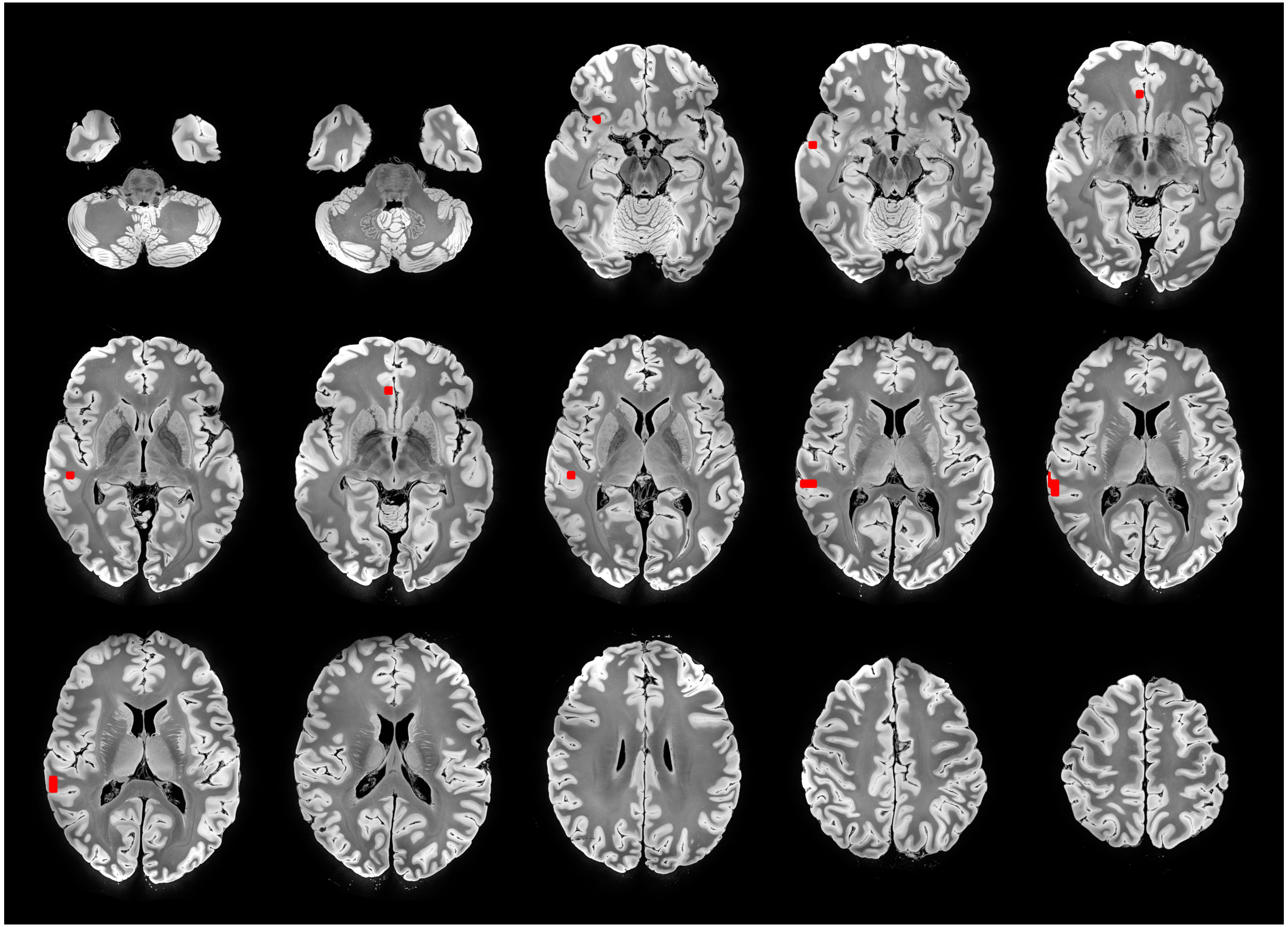

Common activation across all tasks
Common deactivation across all tasks

-2 sec

### Taste Conjunction

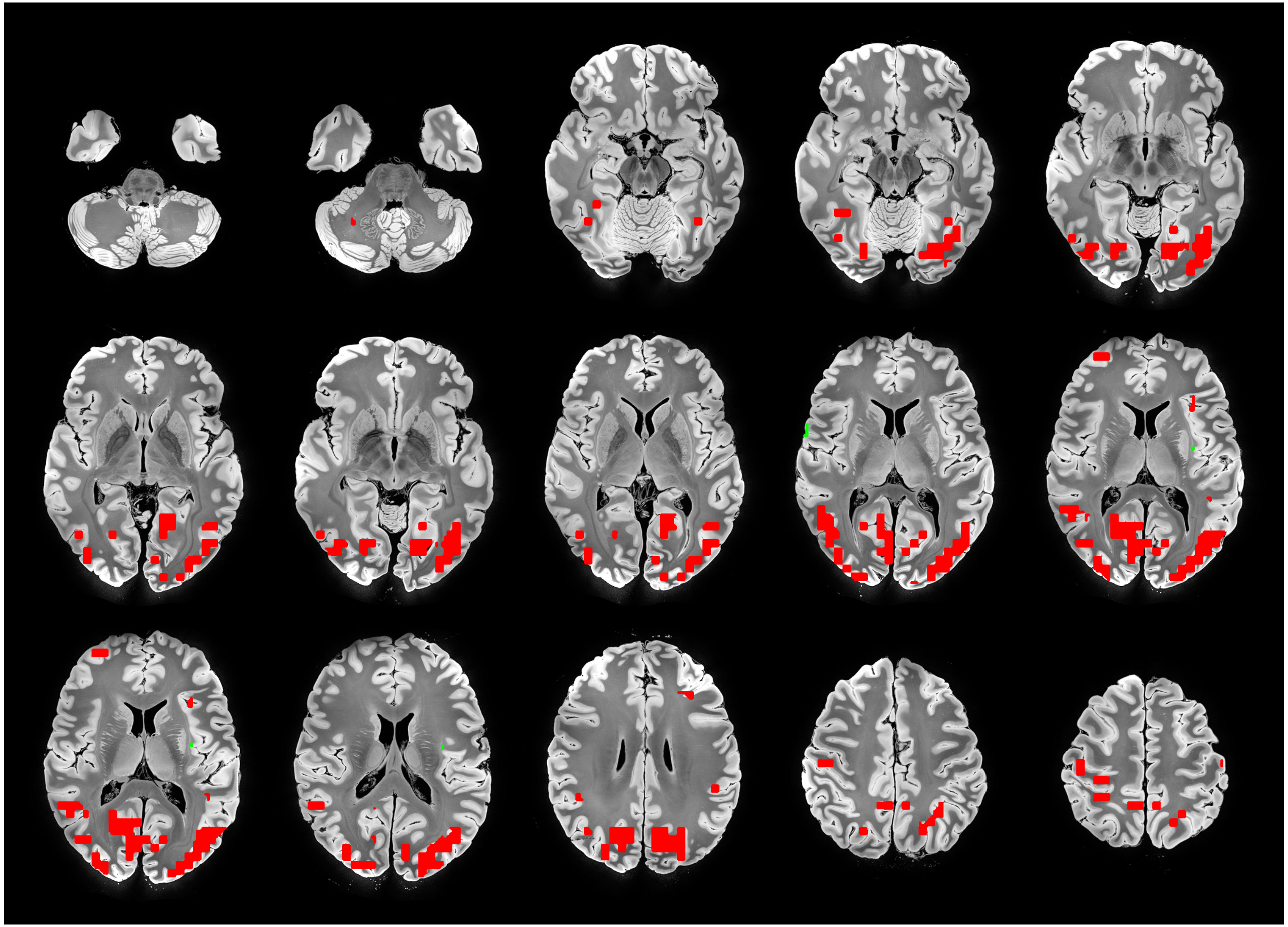

Common activation across all tasks

Common deactivation across all tasks

0 sec

### Taste Conjunction

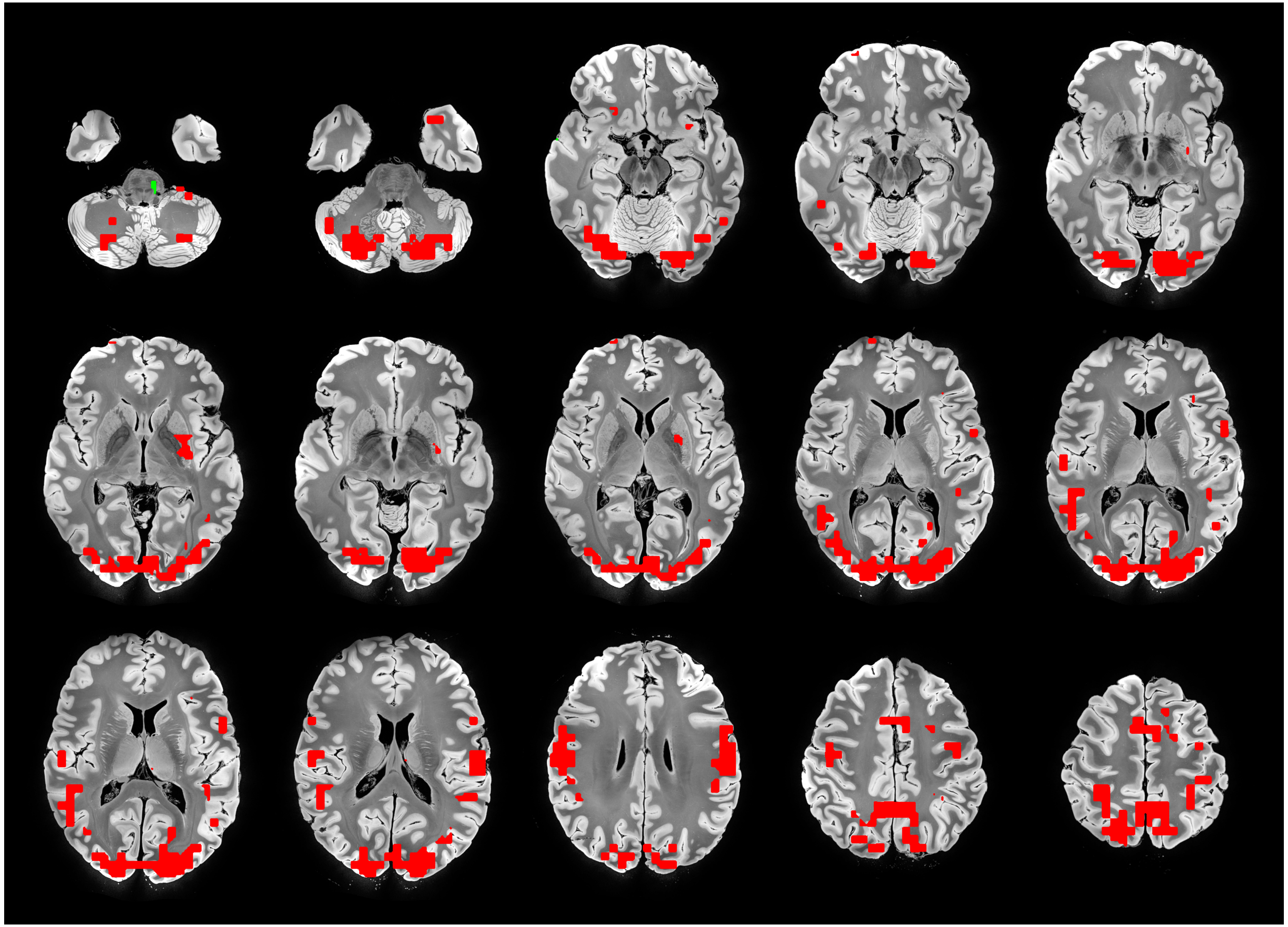

Common activation across all tasks

Common deactivation across all tasks

+2 sec

### Taste Conjunction

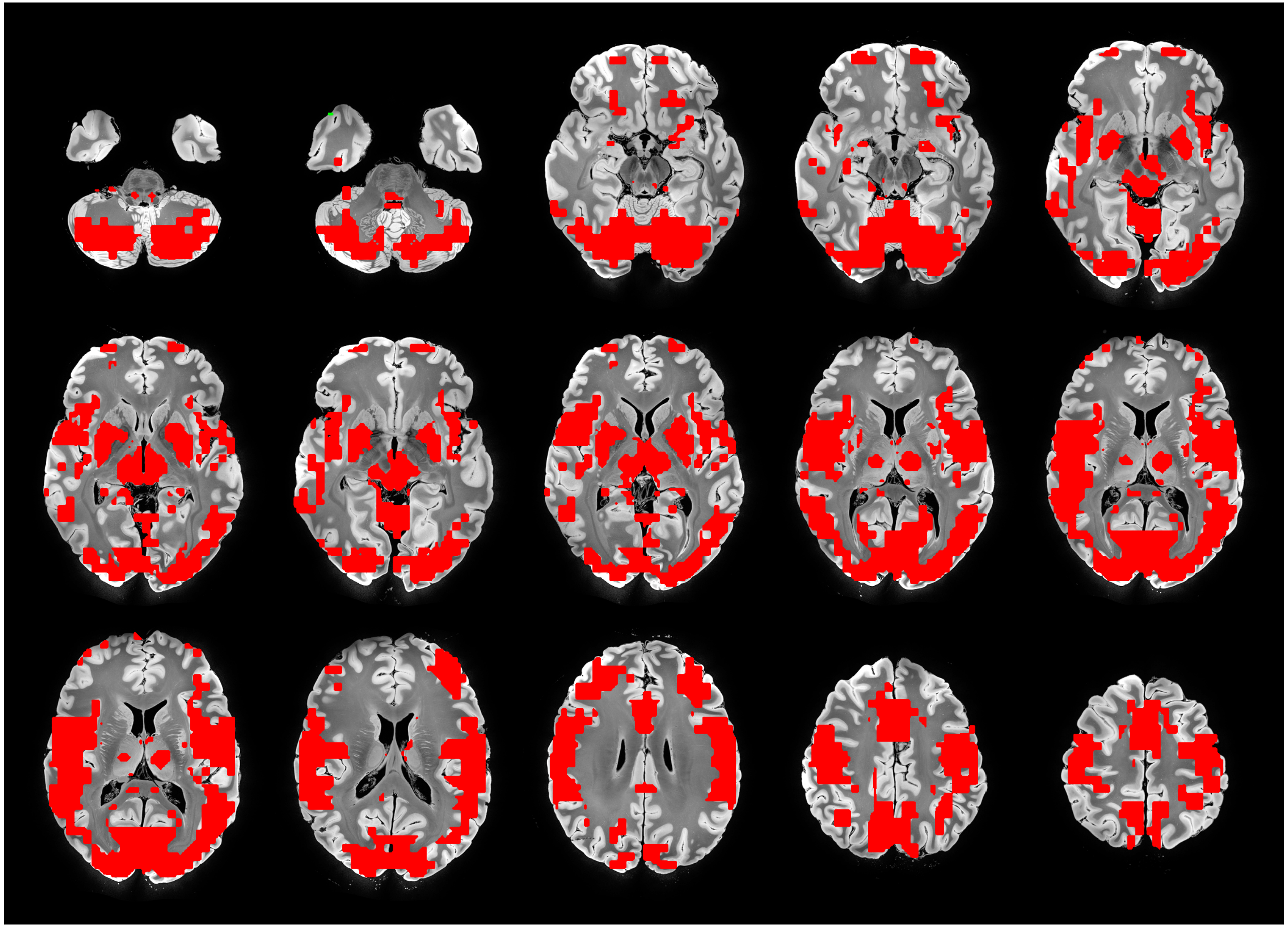

Common activation across all tasks

Common deactivation across all tasks

+4 sec

### Taste Conjunction

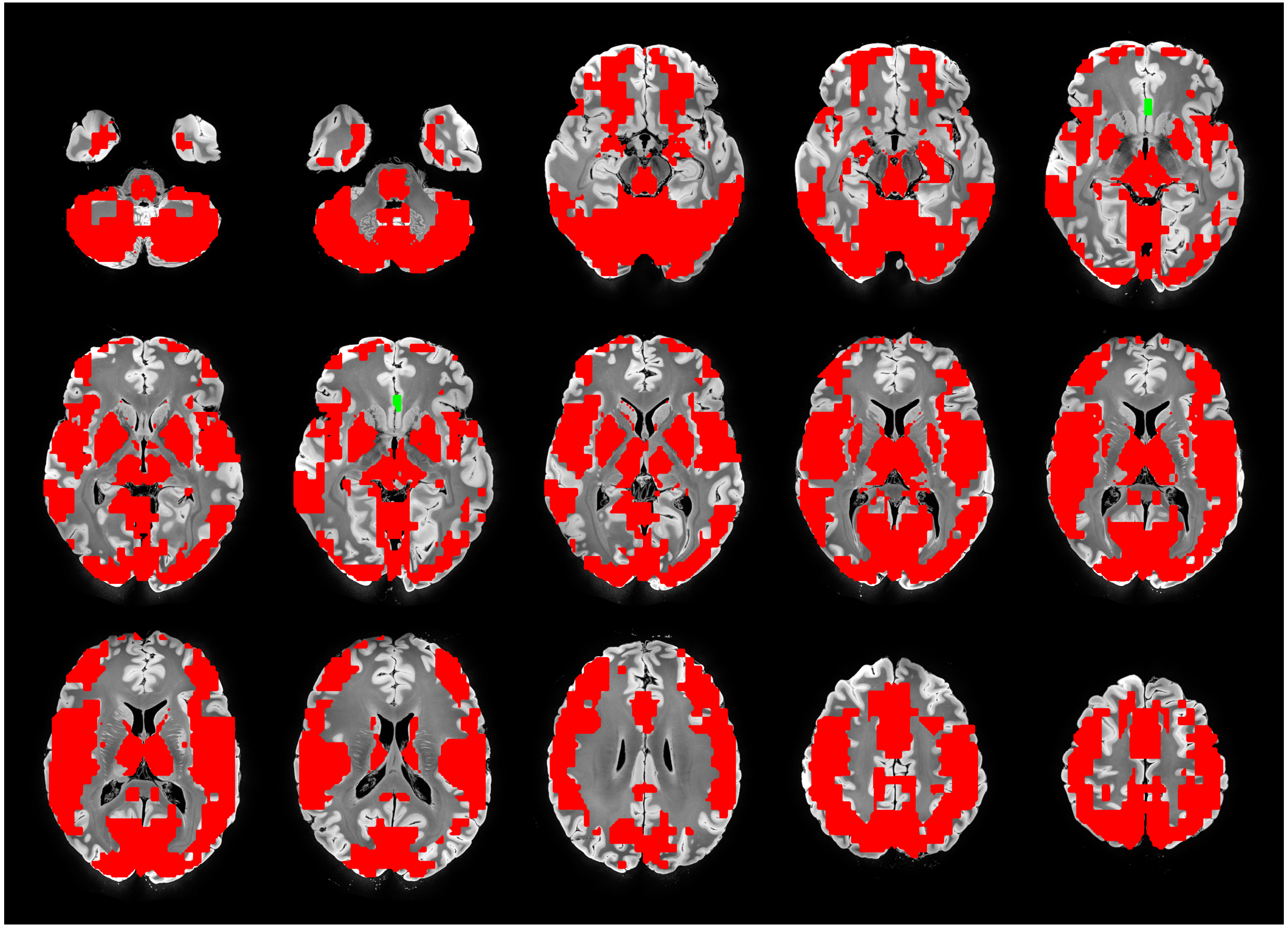

Common activation across all tasks

Common deactivation across all tasks

+6 sec

### Taste Conjunction

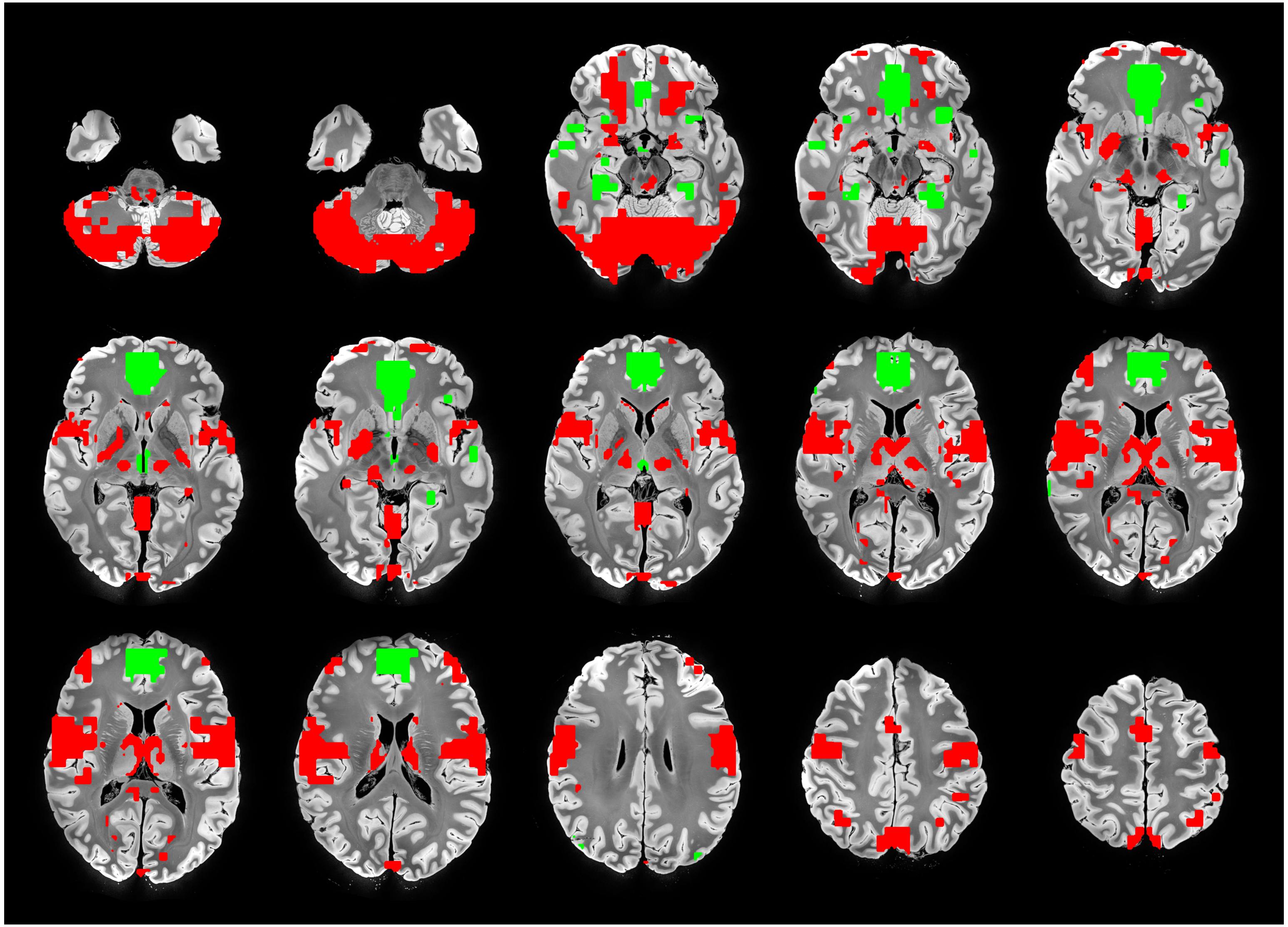

Common activation across all tasks

Common deactivation across all tasks

+8 sec

### Tactile Significant Changes

### Tactile Significant Changes

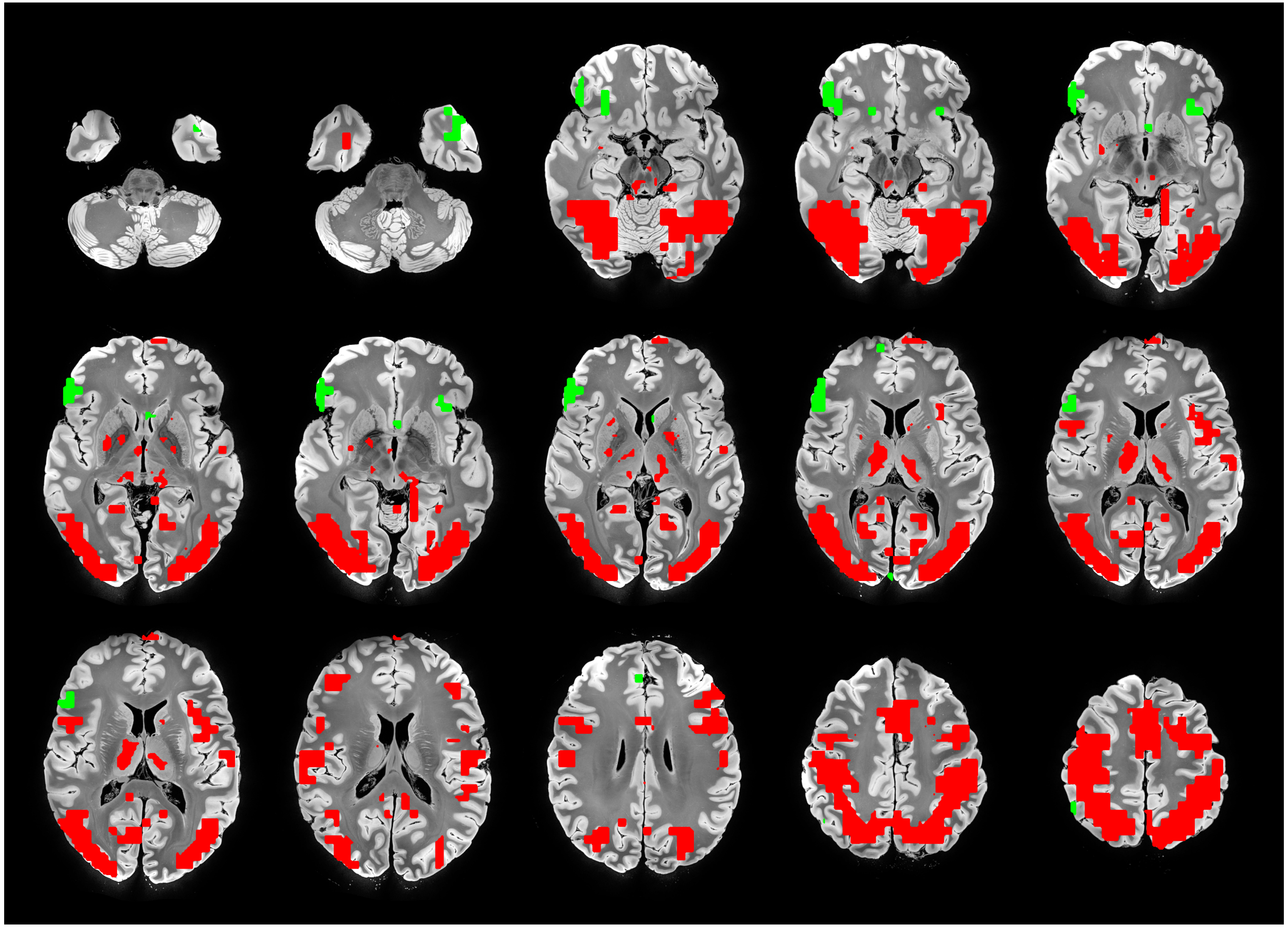

Activation Deactivation

-4 sec

### Tactile Significant Changes

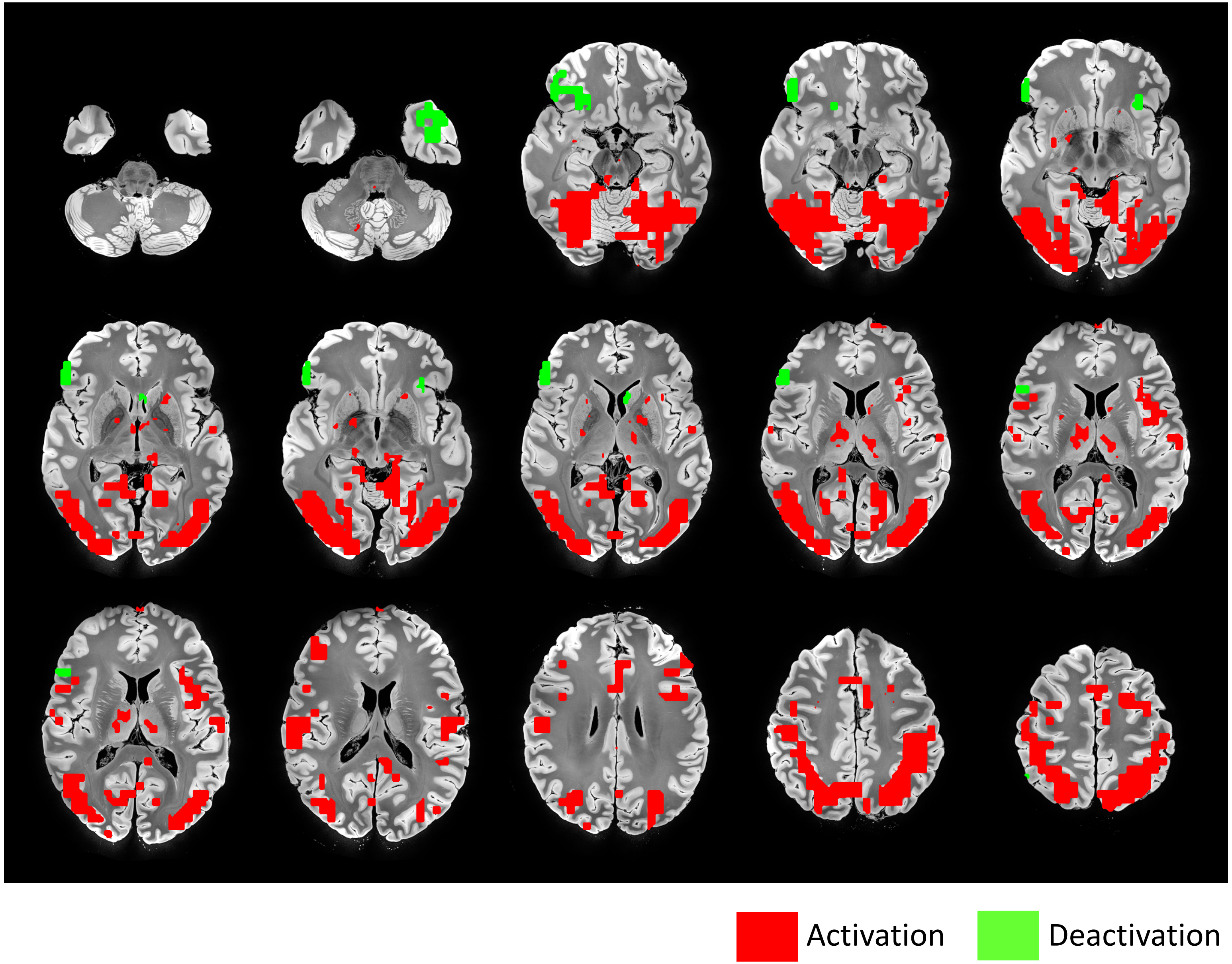



-2 sec

### Tactile Significant Changes

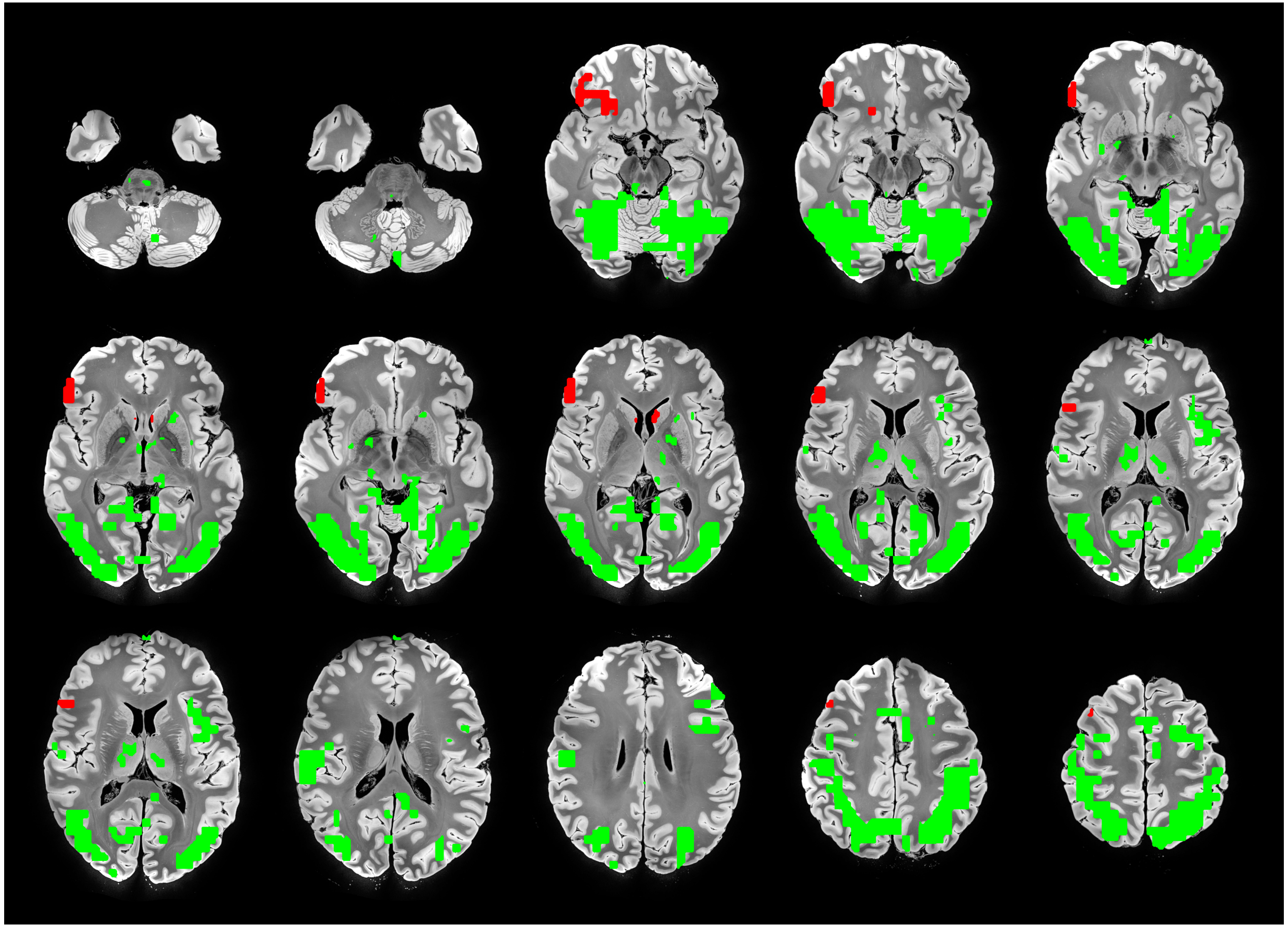

### Tactile Significant Changes

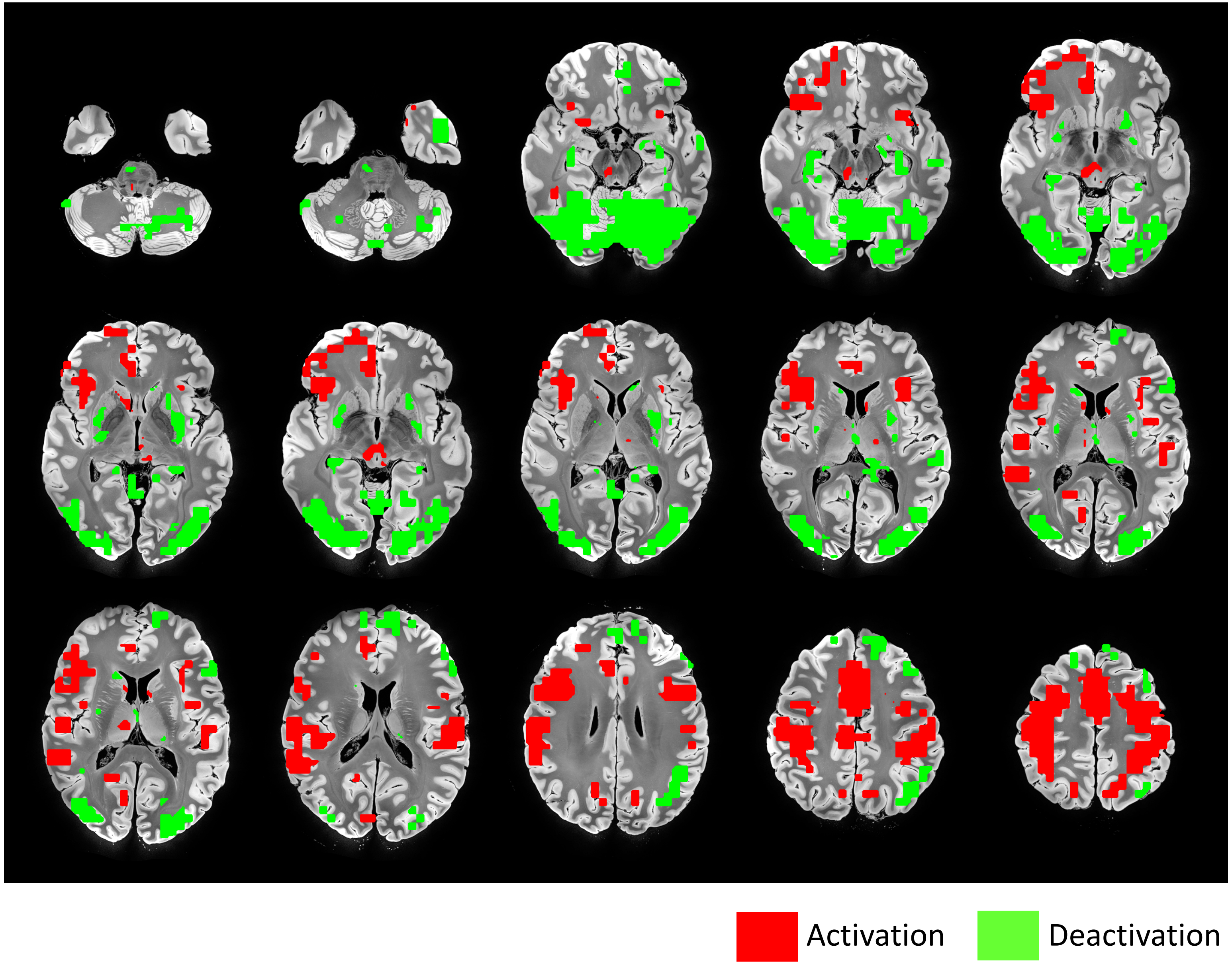


+2 sec

### Tactile Significant Changes

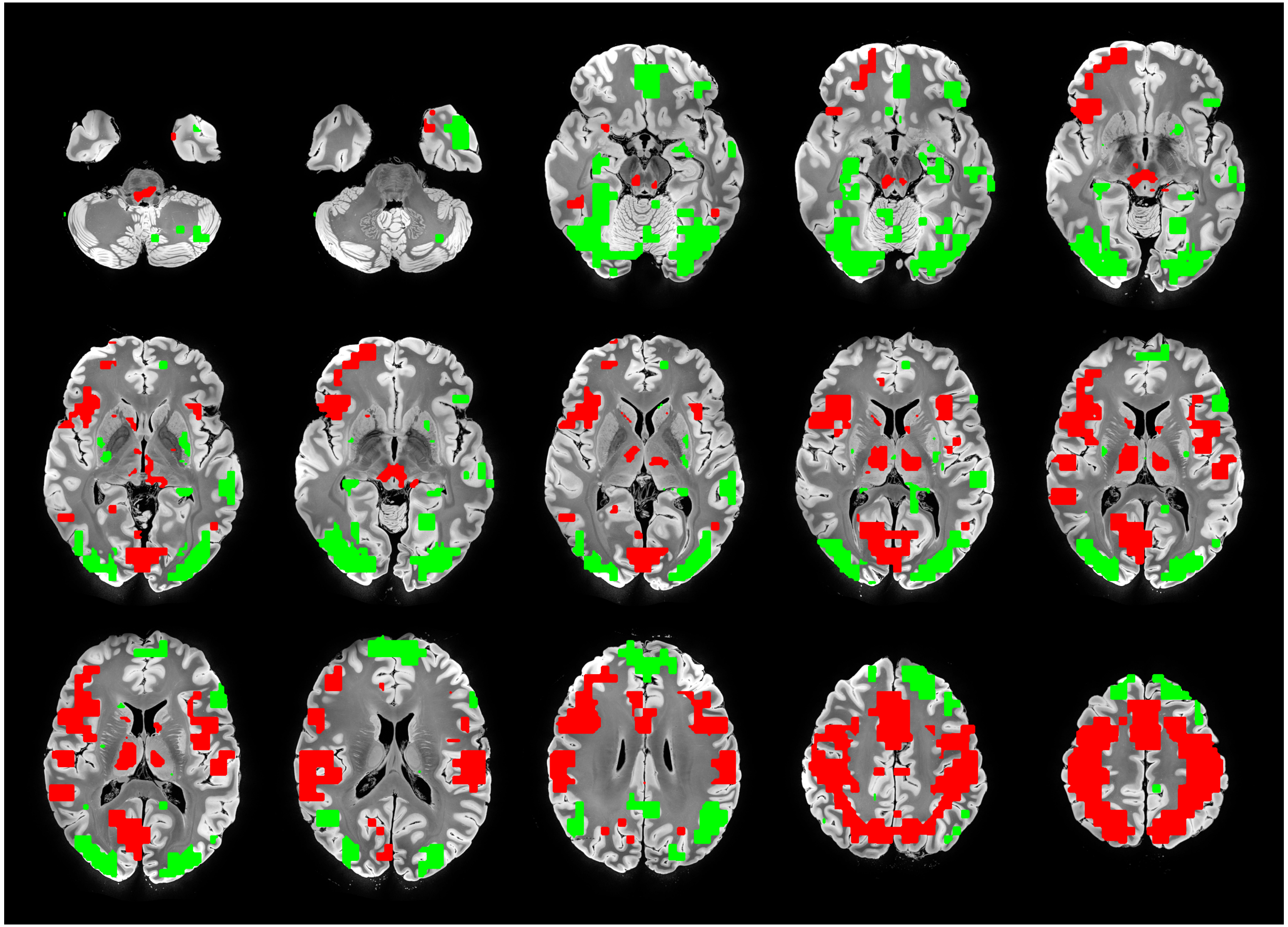

Activation Deactivation

+4 sec

### Tactile Significant Changes

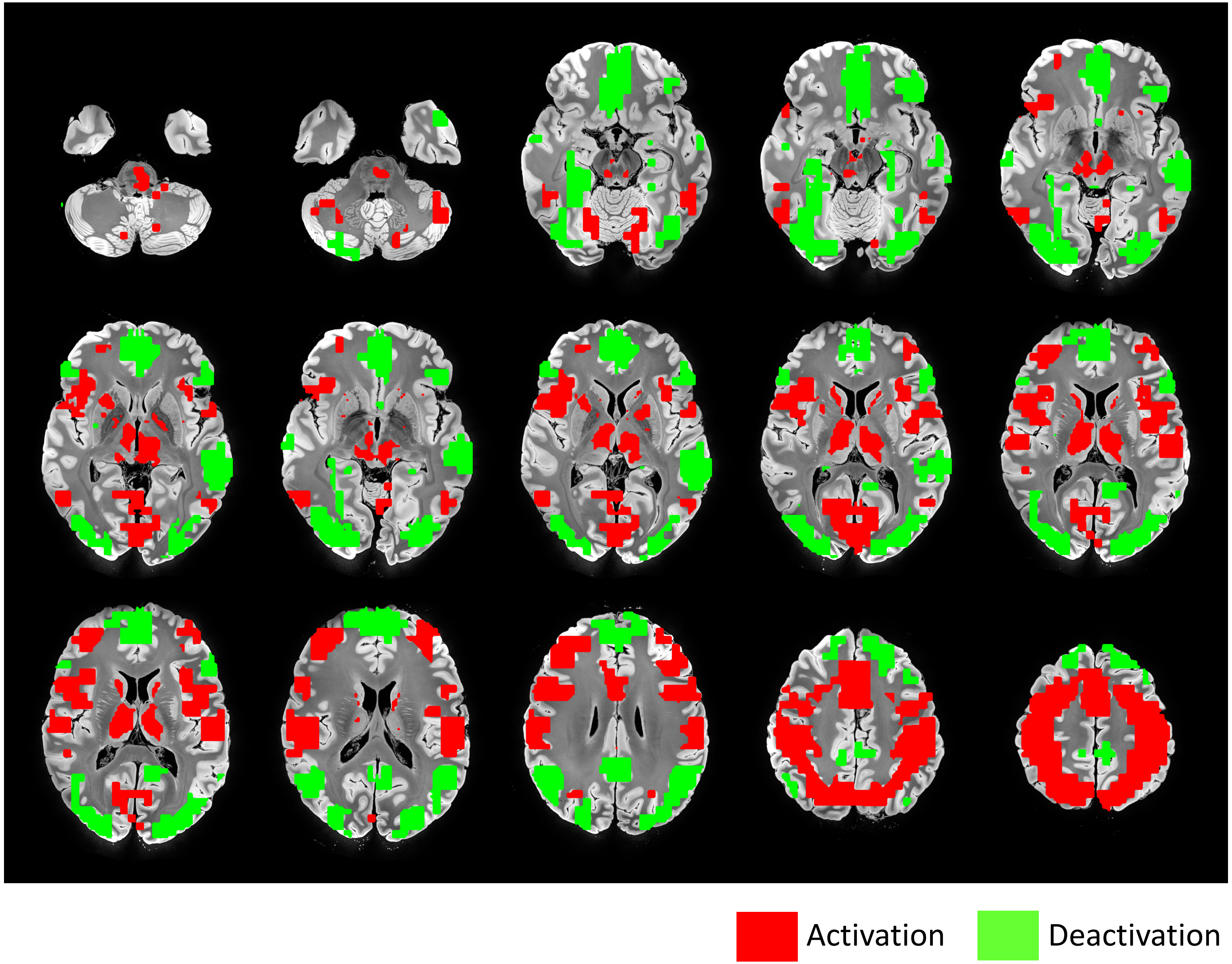


+6 sec

### Tactile Significant Changes

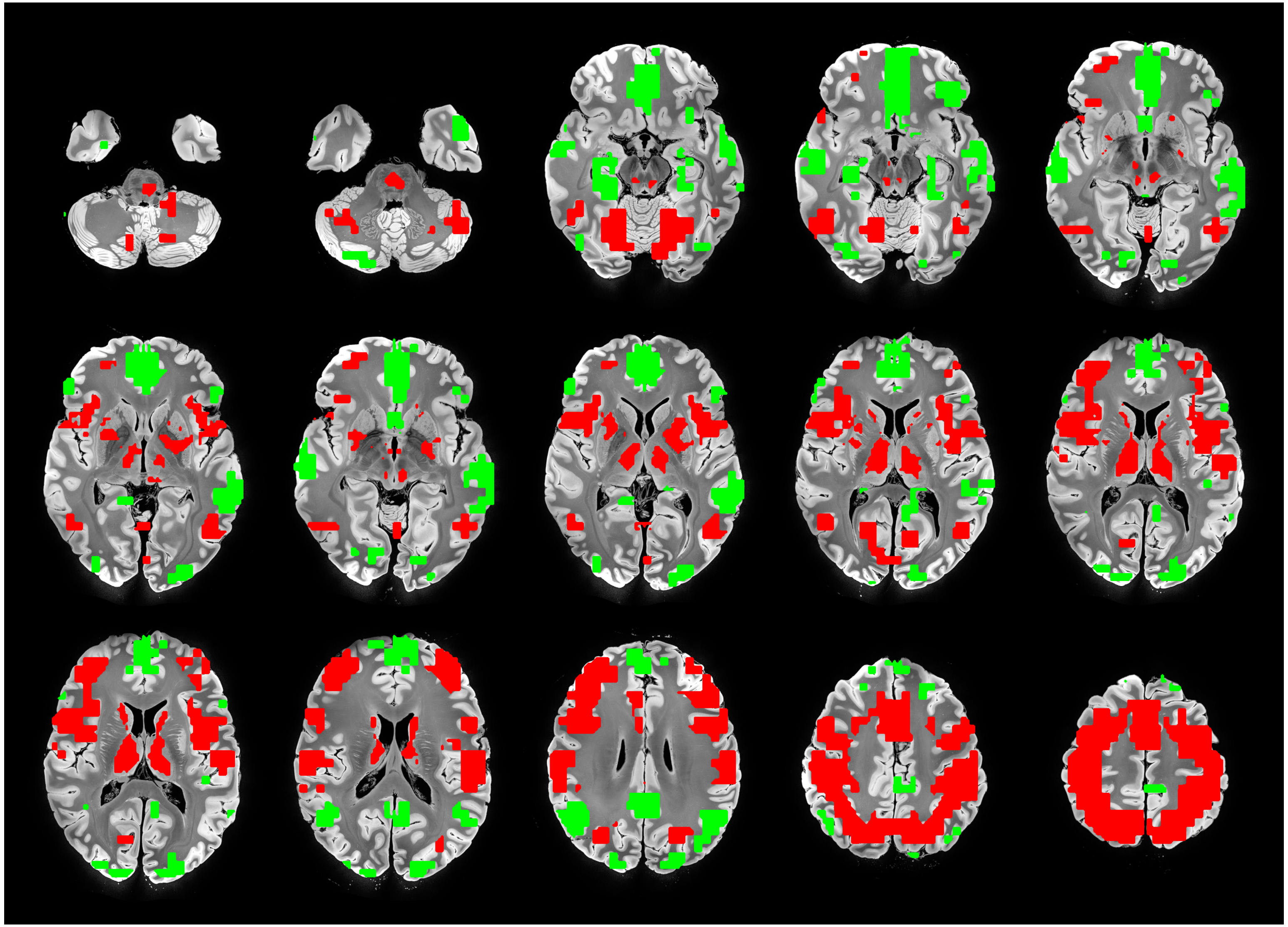



+8 sec
