## Supplementary Presentation S3 for "Shared subcortical arousal systems across sensory modalities during transient modulation of attention"

### Exclusive Disjunction for Each Modality

### Visual Disjunction

### Visual Disjunction

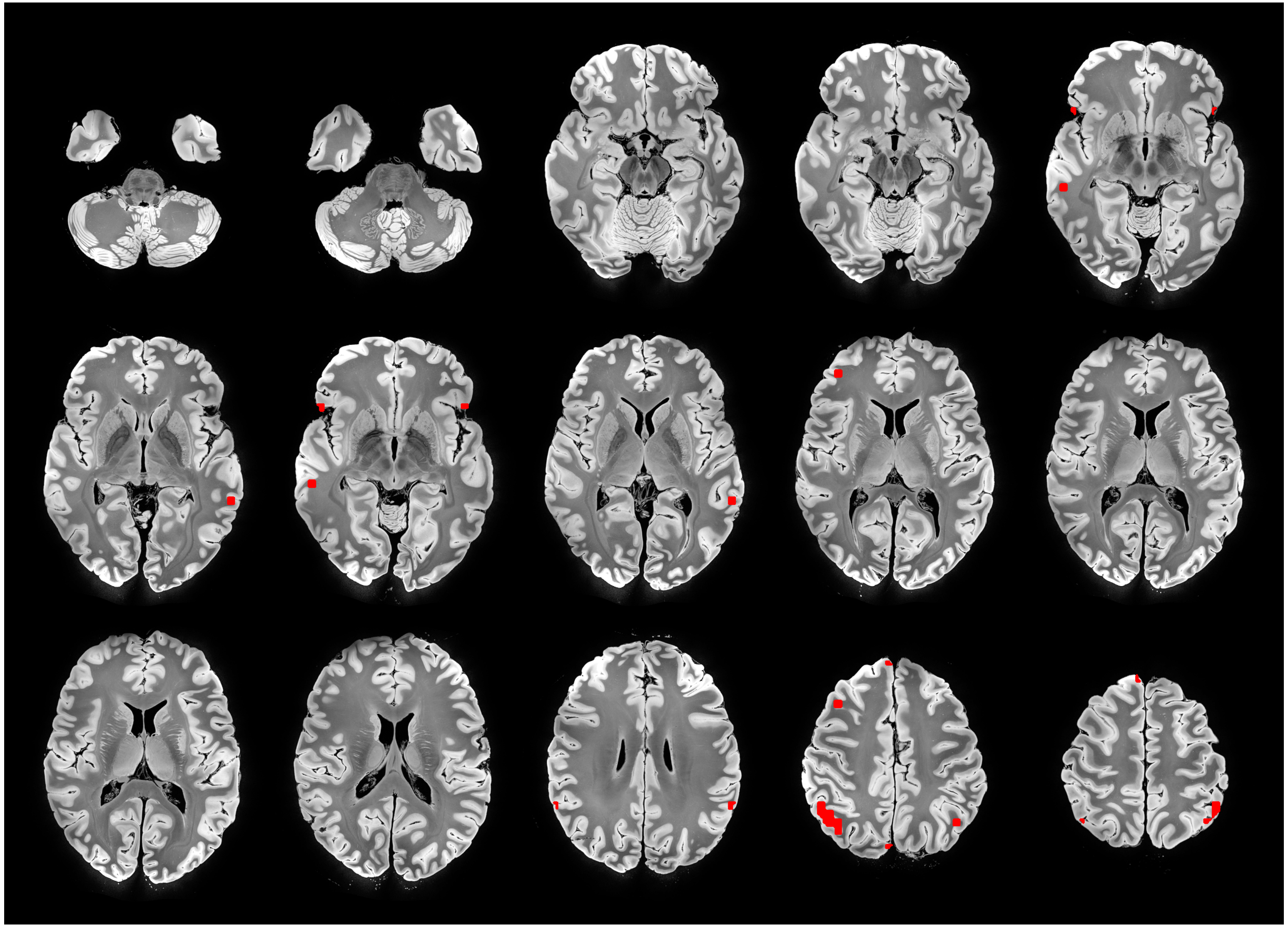

Common activation across all tasks

Common deactivation across all tasks

-4 sec

### Visual Disjunction

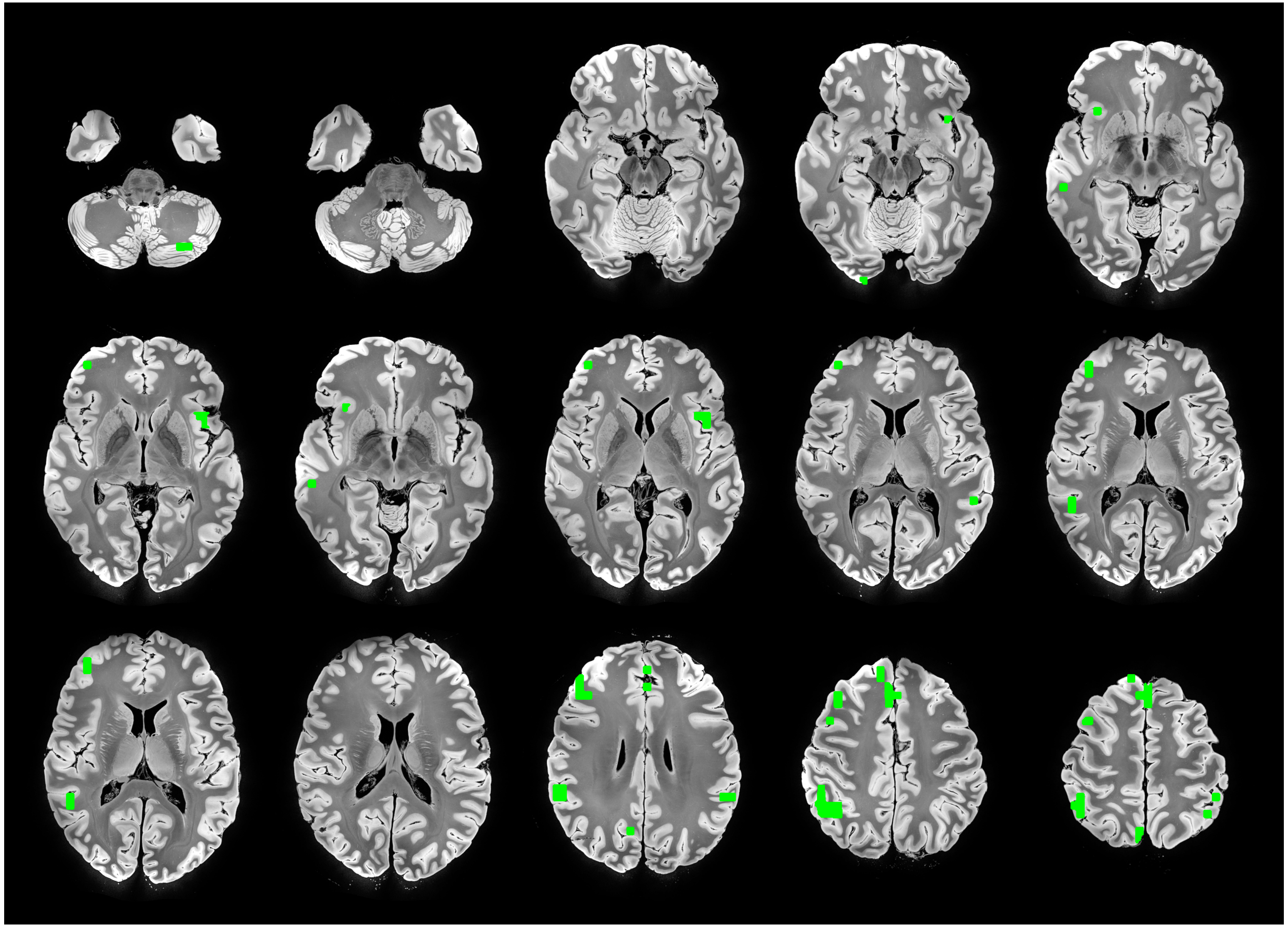

Common activation across all tasks    Common deactivation across all tasks

-2 sec

### Visual Disjunction

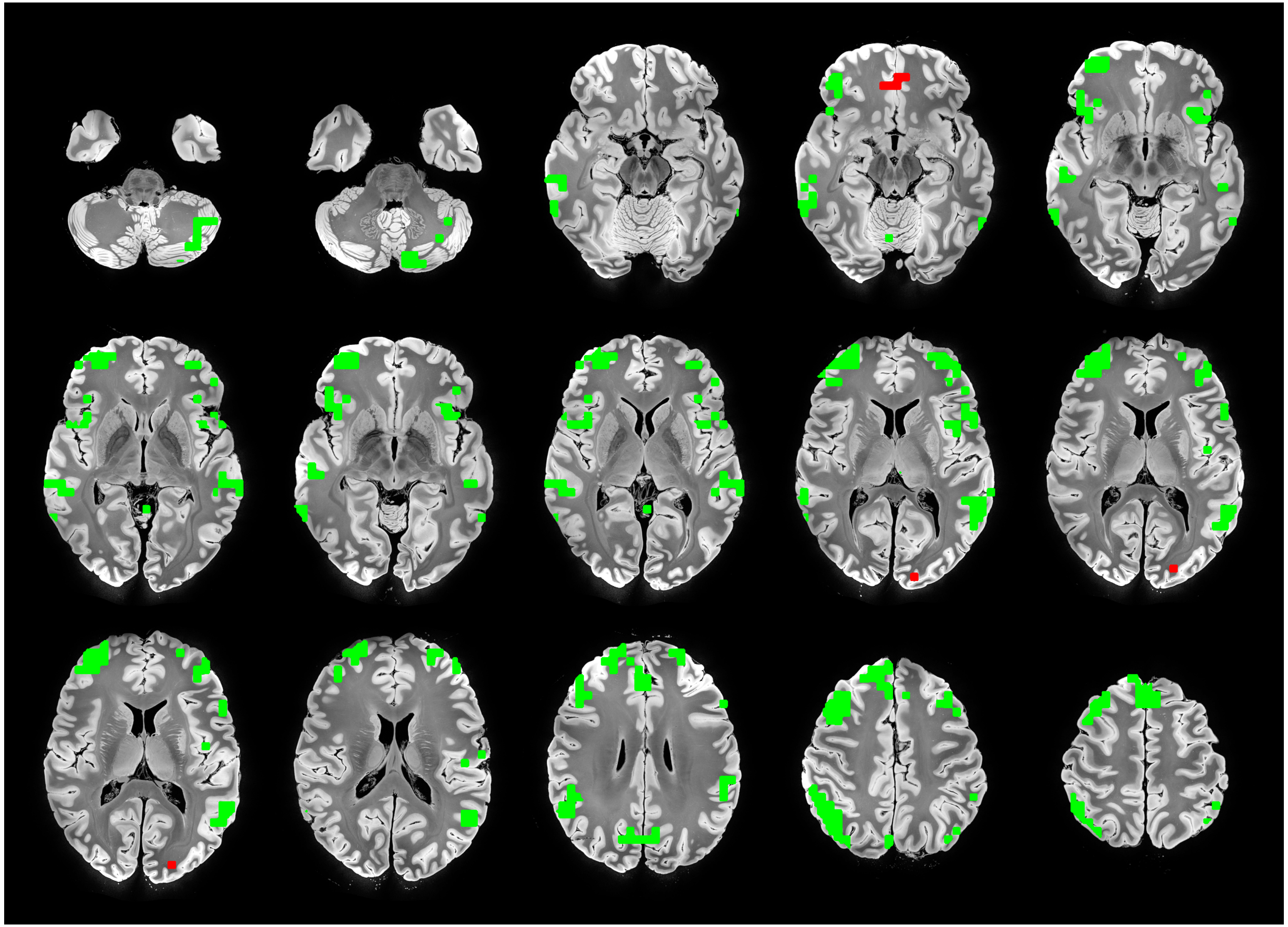

Common activation across all tasks

Common deactivation across all tasks

0 sec

### Visual Disjunction

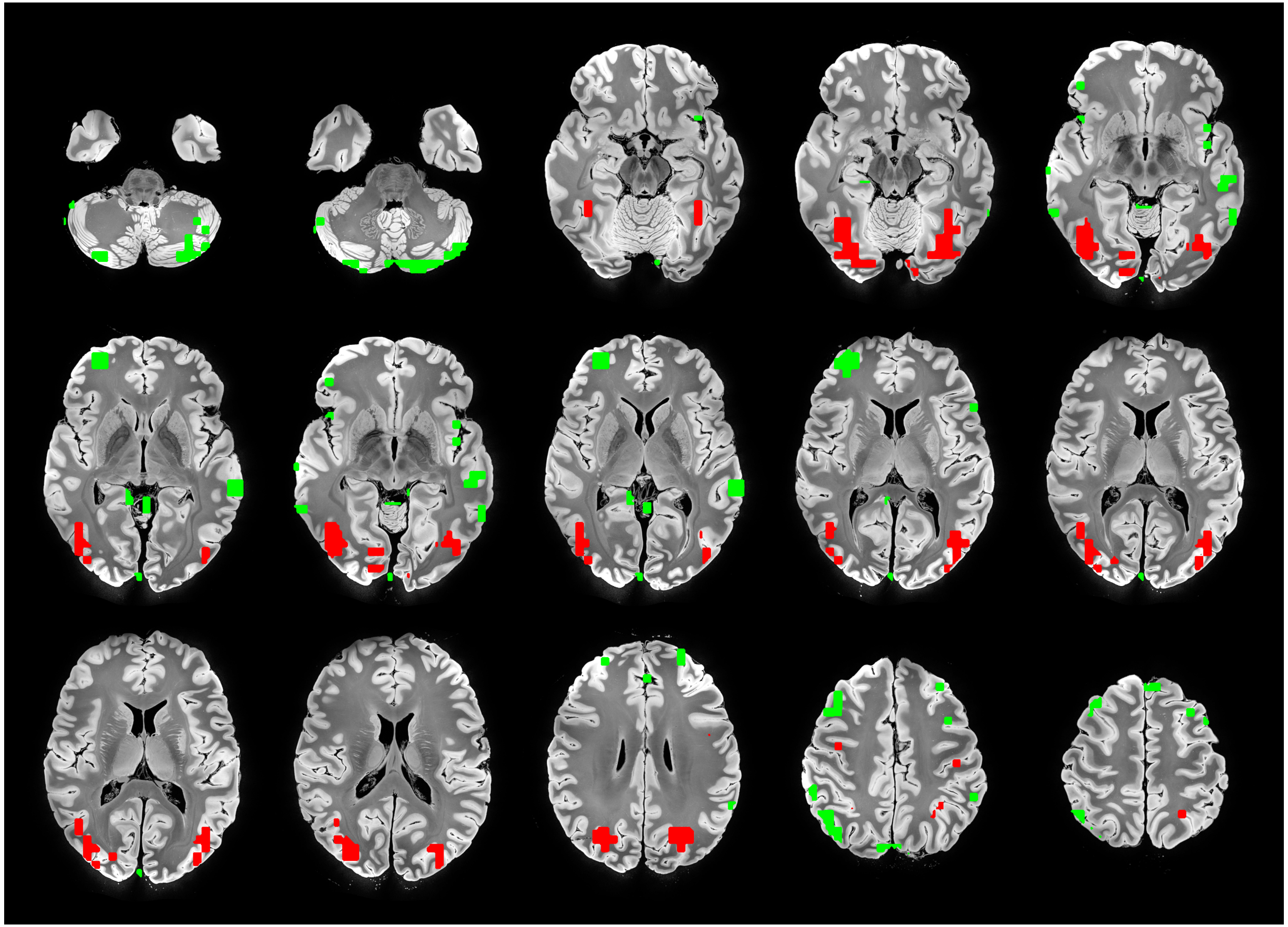

Common activation across all tasks
Common deactivation across all tasks

+2 sec

### Visual Disjunction

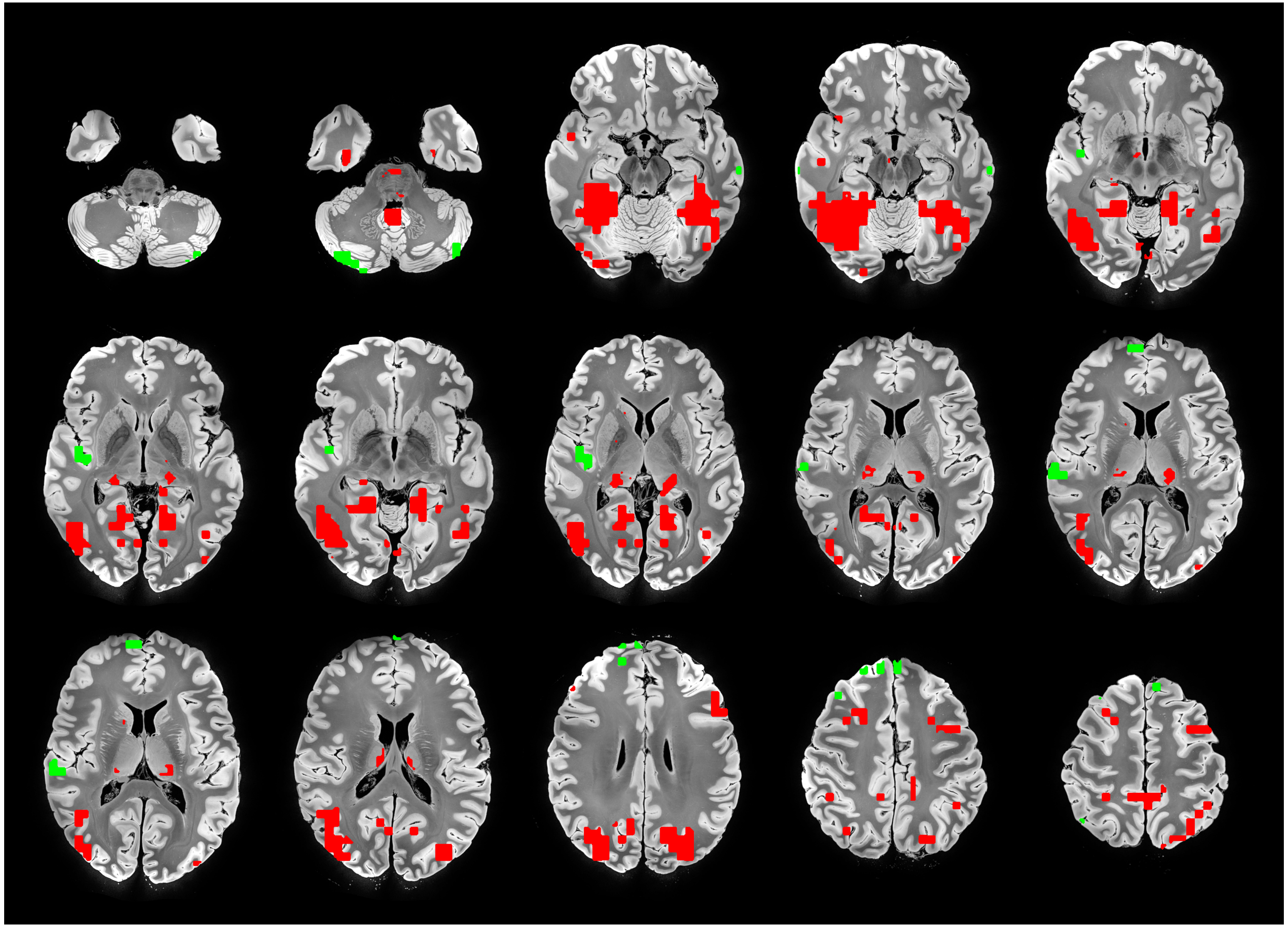



+4 sec

### Visual Disjunction

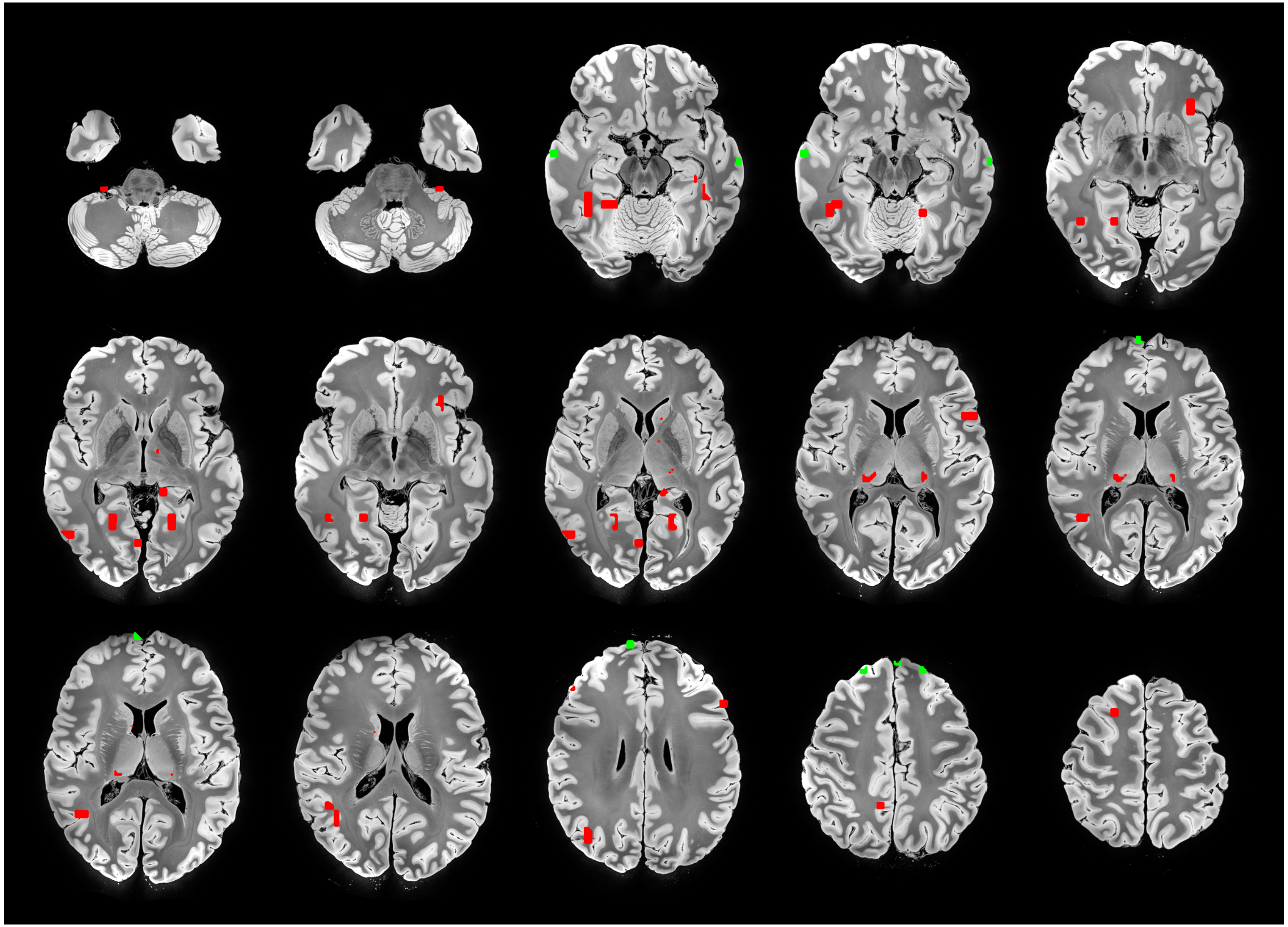

Common activation across all tasks

Common deactivation across all tasks

+6 sec

### Visual Disjunction

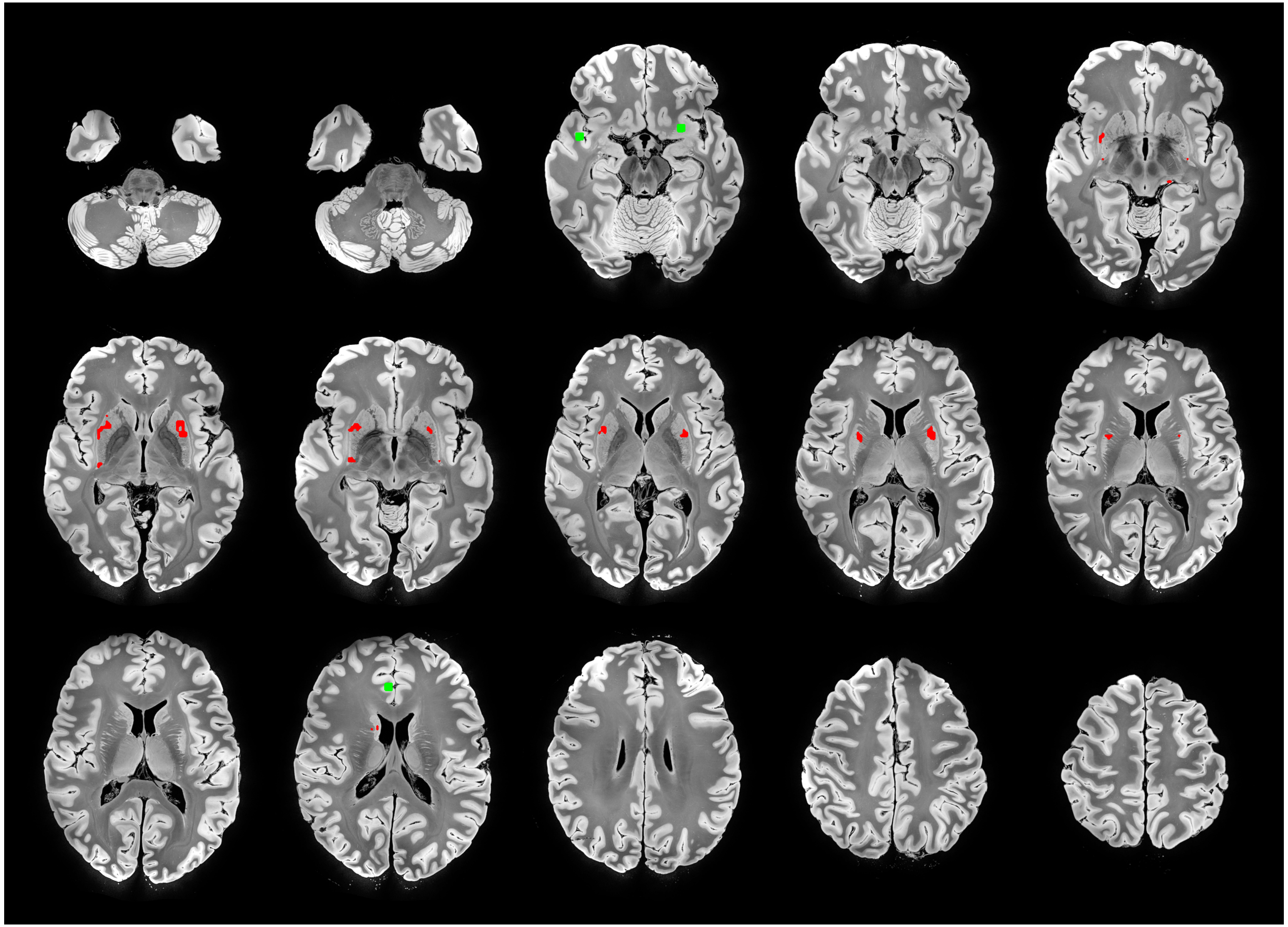

Common activation across all tasks

Common deactivation across all tasks

+8 sec

### Auditory Disjunction

### Auditory Disjunction

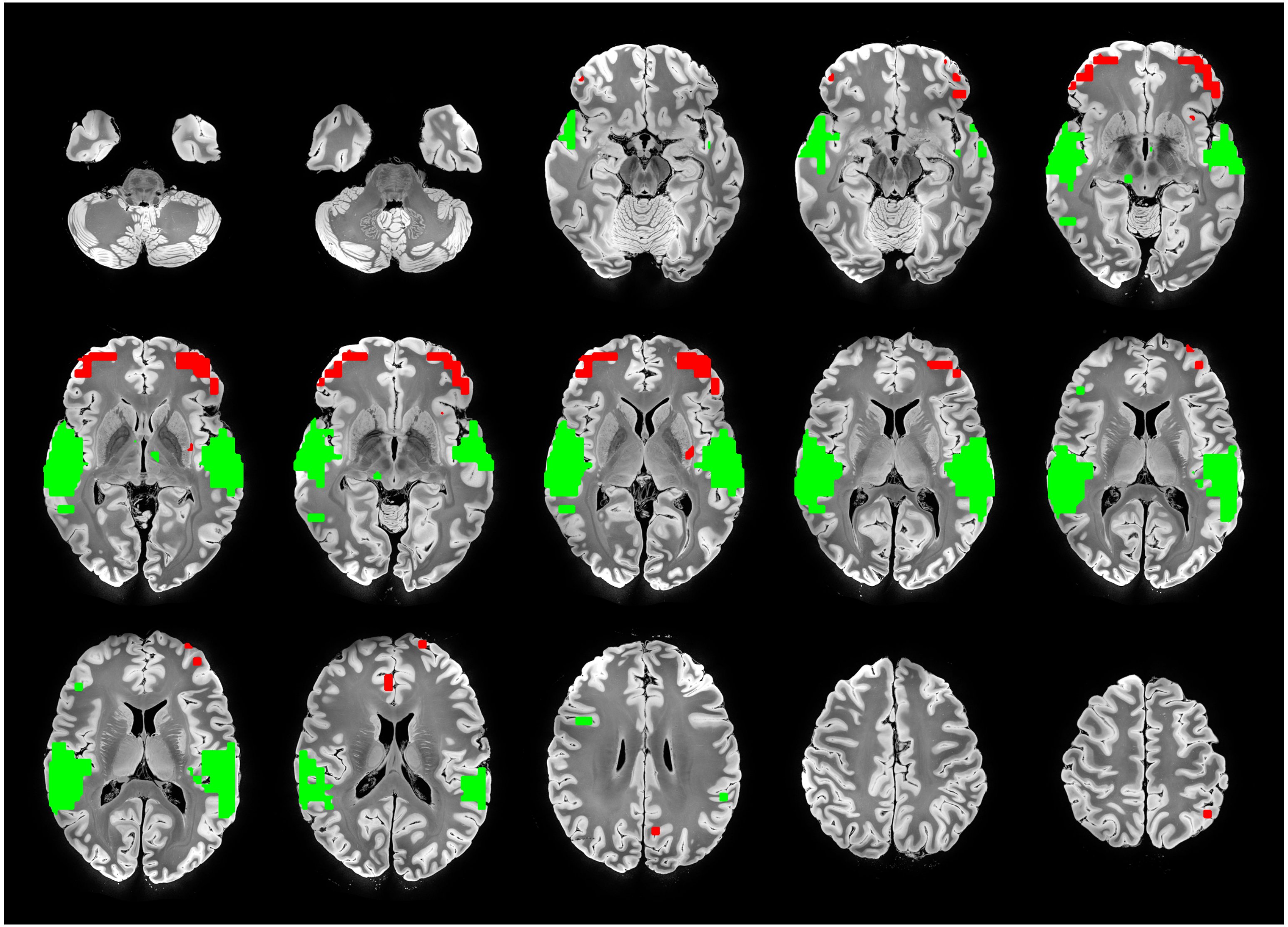

Common activation across all tasks

Common deactivation across all tasks

-4 sec

### Auditory Disjunction

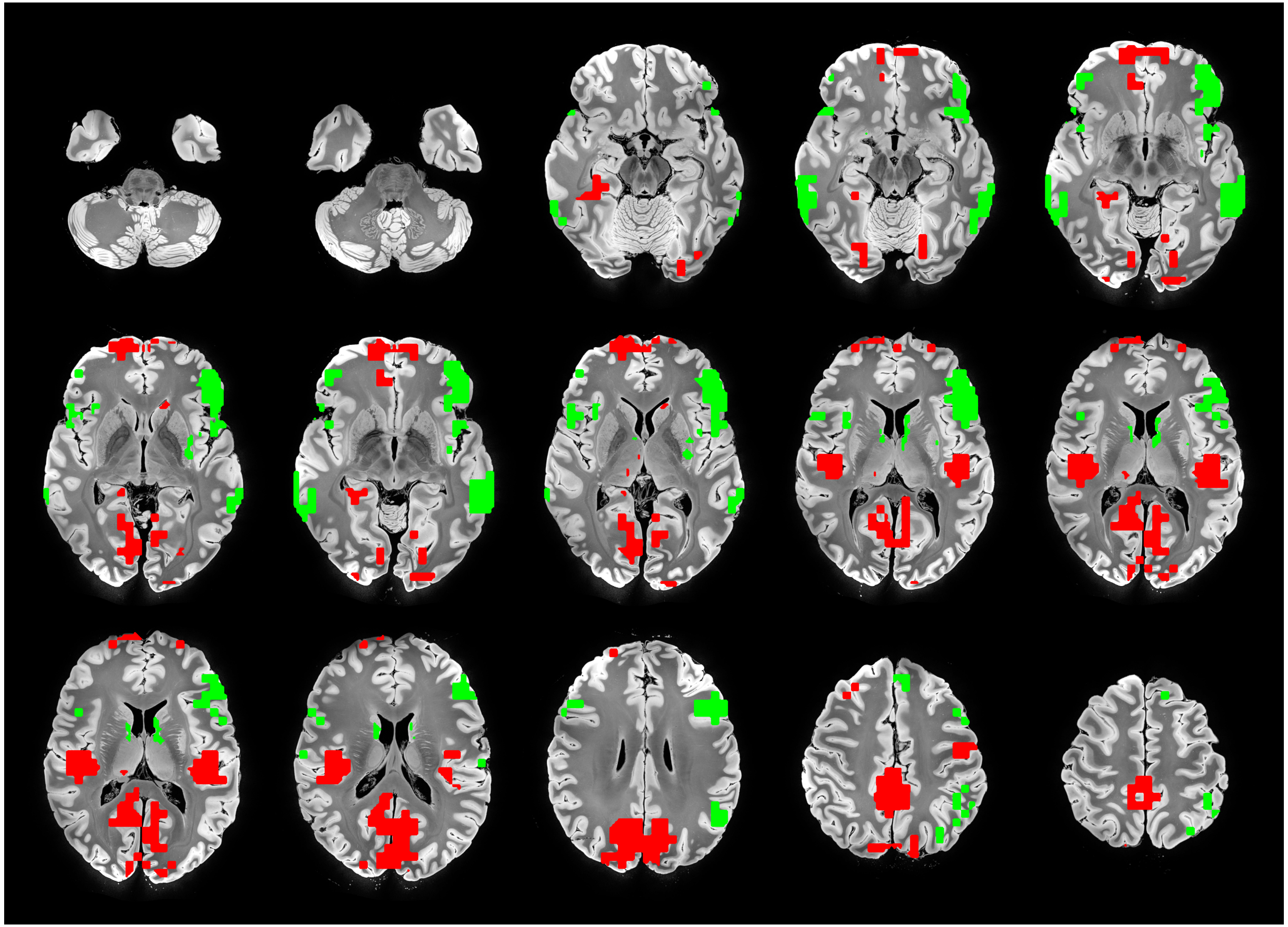

Common activation across all tasks    Common deactivation across all tasks

-2 sec

### Auditory Disjunction

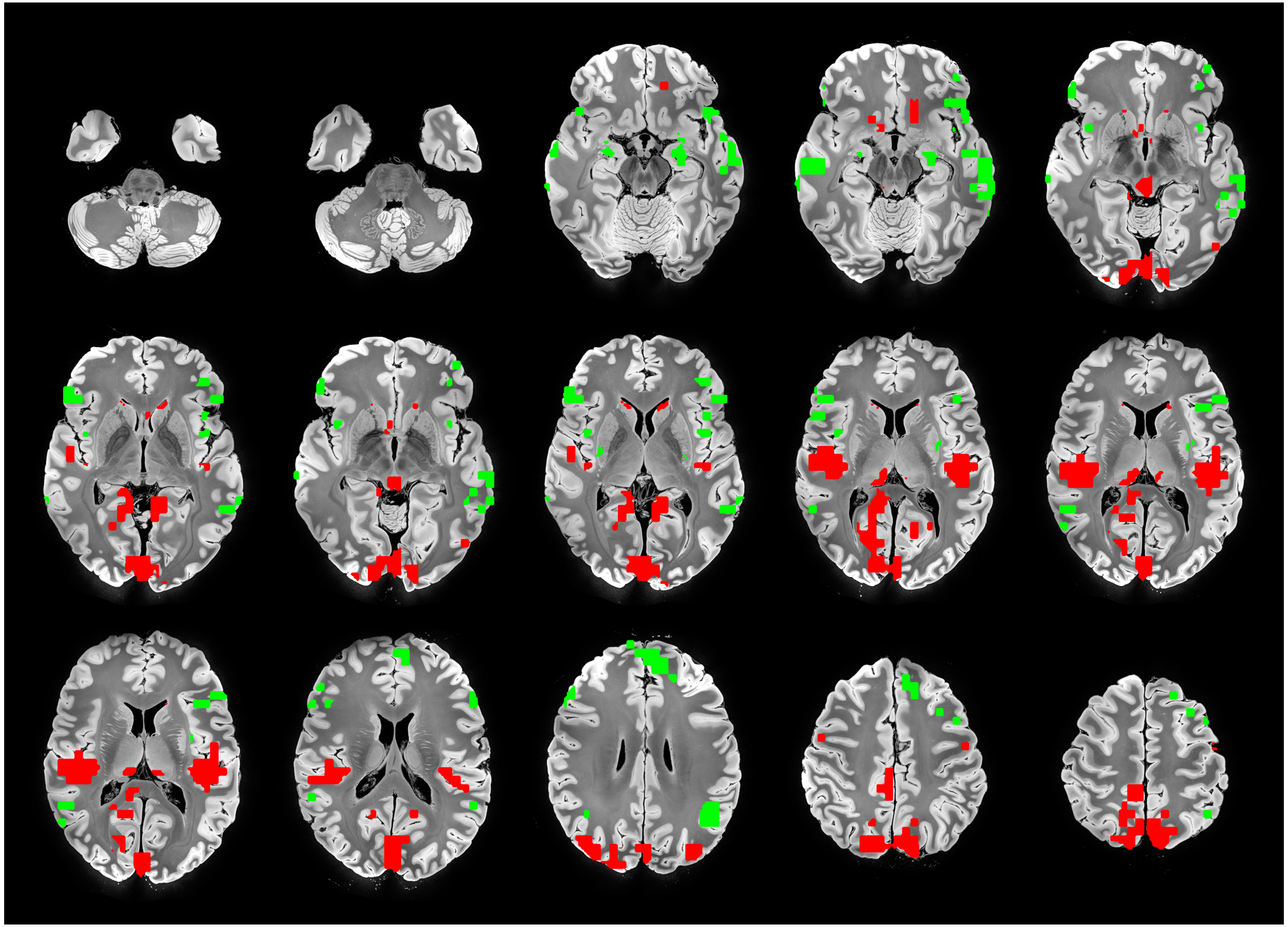

Common activation across all tasks

Common deactivation across all tasks

0 sec

### Auditory Disjunction

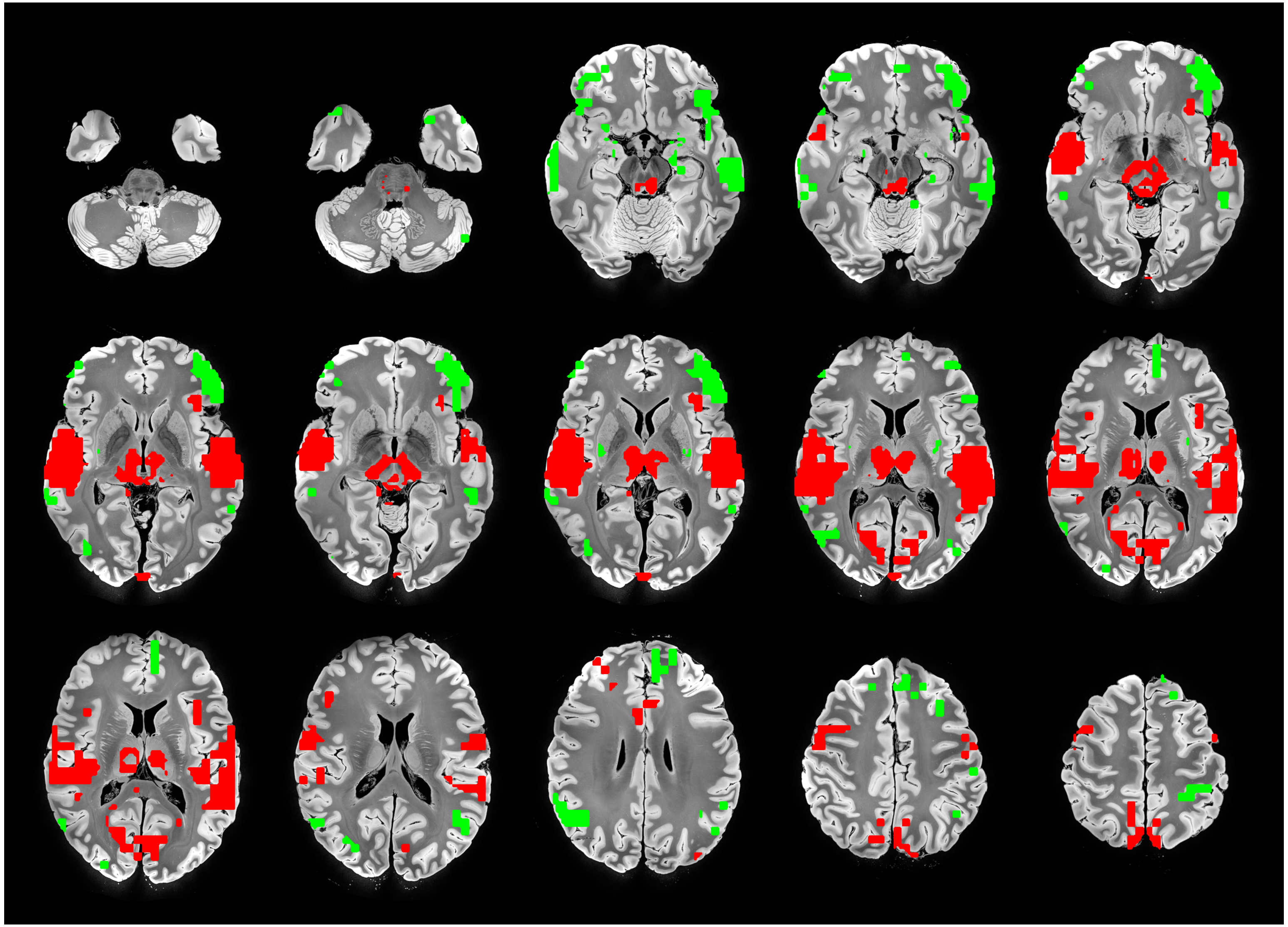

Common activation across all tasks

Common deactivation across all tasks

+2 sec

### Auditory Disjunction

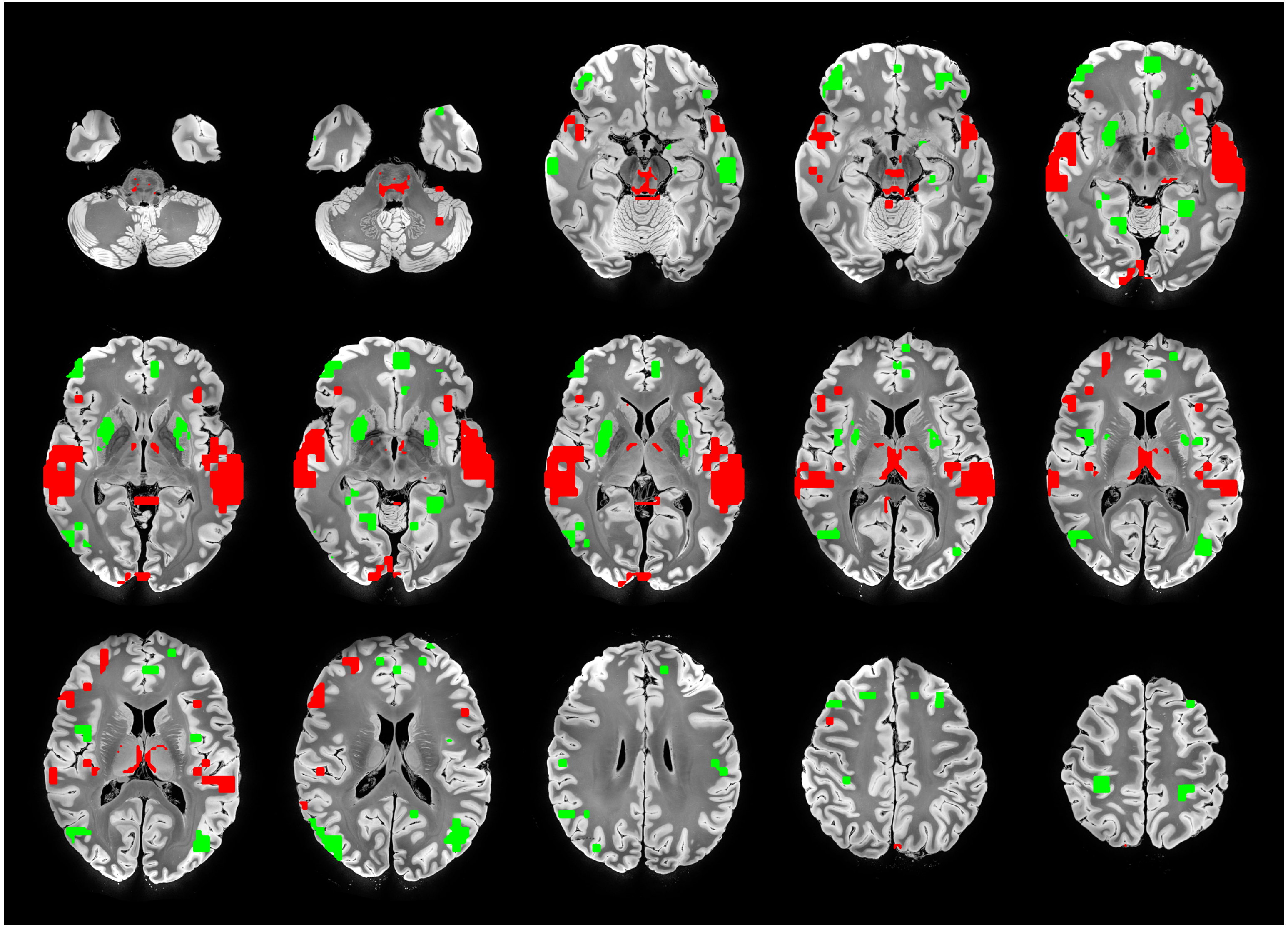

Common activation across all tasks | Common deactivation across all tasks

+4 sec

### Auditory Disjunction

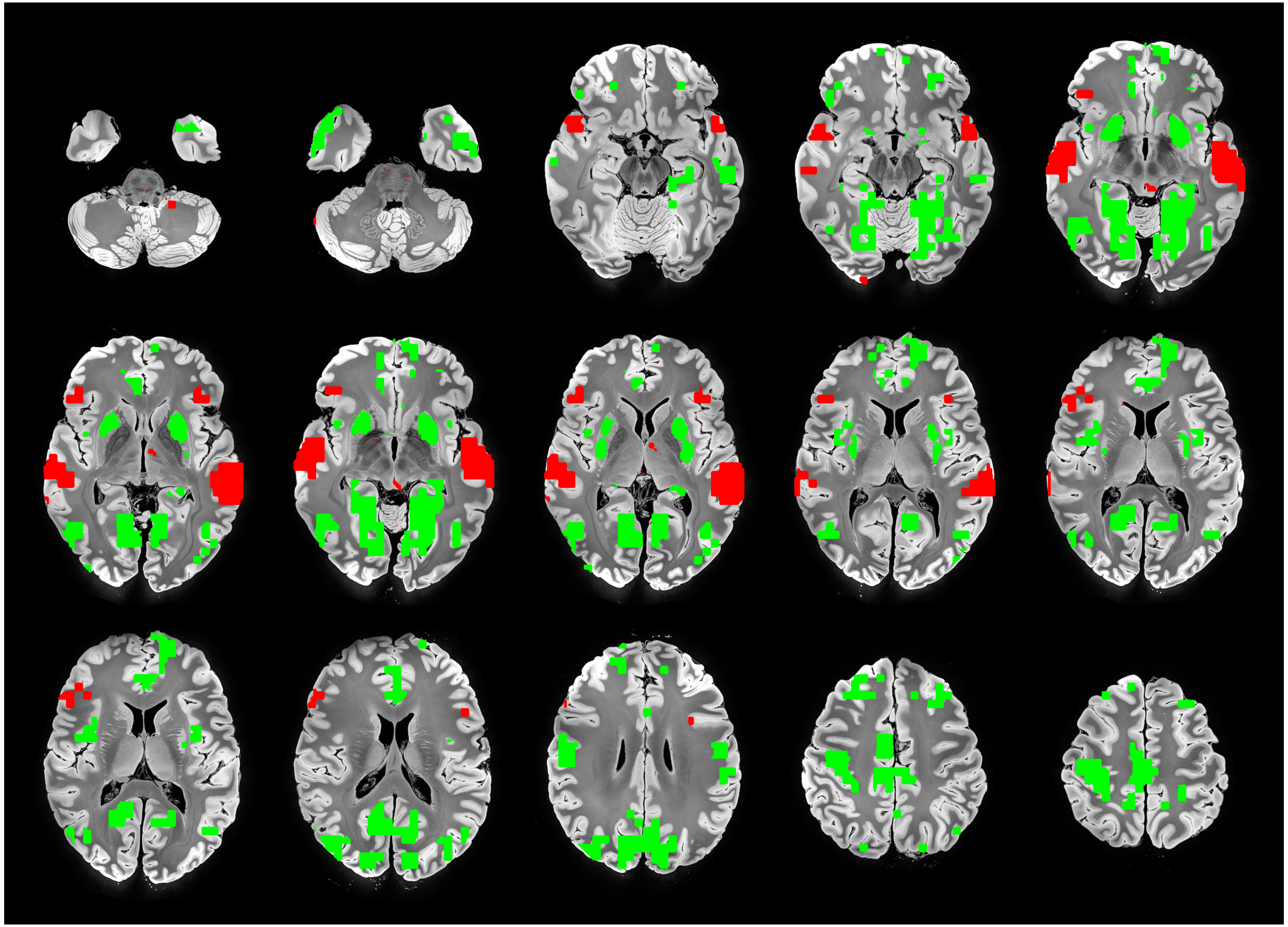

Common activation across all tasks    Common deactivation across all tasks

+6 sec

### Auditory Disjunction

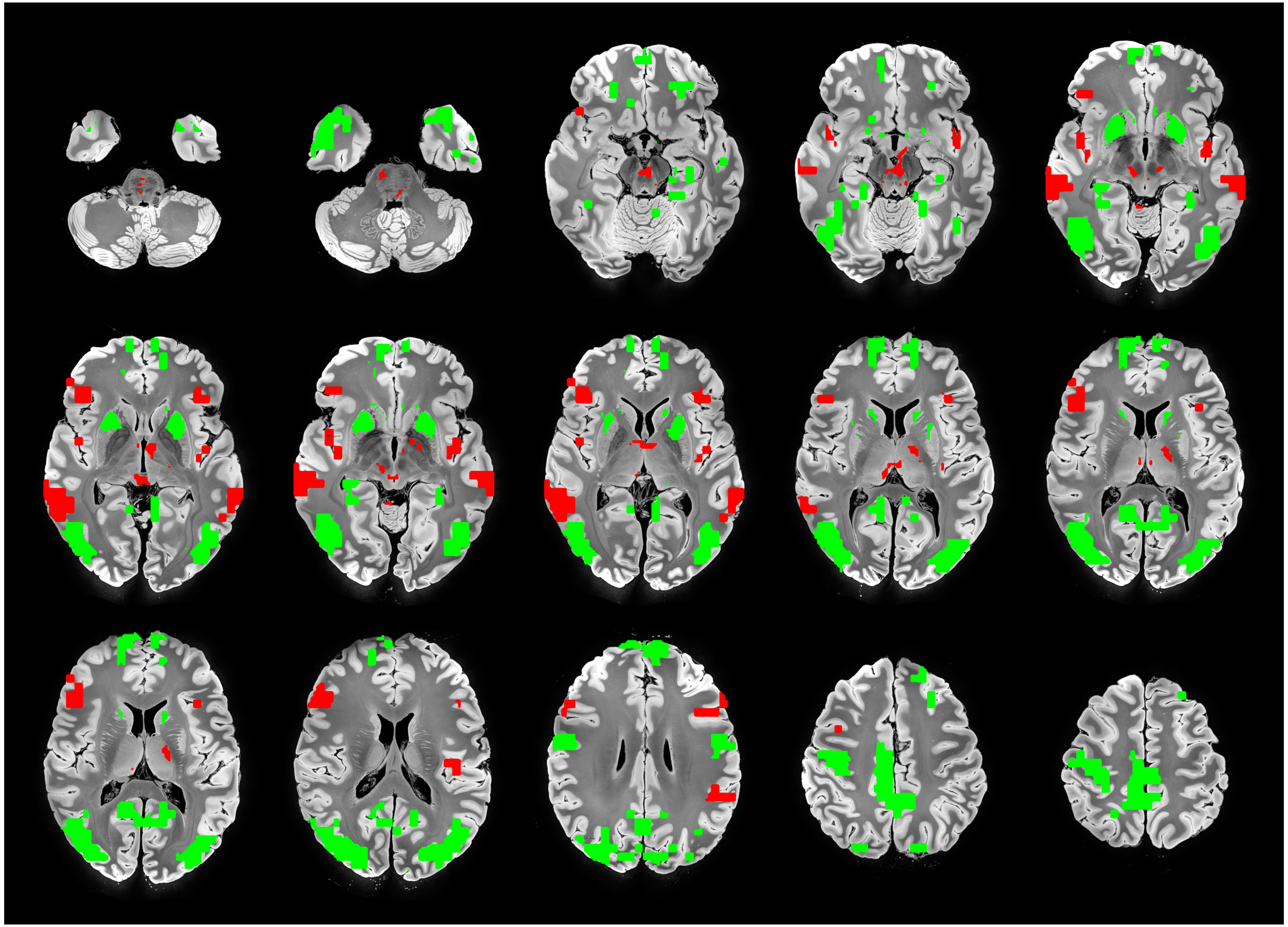

Common activation across all tasks Common deactivation across all tasks

+8 sec

### Taste Disjunction

### Taste Disjunction

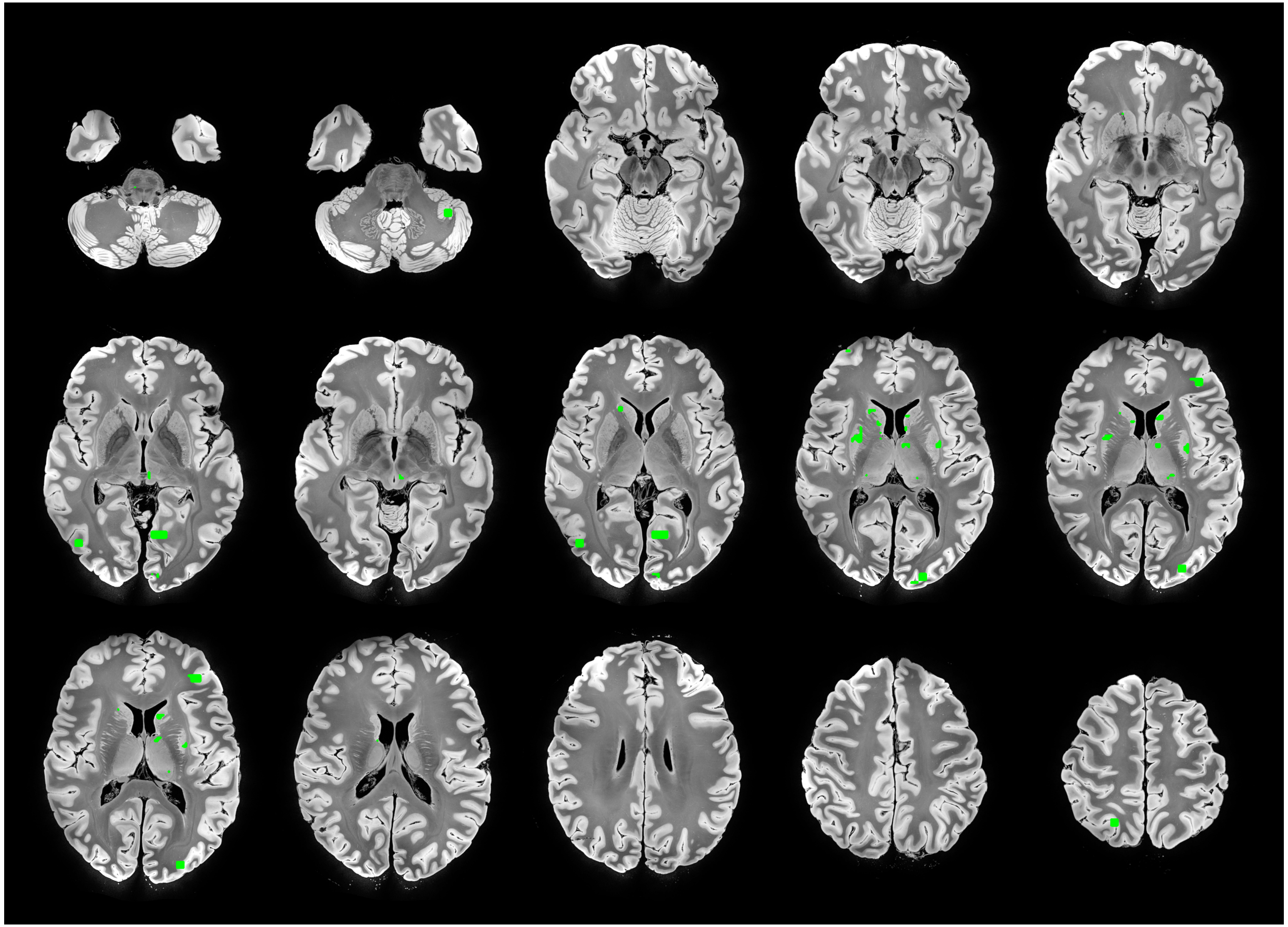

Common activation across all tasks

Common deactivation across all tasks

-4 sec

### Taste Disjunction

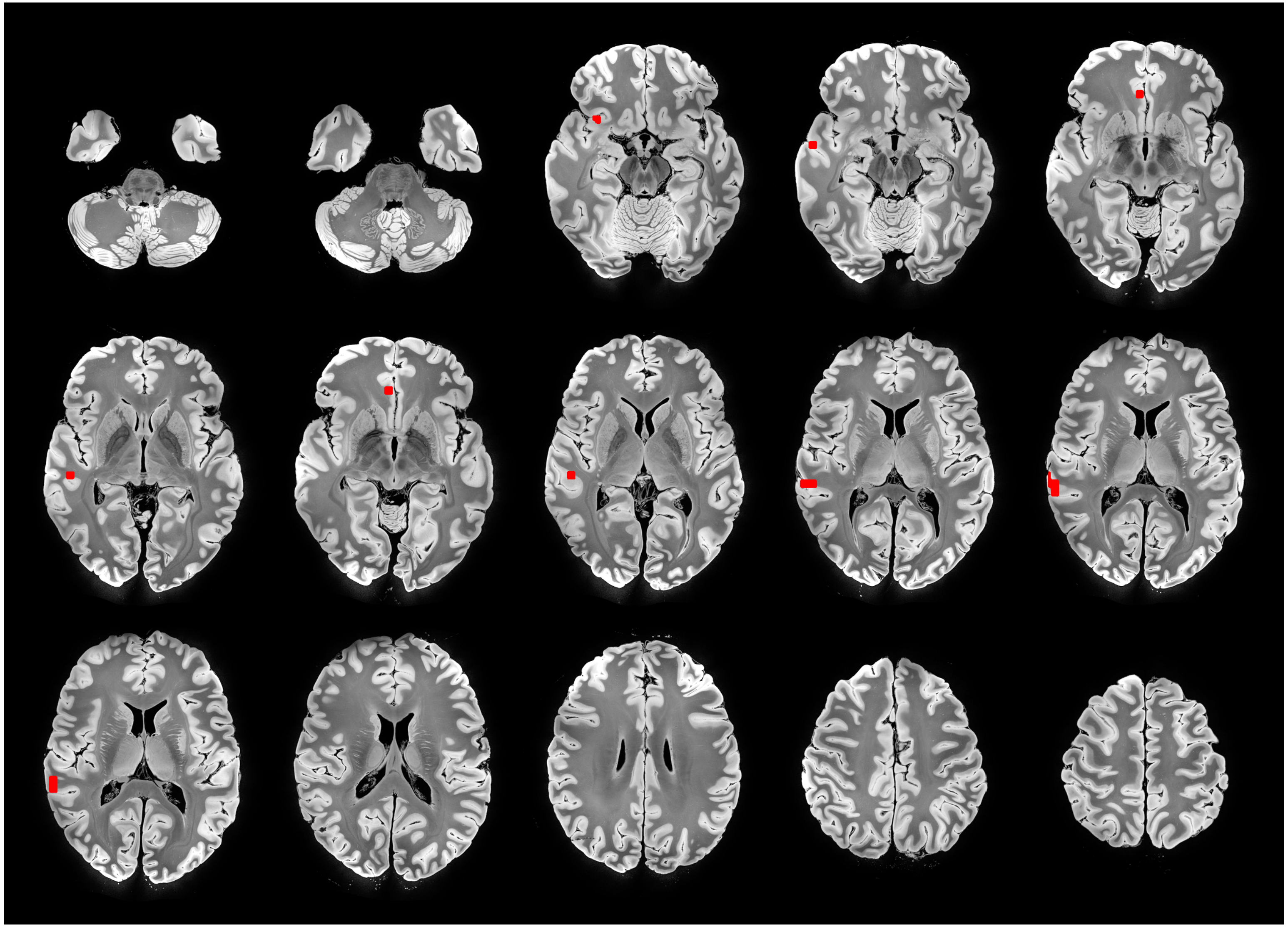

Common activation across all tasks

Common deactivation across all tasks

-2 sec

### Taste Disjunction

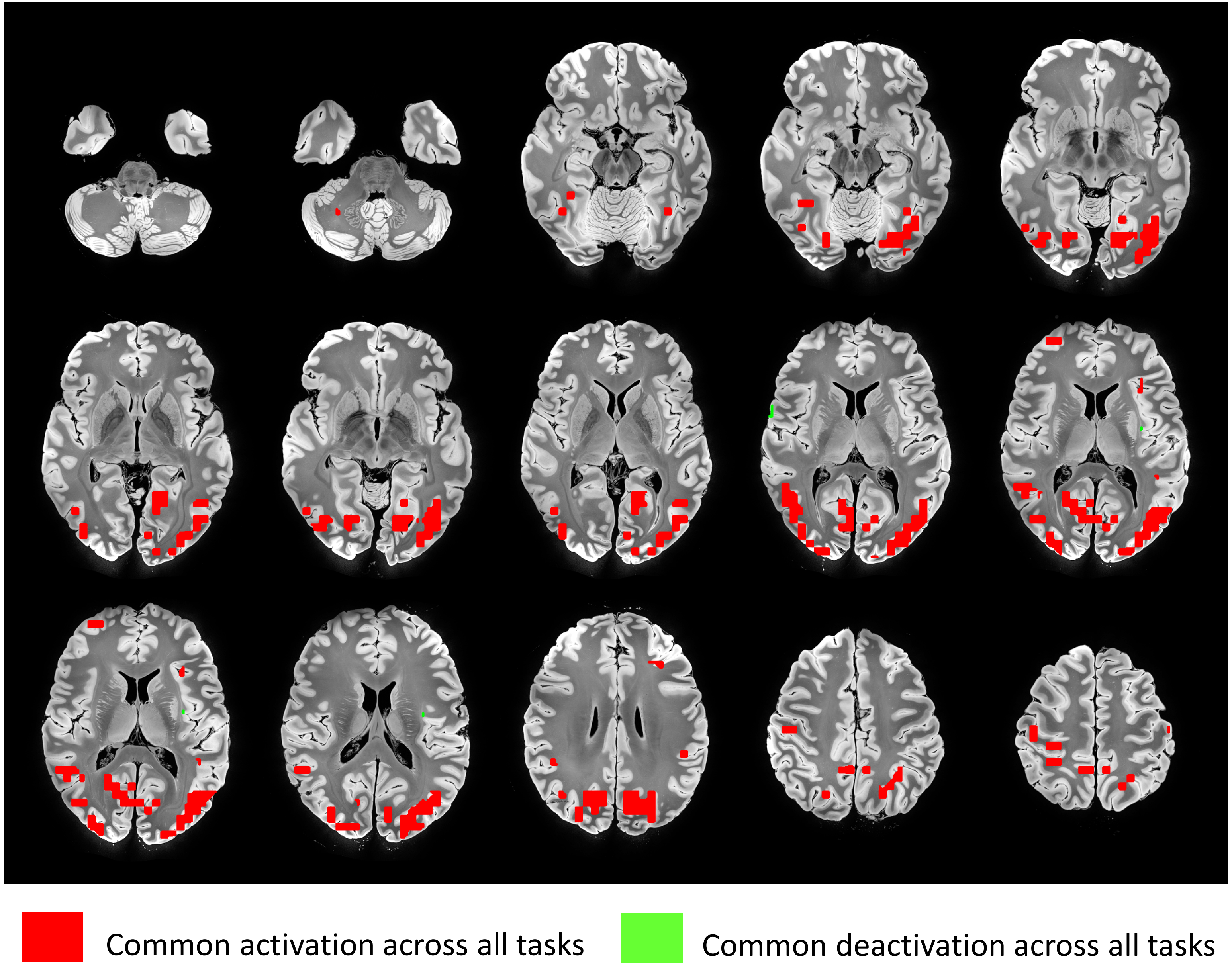



0 sec

### Taste Disjunction

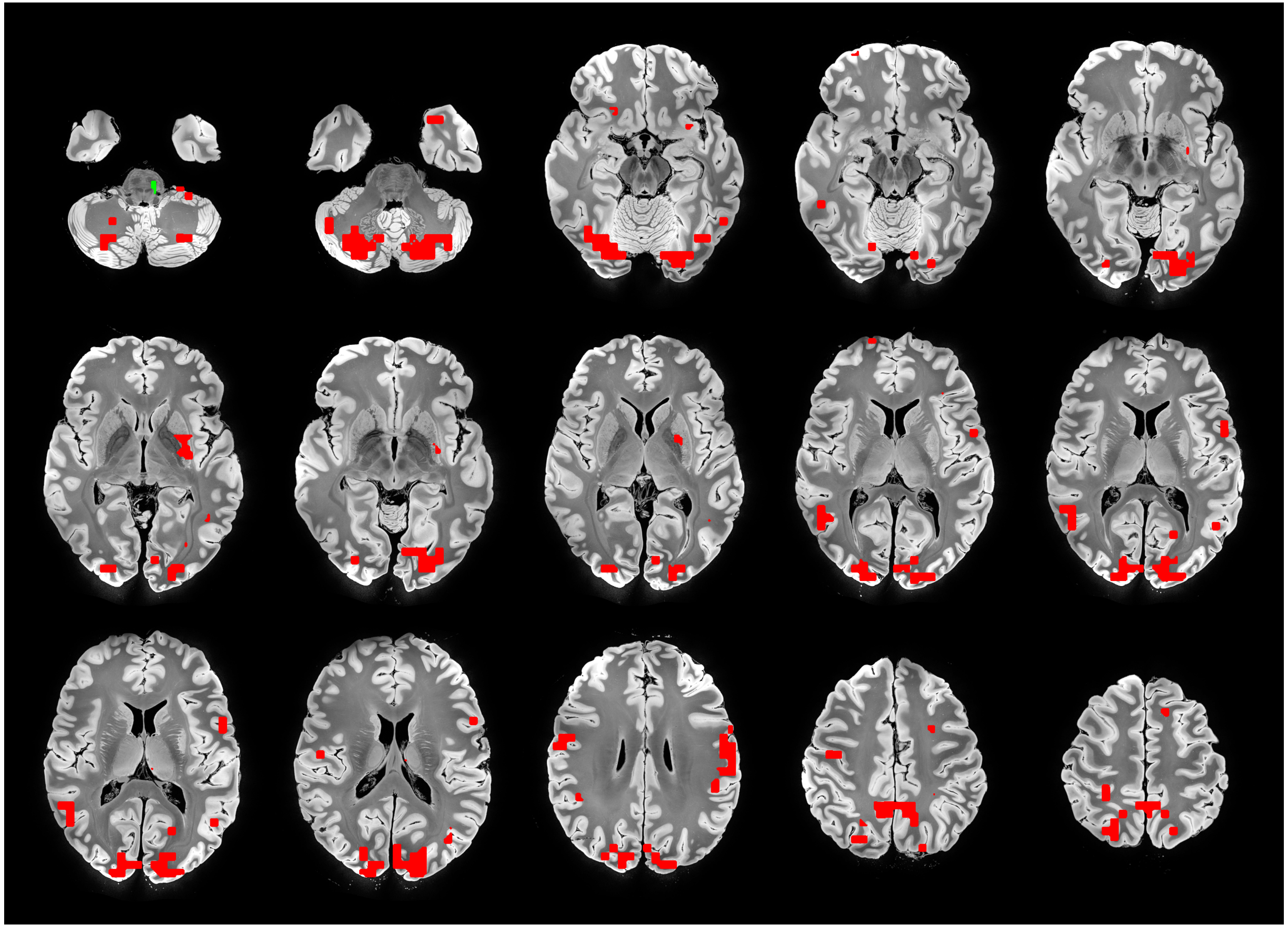

Common activation across all tasks

Common deactivation across all tasks

+2 sec

### Taste Disjunction

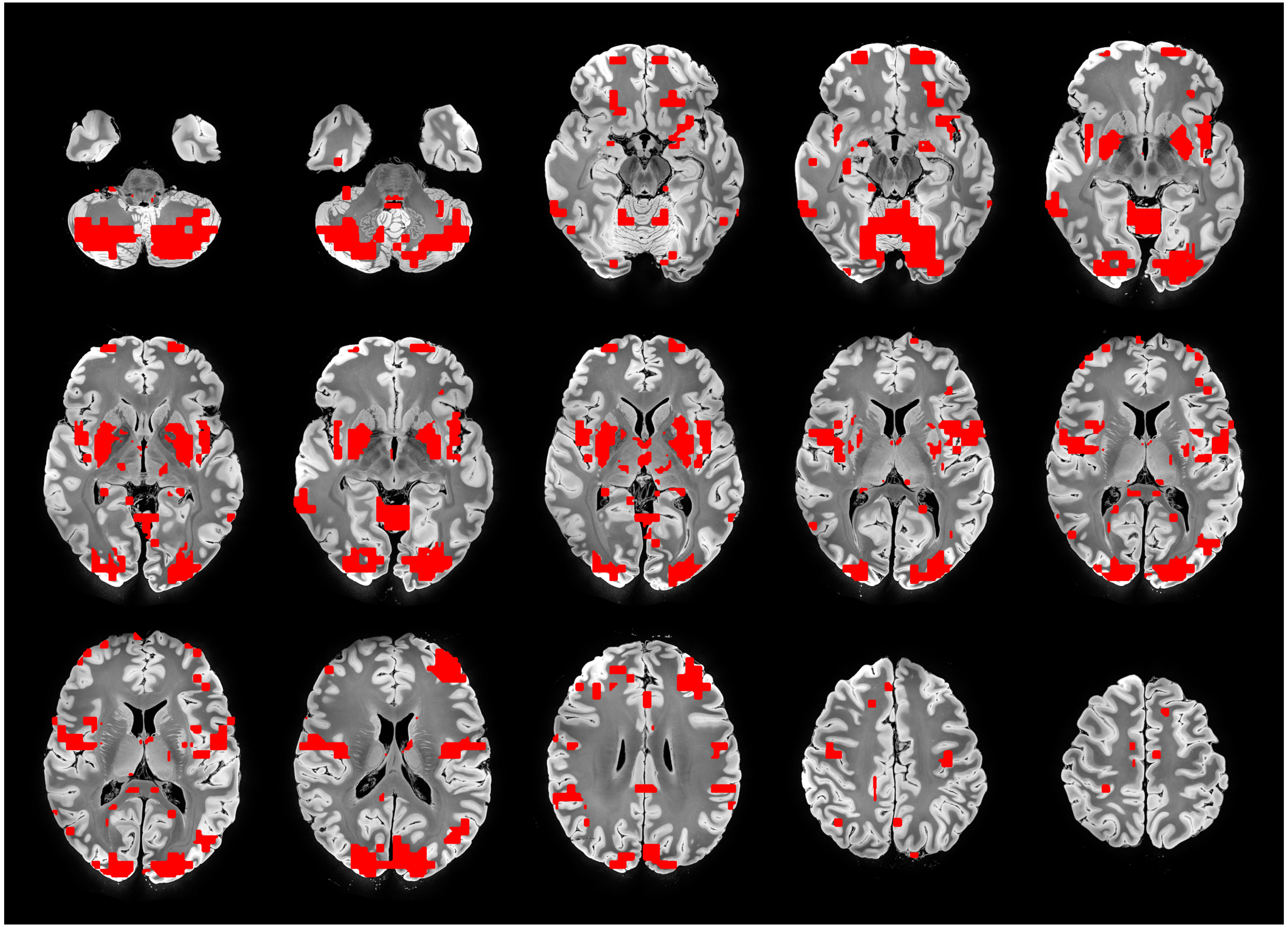

Common activation across all tasks

Common deactivation across all tasks

+4 sec

### Taste Disjunction

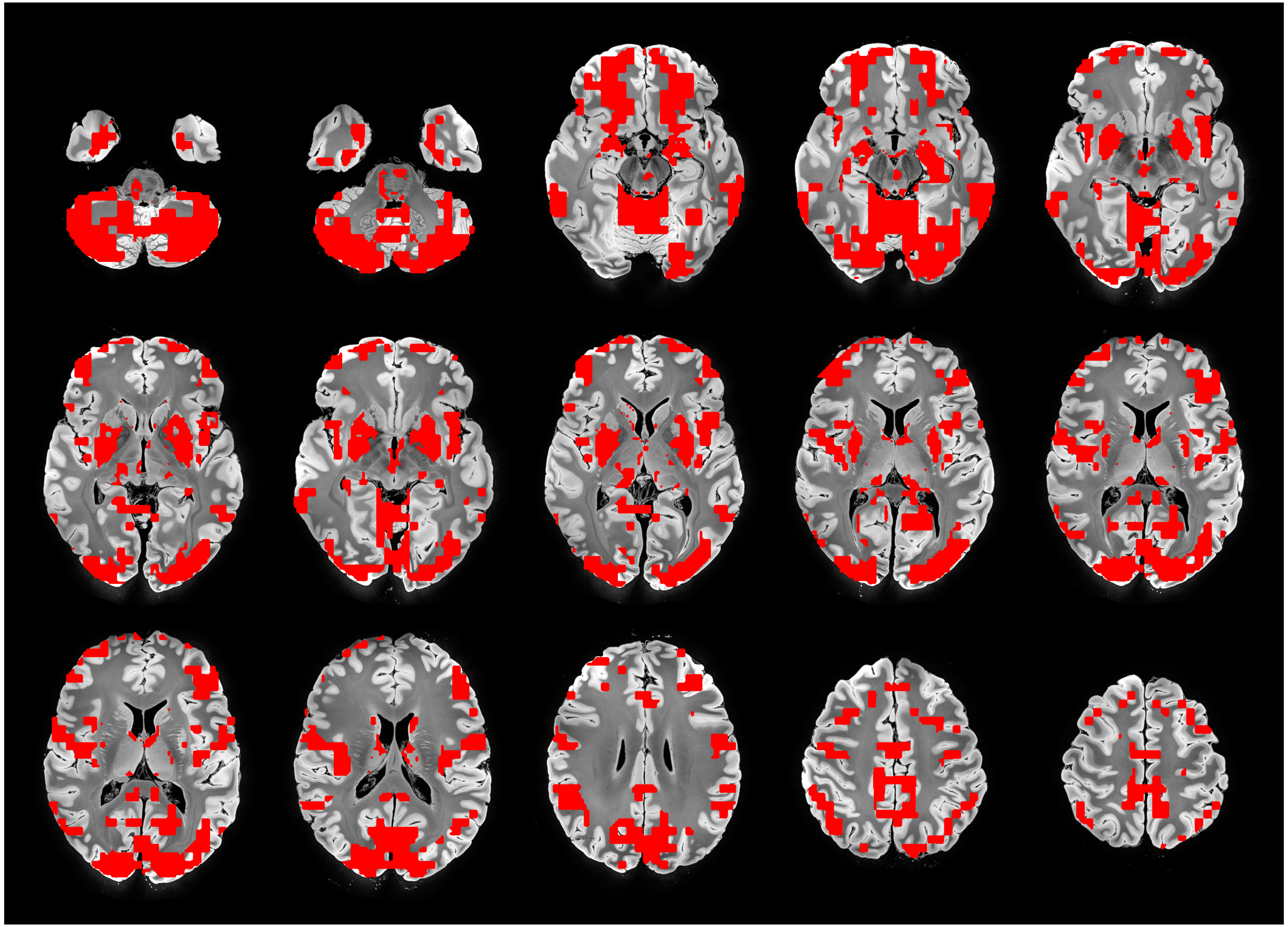

Common activation across all tasks    Common deactivation across all tasks

+6 sec

### Taste Disjunction

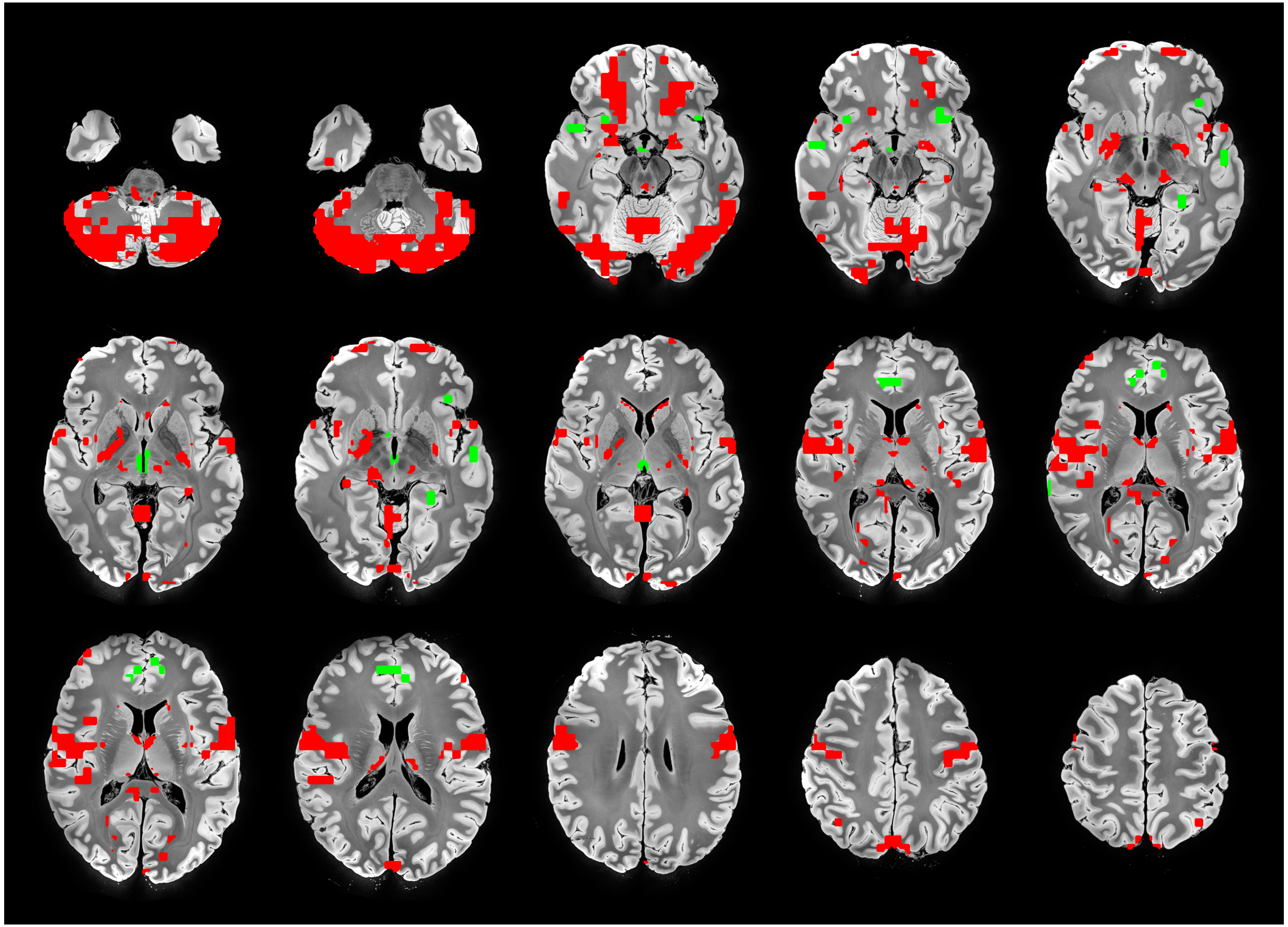

Common activation across all tasks | Common deactivation across all tasks

+8 sec

### Tactile Disjunction

### Tactile Disjunction

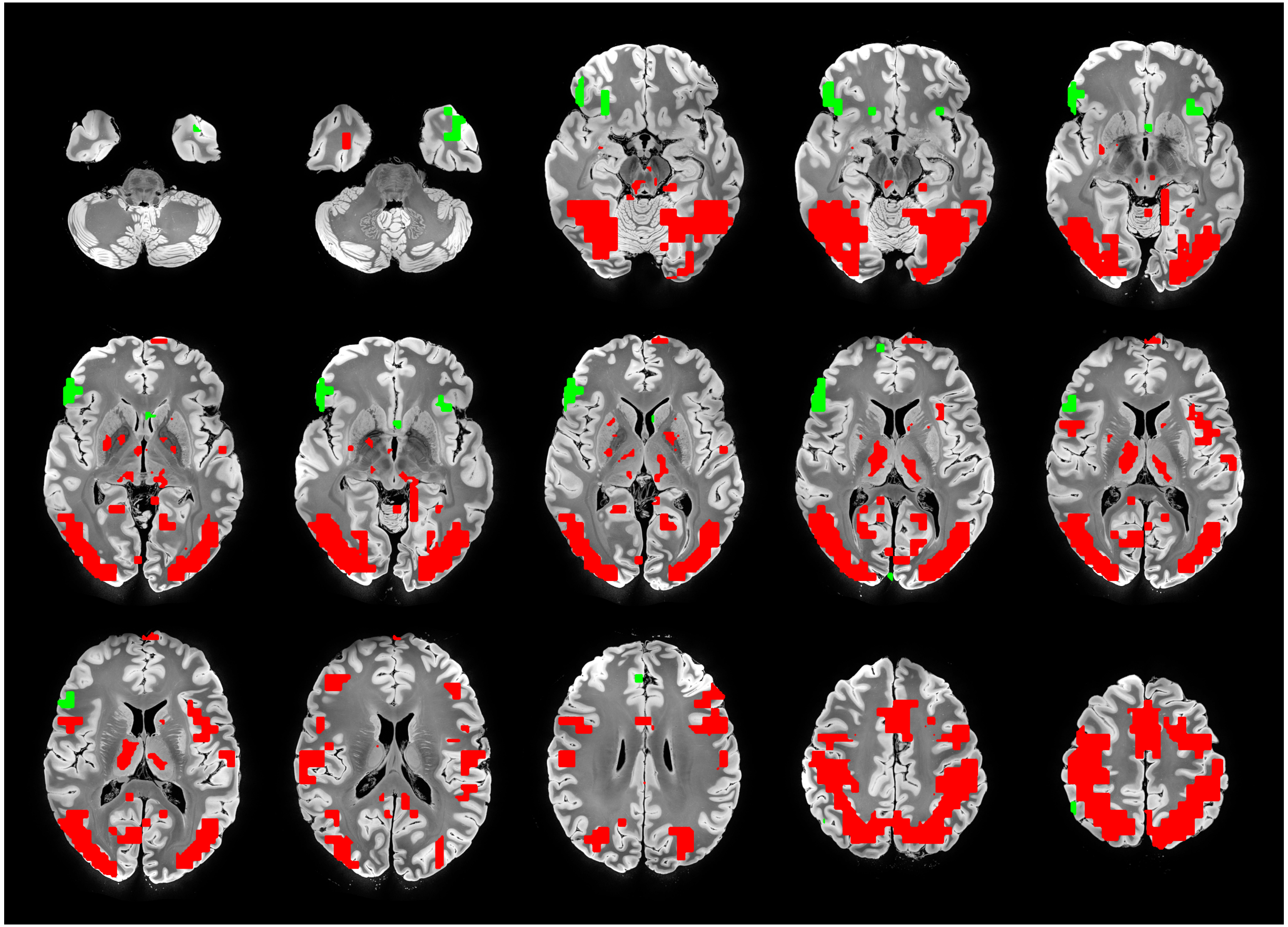

Common activation across all tasks

Common deactivation across all tasks

-4 sec

### Tactile Disjunction

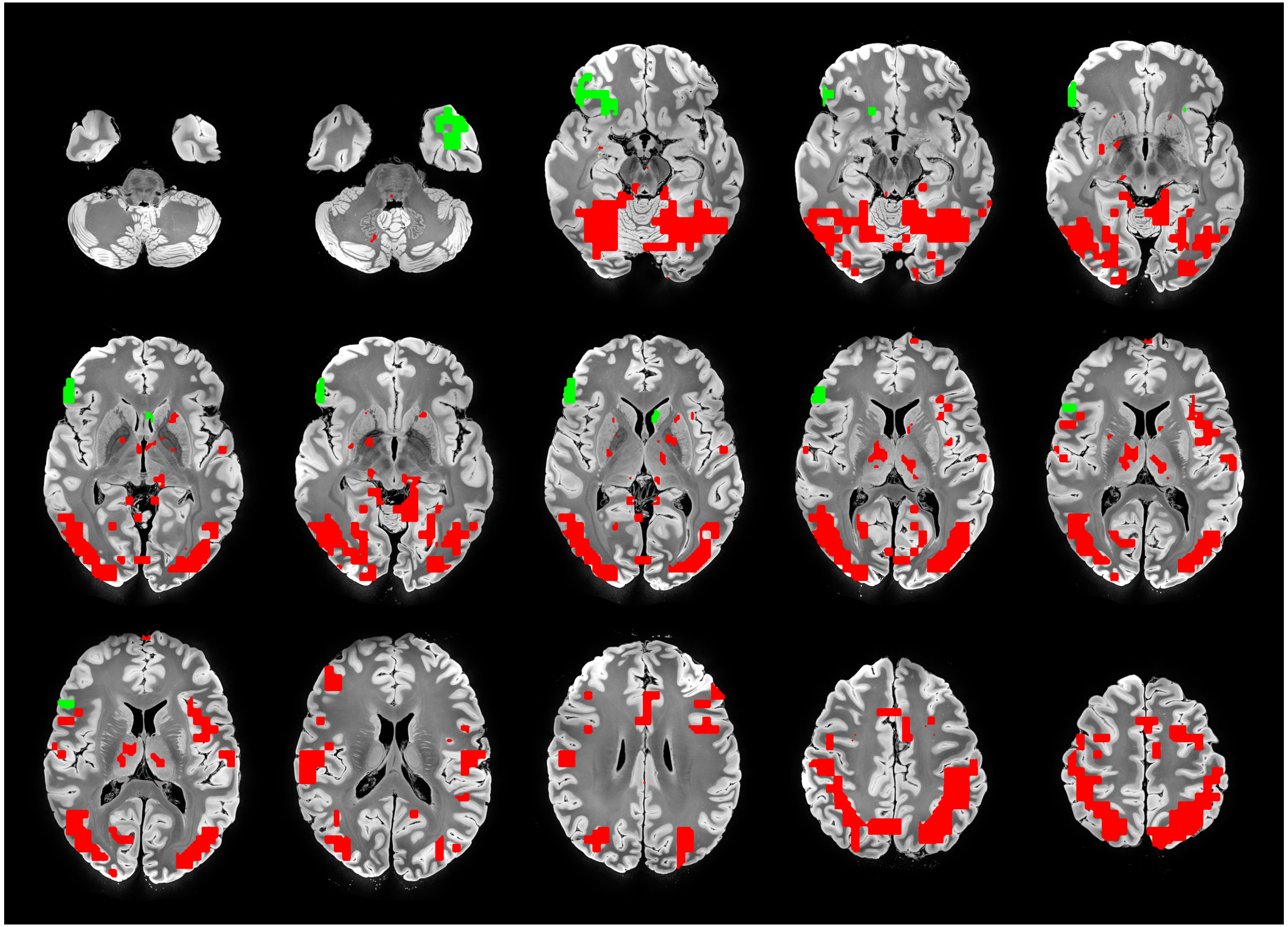

Common activation across all tasks

Common deactivation across all tasks

-2 sec

### Tactile Disjunction

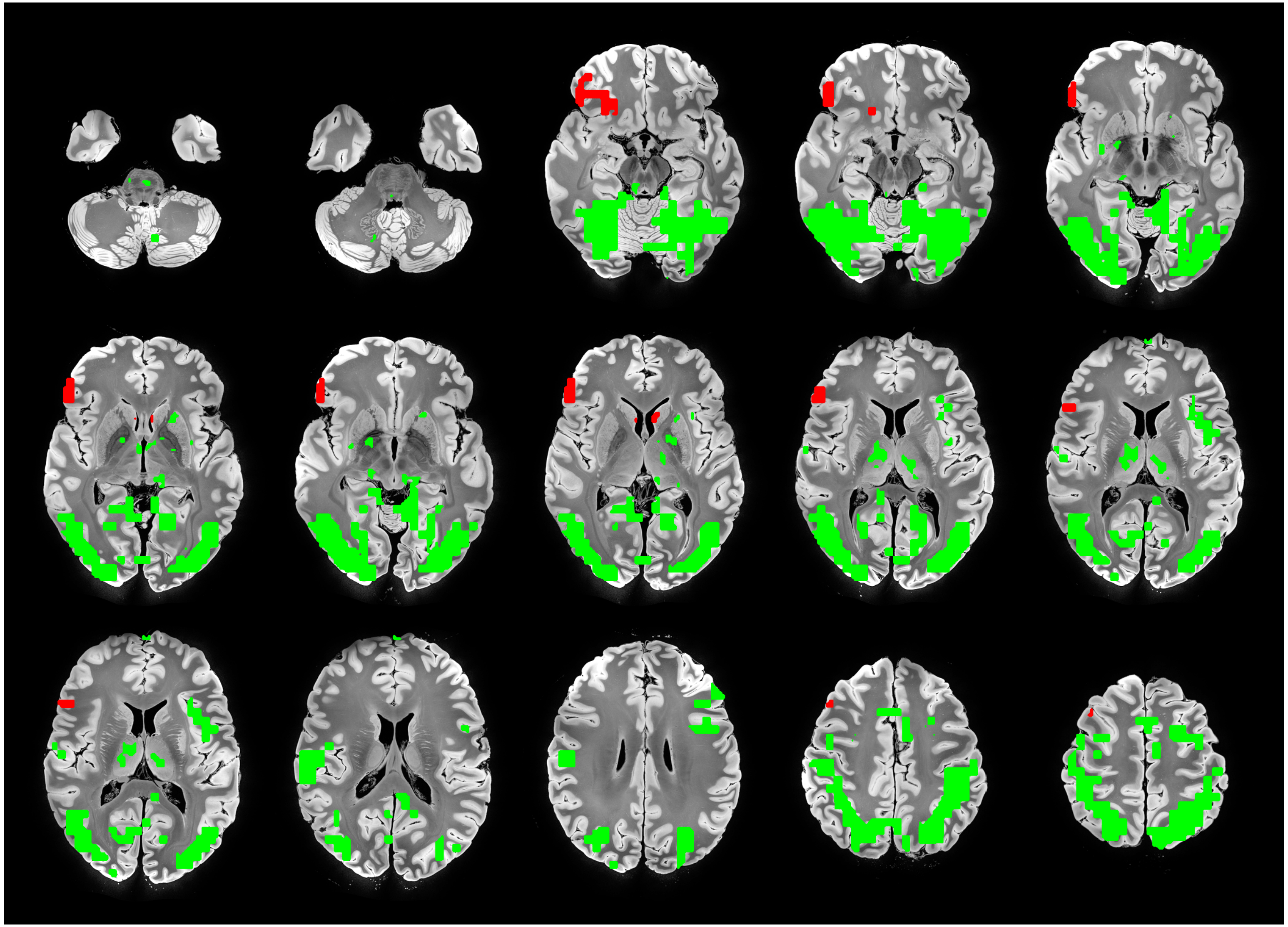

Common activation across all tasks   Common deactivation across all tasks

0 sec

### Tactile Disjunction

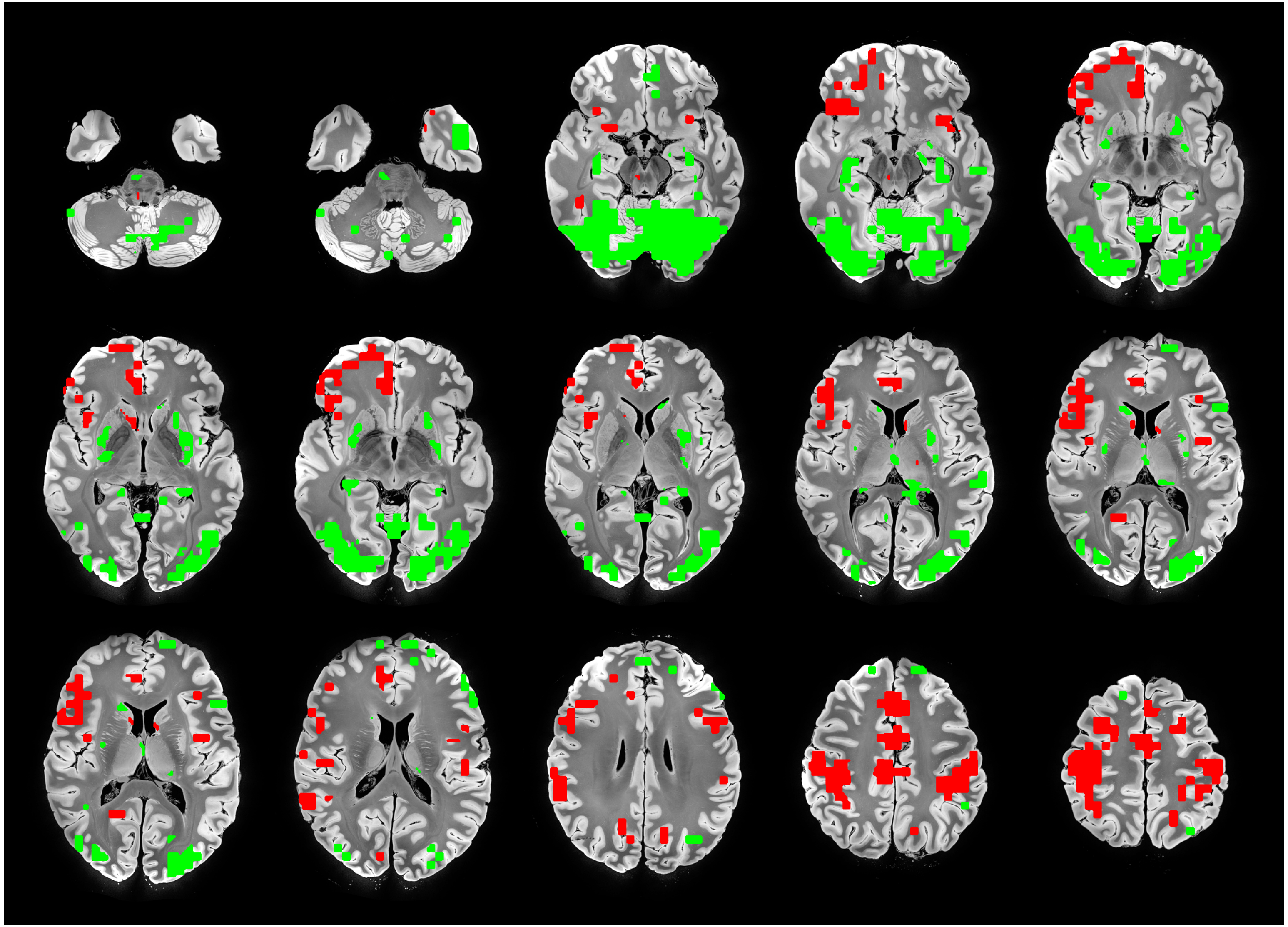

Common activation across all tasks

Common deactivation across all tasks

+2 sec

### Tactile Disjunction

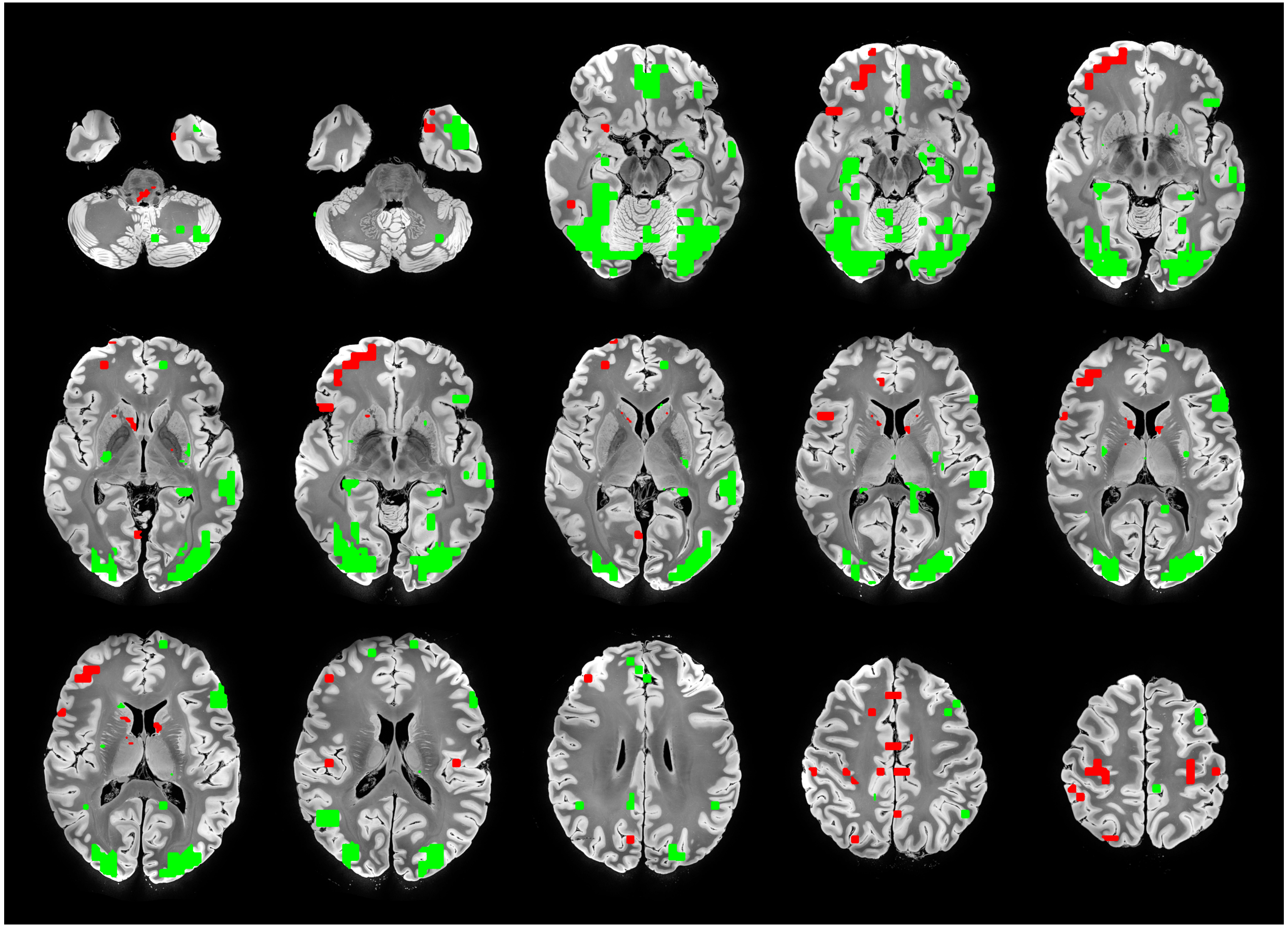

Common activation across all tasks

Common deactivation across all tasks

+4 sec

### Tactile Disjunction

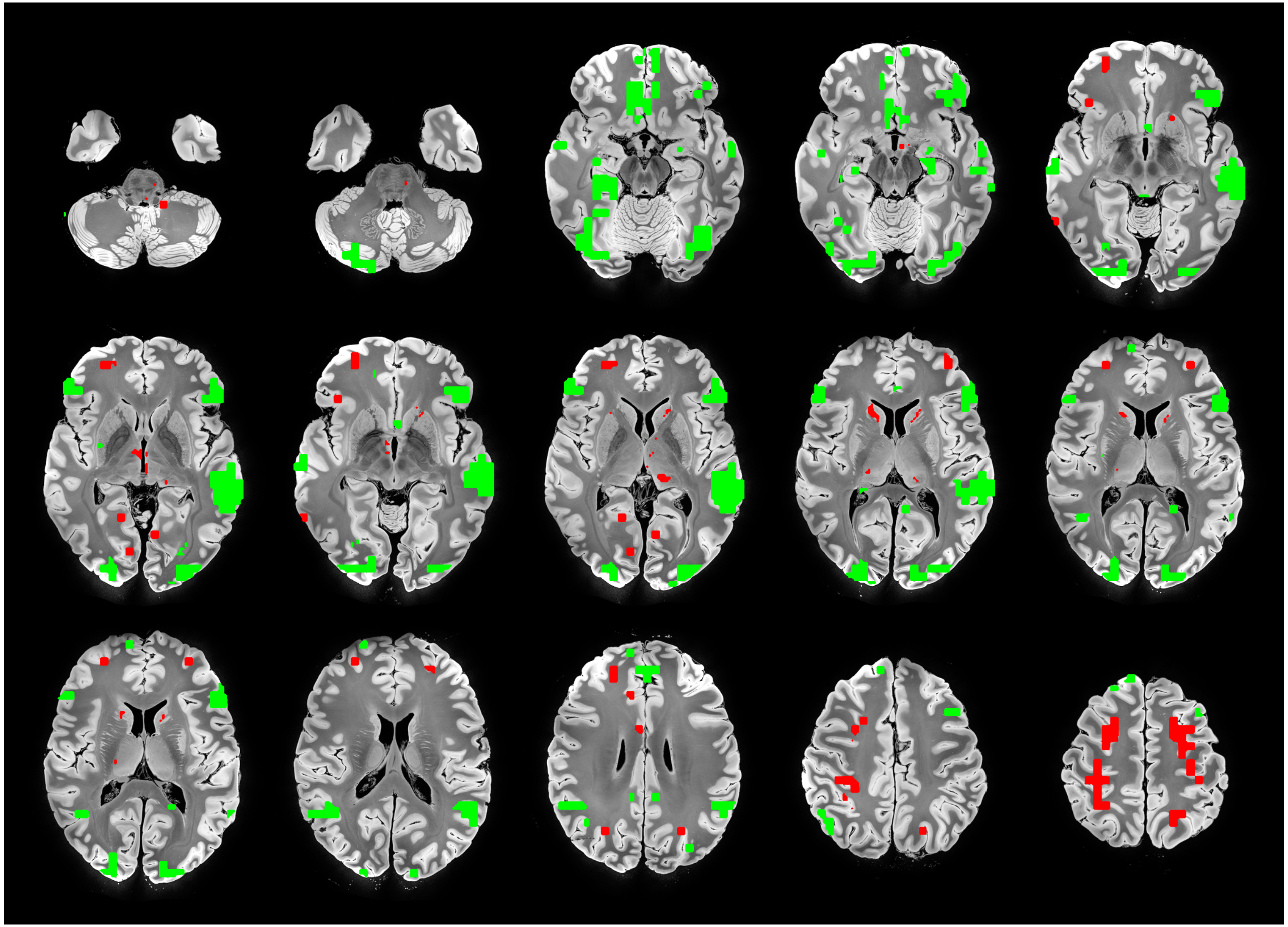

Common activation across all tasks    Common deactivation across all tasks

+6 sec

### Tactile Disjunction

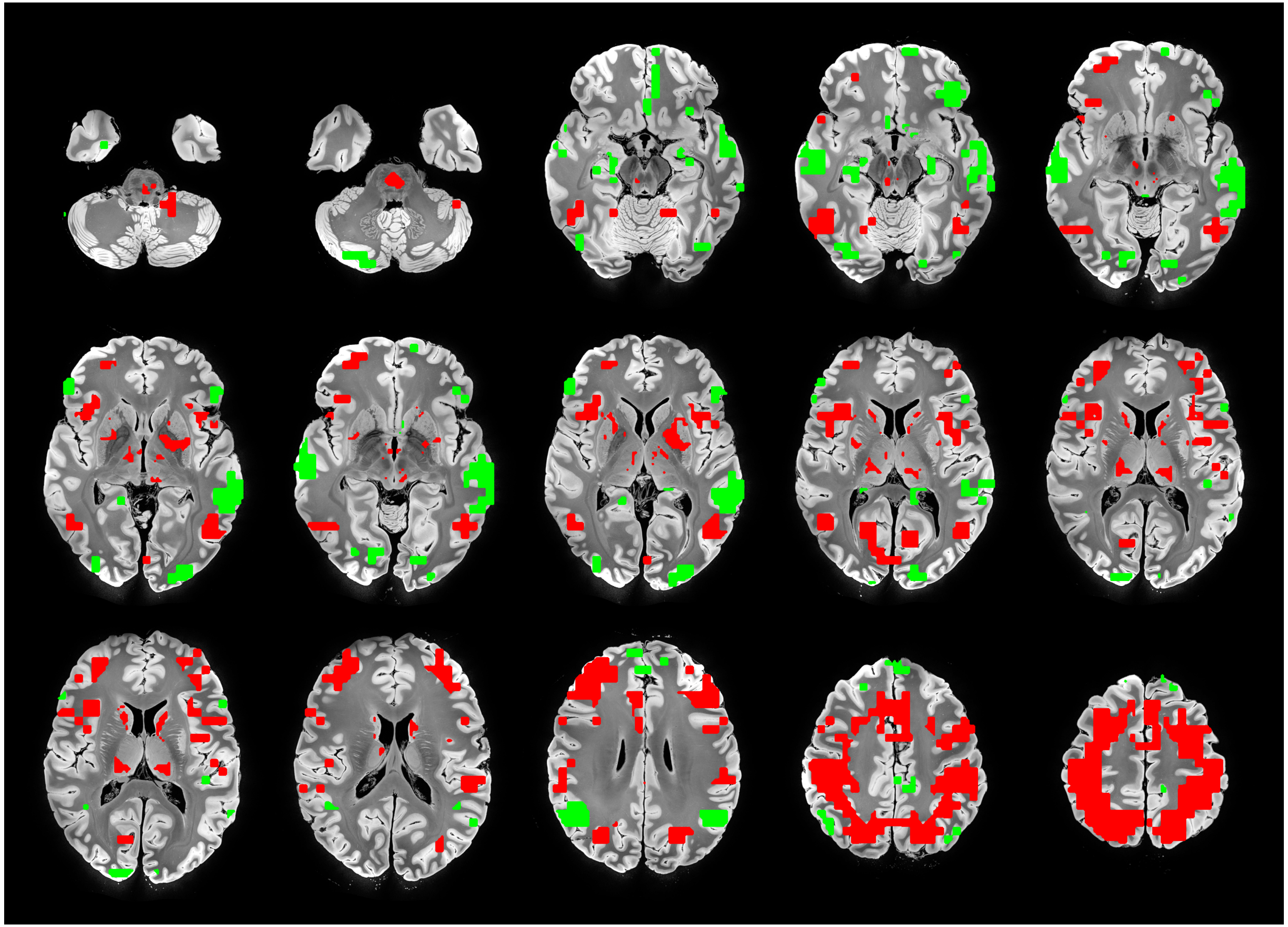

Common activation across all tasks     Common deactivation across all tasks

+8 sec
