## Supplementary Data Guide for "Shared subcortical arousal systems across sensory modalities during transient modulation of attention"

### Supplementary Presentation S1 – Binary and Graded Conjunction Across Modalities

Binary Conjunction Across All Modalities

Shared whole-brain activations (increases) and deactivations (decreases) across four sensory modalities, including, vision, audition, taste, and touch. The shared changes, spanning from -4 to 8 seconds relative to stimulus onset, are visualized on axial MRI brain slices. Cluster-based permutation testing ($p < 0.05$) was employed to identify the statistically significant changes in percentage change BOLD brain maps with respect to the baseline before block/event onset for 11 tasks in four sensory modalities. Whole-brain binary conjunction analysis was applied to the statistical brain maps across all tasks to identify regions and time points sharing statistically significant activations or deactivations across tasks and sensory modalities. The whole-brain binary conjunction analysis showed shared activations in bilateral subcortical arousal networks (midbrain reticular formation and central thalamus), detection, arousal and salience cortical networks (primary visual cortex, anterior insula and anterior cingulate/supplementary motor area), as well as in attention and executive control cortical networks (right anterior inferior parietal lobule, right superior parietal lobule, bilateral medial parietal cortex and bilateral middle frontal gyrus). Shared deactivations were observed in the binary conjunction analysis only at 8 s after the stimulus in the ventral medial frontal cortex, which is a default mode network area.

Graded Conjunction Across All Modalities

Graded conjunction analysis revealed activations (increases) and deactivations (decreases) shared less consistently across sensory modalities. These activations and deactivations were observed across time (-4 to 8 seconds from block/event onset). As in the binary conjunction analysis, the first step was spatiotemporal cluster-based permutation testing ($p < 0.05$) to identify the statistically significant changes in percentage change BOLD brain maps with respect to the baseline before block/event onset for each of the 11 tasks in the different modalities. Binary conjunction was then applied within each modality to identify shared increases or decreases per modality. Finally, graded conjunction analysis was applied across the four sensory modalities to identify regions with shared activations or deactivations in 1, 2, 3 or 4 modalities. The graded conjunction analysis revealed cortical and subcortical regions showing increases or decreases less consistently involved across modalities, in addition to those identified through the binary conjunction analysis.

**Supplementary Presentation S2 – Binary Conjunction Within Each Modality**

Shared whole-brain activations (increases) and deactivations (decreases) analyzed separately for each of the four sensory modalities across time (-4 to 8 seconds from block/event onset). Statistically significant changes in percentage change BOLD brain maps with respect to the baseline before block/event onset were obtained using spatiotemporal cluster-based permutation testing ($p < 0.05$) for each of the 11 tasks in the four modalities. Binary conjunction analysis was then applied across tasks within each sensory modality (vision, audition, taste) to identify regions sharing significant activations or deactivations within that modality. Note that for touch, conjunction analysis was not needed because there was only one tactile task, so we simply show significant activations and deactivations for that task.

**Supplementary Presentation S3 – Exclusive Disjunction for Each Modality**

Unique subcortical and cortical activations (increases) and deactivations (decreases) were observed for each of the four sensory modalities (vision, audition, taste, and touch) from -4 to 8 seconds from block/event onset. Again, the first step was spatiotemporal cluster-based permutation testing ($p < 0.05$) for each of the 11 tasks to identify statistically significant changes in percentage change BOLD brain maps with respect to the baseline before block/event onset, followed by binary conjunction analysis to obtain shared statistically significant increases or decreases within each modality. Disjunction analysis was then applied to identify unique subcortical and cortical activations and deactivations for each sensory modality, not present in the other modalities.
