## Supplementary Information for "Shared subcortical arousal systems across sensory modalities during transient modulation of attention"

### Functional MRI tasks Description

We analyzed 3T task fMRI data collected from healthy adults while performing 11 different tasks spanning four sensory modalities, including, vision, audition, taste, and touch. These data were obtained from six publicly available datasets, including the Human Connectome Project (HCP)[1, 2], UCLA Consortium for Neuropsychiatric Phenomics[3], Glasgow University[4], Jagiellonian University[5], and two datasets from Yale University[6, 7]. For an overview of the task design characteristics and analysis-relevant details see Table 1 in the main manuscript. Detailed description of the tasks and fMRI data acquisition parameters is reported below.

*Human Connectome Project (HCP) Tasks*[1, 2]

HCP structural and functional MRI data were collected on 3T Siemens Connectome Skyra MRI scanner with a 32-channel head coil. Functional MRI data were acquired using a multiband gradient-echo EPI pulse sequence (TR/TE = 720/33.1 ms, 52° flip angle, 72 axial slices, 2 mm isotropic voxels) in addition to a structural high resolution T1 scan (TR/TE = 2400/2.14 ms, 8° flip angle, 256 axial slices, 0.7 mm isotropic voxels). Two functional MRI runs were acquired per participant. See Reference Manual of HCP 1200 Subjects Data Release for detailed information about the acquisition parameters and the tasks[8].

<u>HCP Gambling Task</u>

The gambling task was adapted from a previous version developed by Delgado and Fiez[9]. In this task, participants played a card guessing game in which they were asked to guess if the number on a mystery card (represented by a "?") is more or less than 5 to win or lose money. Participants responded with their right hand through pressing one of two buttons on the response box. Each trial started with the presentation of the mystery card for up to 1500 ms. If participants responded before this interval elapsed, a fixation cross was displayed for the remaining time followed by a 1000 ms feedback period during which the value of the mystery card was revealed. The feedback included either 1) a green upward arrow with "$1" indicating a reward trial, 2) a red downward arrow adjacent to "-$0.50" representing a loss trial, or 3) the number 5 alongside a gray double-headed horizontal arrow for a neutral trial. A 1000-millisecond inter-trial interval (ITI) with a fixation cross was then presented, yielding a total trial duration of 3.5 seconds. Each task block contained 8 trials and lasted for 28 seconds. Two runs were acquired per subject. In each run, there was an initial 8s countdown sequence of numbers displayed before the first task block, followed by 2 mostly reward and 2 mostly loss blocks, interleaved with 4 fixation blocks (15 seconds each). Total fMRI run duration was 192 seconds with 258 volumes acquired per run.

Within each run, we analyzed fMRI data marking the transition from baseline (fixation) blocks to task blocks.

HCP Relational Processing Task

This task was adapted from a previous version developed by Christoff and colleagues[10]. The task included two conditions: relational processing and control matching. Stimuli presented in the two conditions included 6 different shapes filled with one of six different textures. In the relational processing condition, participants were presented with 2 pairs of objects, one pair positioned at the top and the other pair at the bottom of the screen. They were instructed to assess what dimension differs across the top pair (either shape or texture) and then determine whether the bottom pair also differs along that same dimension. In the control matching condition, participants observed two objects at the top of the screen and one object at the bottom, as well as a word in the middle of the screen ("shape" or "texture"). Their task was to decide whether the bottom object matches either of the top two objects along the dimension specified by the written word. In both conditions, participants responded yes or no with their right hand using one of two buttons. For the relational condition, each stimulus was presented for 3500 ms, with a 500 ms ITI. In the matching condition, stimuli were presented for 2800 ms, with a 400 ms ITI. Relational processing blocks contained 4 trials per block while matching blocks contained 5 trials per block. Each block (relational or matching) lasted for a total of 18 seconds. In each run, there were three relational blocks, three matching blocks and three 16-second fixation blocks. Total fMRI run duration was 176 seconds with 232 volumes acquired per run. Within each run, we analyzed fMRI data marking the transition from baseline (fixation) blocks to task blocks.

HCP Working Memory Task

The working memory task was developed based on previous experimental designs[11, 12]. Participants were presented with blocks of trials featuring pictures of places, tools, faces and body parts. Within each run, the four different stimulus types were presented in separate blocks. Half of the blocks within each run involved a 2-back working memory task and the other half involved a 0-back working memory task and participants responded with their right hand. A 2.5 second cue at the onset of each block indicated the task type (and target for 0-back). Each run included eight task blocks, each lasting 25 seconds (with 10 trials of 2.5 seconds each), interleaved with four fixation blocks, each lasting 15 seconds. During each trial, the stimulus was presented for 2 seconds, followed by a 500 ms ITI. Total fMRI run duration was 301 seconds with 405 volumes acquired per run. Within each run, we analyzed fMRI data marking the transition from baseline (fixation) blocks to task blocks.

HCP Motor Task

This task was adapted from the one developed previously by Buckner and colleagues[13]. Participants were instructed to respond to visual cues by either tapping their left or right fingers, squeezing their left or right toes, or moving their tongue to map motor areas. Each block corresponding to a specific movement type lasted 12 seconds, encompassing 10 movements, and was preceded by a 3-second cue to indicate the specific movement type that participants should do. In each of the two runs, participants completed a total of 10 task blocks interleaved with three 15-second fixation blocks. The task blocks included 2 blocks involving tongue movements, 4 blocks involving hand movements (2 for the right hand and 2 for the left hand), and 4 blocks involving foot movements (2 for the right foot and 2 for the left foot). Total fMRI run duration was 214 seconds with 284 volumes acquired per run. Within each run, we analyzed fMRI data marking the transition from baseline (fixation) blocks to task blocks.

HCP Social Cognition Task

Participants viewed video clips of 20 seconds duration showing objects (squares, circles, triangles) either interacting in some way or moving randomly across the screen. These videos were developed by either Castelli and colleagues[14] or Martin and colleagues[15]. Subsequent to each video clip, participants were asked to report with their right hand whether the objects portrayed a mental interaction (defined as an interaction as if the shaped are considering each other's feelings and thoughts), not sure, or no interaction indicating seemingly random movement. Each of the two runs consisted of five video blocks lasting 20 seconds each interleaved with five fixation blocks lasting 15 seconds each. One run comprised two mental interaction and 3 random interaction video blocks, while the other run included three mental interaction and two random interaction video blocks. Total fMRI run duration was 207 seconds with 274 volumes acquired per run. Within each run, we analyzed fMRI data marking the transition from baseline (fixation) blocks to task blocks.

HCP Language Processing Task

The language task was developed by Binder and colleagues[16]. Each run in the task contained four blocks of a story task interleaved with four blocks of a math task. The durations of the blocks varied, averaging approximately 30 seconds. However, the design ensured that the length of the math blocks matches that of the story blocks. During the story blocks, participants listened to brief auditory stories (consisting of 5-9 sentences) adapted from Aesop's fables. Following each story, participants were presented with a 2-alternative forced-choice question regarding the topic of the story. The math task blocks also presented auditory trials, requiring participants to solve addition and subtraction problems. Each trial included a series of arithmetic operations (e.g., "fourteen plus twelve"), followed by the word "equals" and two answer choices (e.g., "twenty-nine or twenty-six"). Participants indicated their selection by pressing a button with their right hand corresponding to the first or the second answer. The math task incorporated adaptive features to maintain a consistent level of difficulty across participants. Participants were asked to keep their eyes closed during the scan. Total fMRI run duration was 237 seconds with 316 volumes acquired per run. For analysis purposes, we analyzed fMRI events defined as the onset of the auditory question after each story or math problem.

*UCLA Consortium for Neuropsychiatric Phenomics*[3]

MRI data were acquired on 3T Siemens Trio scanners at UCLA. Functional MRI data were acquired using a T2*-weighted EPI sequence (TR/TE = 2000/30 ms, 90° flip angle, 34 axial slices, 4 mm isotropic voxels). Additionally, a T1-weighted MPRAGE high-resolution anatomical scan was collected for each participant (TR/TE = 1900/2.26 ms, 176 axial slices, 1 mm isotropic voxels).

We analyzed fMRI data from the spatial capacity task (SCAP) acquired as part of the UCLA consortium dataset that included six fMRI tasks[3]. Participants were presented with a target array of 1, 3, 5, or 7 yellow circles positioned pseudo-randomly around a central fixation cross. To ensure adequate encoding of the target array and minimize potential biases related to set size interaction, a relatively lengthy stimulus presentation time of two seconds was employed. Following a variable delay of 1.5, 3 or 4.5 seconds, participants were presented with a single green circle, and were then asked during a 3-second fixed response interval to determine whether the green circle was in the same position as one of the target circles from the previous array. Half of the trials were true-positive while the other half were true-negative. Each run included 48 trials and each participant attended one run. Throughout the 48 trials, a central fixation point remained visible. For fMRI analysis purposes we defined event onset as appearance of the green circle.

*Glasgow University*[4]

All Functional and structural scans were acquired on a 3 T Siemens Tim Trio scanner at the Centre for Cognitive Neuroimaging in the University of Glasgow. fMRI data were acquired with a single-shot gradient-echo EPI sequence (TR/TE = 2000/30 ms, 77° flip angle, 32 axial slices, 3 mm isotropic voxels). In addition to the functional data, a high-resolutionT1-weighted sagittal scan was acquired for each subject (1 mm3 isotropic voxels; acquisition matrix 256 × 256 × 192). Participants were scanned while passively listening to the stimuli and keeping their eyes closed.

Participants performed a passive listening task that included 40 blocks of either vocal (20 blocks) or non-vocal (20 blocks) sounds[4] with each block lasting for 8 seconds. These sound blocks were interleaved with 20 periods of silence (baseline) each lasting for 12 seconds. Blocks consisted of a variety of either vocal or non-vocal sounds, with a maximum delay of 400 ms between consecutive stimuli. The order of stimuli was randomly assigned but remained fixed for all subjects. Vocal blocks contained sounds of human vocal origin (excluding sounds without vocal fold vibration such as whistling or whispering). Non-vocal blocks included natural sounds (e.g., falls, sea waves, wind) and animal sounds (e.g., cats, dogs, lions, elephants), alongside sounds from man-made sources (e.g., cars, glass, alarms, clocks, and classical music pieces). Each participant completed one run with total run duration of 620 seconds corresponding to 310 volumes. fMRI data marking the transition from baseline (silence) blocks to task (sound) blocks were analyzed.

*Yale University*[6, 7]

MRI scans were acquired using a Siemens 3.0 Tesla TIM Trio scanner with a 32-channel head coil at Yale University Magnetic Resonance Research Center. A T1-weighted 3D MPRAGE whole brain image was acquired (TR/TE= 1900ms/2.52ms, 9° flip angle, 176 axial slices, 1 mm isotropic voxels). T2*-weighted functional brain images were acquired using a multiband susceptibility-weighted single-shot EPI sequence (TR/TE: 1000ms/30ms; MB=4, 60° flip angle, 60 axial slices, 2 mm isotropic voxels). Participants were instructed to keep their eyes closed during the scan.

<u>Taste Perception I</u>

Five taste stimuli were administered to the participants in the scanner using a custom-designed gustometer that ensures delivery of precise amounts at precisely timed intervals and durations. The taste stimuli included a sweet (sucrose), sour (citric acid), salty (sodium chloride), bitter (quinine sulfate), and a tasteless solution that mimics the ionic composition of saliva. Participants completed four runs lasting 580 seconds each. Within a run, there were eight taste blocks (one block for each of the four taste qualities and four tasteless blocks), and eight rest blocks. During taste each block, uncued delivery of one stimulus was repeated for four, six, or eight times. The stimulus was delivered over a period of 3 seconds followed by a 6-second interval for the participant to swallow. After each taste block, there was a 1 ml rinse with water followed by a 15-second rest period. fMRI data marking the transition from baseline (rest) periods to task (taste) blocks were analyzed.

Taste Perception II

This task was designed by the same group that developed the taste task detailed earlier (Taste Perception I) and shares a similar task structure. The delivered taste stimuli included a sweet sucrose solution, a sour citric acid solution, a salty sodium chloride solution, an umami monopotassium glutamate solution, as well as a tasteless and odorless solution. Each run consisted of 12 taste blocks (two blocks for each taste quality and four tasteless blocks) and 12 rest blocks. A single stimulus was presented in each taste block for four, six, or eight times. Each time, the stimulus was delivered over two seconds followed by a seven-second swallowing period. Following each taste block, there was a rinsing period with water over 2 seconds then a 10-second rest-period before the onset of the next taste block. Each participant completed two runs, lasting 722 seconds each. fMRI data marking the transition from baseline (rest) periods to task (taste) blocks were analyzed.

*Jagiellonian University*[5]

Structural and functional MRI data were acquired using a 3 T S MAGNETOM Trio scanner with a 32-channel head coil. Functional MRI data were collected using an EPI sequence (TR/TE= 3000/30 ms, 90° flip angle, 48 axial slices, 2.1 x 2.1 mm in plane voxel size, slice thickness 2 mm). A 3D T1-weighted MPRAGE scan was acquired for each subject(TR/TE=2530/3.32 ms, 176 axial slices, 1 mm isotropic voxels).

During the fMRI scan, participants engaged in a task in which they were presented with numerosities 2, 4, 6, and 8 in a visual form (Arabic digits and visual sets of dots) as well as a tactile form (Braille symbols). For analysis purposes, only numerosities presented in a tactile form were considered. Each participant completed eight runs, with each run lasting 400 seconds. Stimuli were presented in a rapid block design, with each run comprising 72 task blocks lasting between four and six seconds, interspersed with five fixation blocks lasting randomly between four and six seconds, occurring once every 12 task blocks. Additionally, there were two 8-second fixation blocks: one presented before the first task block within the run and one after the last task block, yielding a total of seven fixation blocks. Each task block included four, five, or six stimuli of the same condition (visual Arabic digits, visual sets of dots, or Braille symbols), with variations in visual features for visual stimuli and different spacing on the Braille display for tactile stimuli to avoid adaptation. Each stimulus lasted for one second, including, 300 ms of number presentation followed by 700 ms of centered white circles on a black background. Participants were instructed to evaluate whether the presented number was less or greater than five, indicating their responses by pressing one of two buttons on a response pad. Participants responded with their non-reading hand, with their choice of reading hand determined by their personal preference. Despite all participants being right-handed, 10 of the 25 participants included in the study read with their left hand. fMRI data marking the transition from baseline blocks to tactile Braille blocks were analyzed.

### Motion-based Rejections

Runs were excluded from the analysis if transient head movement exceeded 2 mm of translation and 1° of rotation in any of the three directions. These criteria resulted in excluding 2506 runs out of 13643 runs (18.37%) from the analysis. Information on excluded runs per task are reported in the table below.

**Suppl Table S1:** Number and percentage of runs excluded from the analysis per task based on motion-based rejection criteria.

| Dataset | Task | Total Number of Runs | Number of Excluded Runs | Percentage of Excluded Runs (%) |
|---|---|---|---|---|
| HCP | Gambling | 2170 | 282 | 13% |
| HCP | Relational Processing | 2084 | 367 | 18% |
| HCP | Working Memory | 2173 | 531 | 24% |
| HCP | Social Cognition | 2104 | 374 | 18% |
| HCP | Motor | 2167 | 531 | 25% |
| HCP | Language | 2102 | 355 | 17% |
| UCLA | Spatial Capacity | 122 | 11 | 9% |
| Glasgow | Passive Listening | 217 | 34 | 16% |
| Yale | Taste Perception I | 112 | 5 | 5% |
| Yale | Taste Perception II | 192 | 6 | 3% |
| Jagiellonian Univ. | Reading Braille | 200 | 10 | 5% |
